# Exon definition facilitates reliable control of alternative splicing in the *RON* proto-oncogene

**DOI:** 10.1101/714022

**Authors:** M. Enculescu, S. Braun, S. T. Setty, K. Zarnack, J. König, S. Legewie

## Abstract

Alternative splicing is a key step in eukaryotic gene expression that allows the production of multiple protein isoforms from the same gene. Even though splicing is perturbed in many diseases, we currently lack insights into regulatory mechanisms promoting its precision and efficiency. We analyse high-throughput mutagenesis data obtained for an alternatively spliced exon in the proto-oncogene *RON* and determine the functional units that control this splicing event. Using mathematical modeling of distinct splicing mechanisms, we show that alternative splicing is based in *RON* on a so-called ‘exon definition’ mechanism. Here, the recognition of the adjacent exons by the spliceosome is required for removal of an intron. We use our model to analyze the differences between the exon and intron definition scenarios and find that exon definition is crucial to prevent the accumulation of deleterious, partially spliced retention products during alternative splicing regulation. Furthermore, it modularizes splicing control, as multiple regulatory inputs are integrated into a common net input, irrespective of the location and nature of the corresponding cis-regulatory elements in the pre-mRNA. Our analysis suggests that exon definition promotes robust and reliable splicing outcomes in *RON* splicing.

**SIGNIFICANCE:** During mRNA maturation, pieces of the pre-mRNA (introns) are removed during splicing, and remaining parts (exons) are joined together. In alternative splicing, certain exons are either included or excluded, resulting in different splice products. Inclusion of *RON* alternative exon 11 leads to a functional receptor tyrosine kinase, while skipping results in a constitutively active receptor that promotes epithelial-to-mesenchymal transition and contributes to tumour invasiveness. Intron retention results in to deleterious isoforms that cannot be translated properly. Using kinetic modeling, we investigate the combinatorial regulation of this important splicing decision, and find that the experimental data supports a so-called exon definition mechanism. We show that this mechanism enhances the precision of alternative splicing regulation and prevents the retention of introns in the mature mRNA.

## INTRODUCTION

Eukaryotic gene expression is controlled at multiple levels. One important step in eukaryotic gene regulation is splicing, the removal of intronic sequences from pre-mRNA precursors to yield mature mRNAs. Spliced mRNAs are then exported from the nucleus and translated into protein. In alternative splicing, certain exons are either included or excluded (skipped) to yield distinct mRNA and potentially protein isoforms. Additional isoforms can arise from intron retention, meaning that one or more introns are not removed during splicing. Alternative splicing is thought to be key to transcriptome and proteome complexity in higher eukaryotes and is perturbed in multiple diseases including cancer (1, 2).

Missplicing involving intron retention usually leads to deleterious isoforms containing stop codons or frameshifts that disrupt the open reading frame. Intron retention thus should be in most cases avoided in alternative splicing regulation, since it reduces the amount of functional protein resulting by translation. This is also the case in *RON*, since the intron upstream of alternative exon 11 contains a stop codon, and the downstream intron shifts the open reading frame. High effectiveness of splicing in *RON* requires therefore prevention of intron retention isoforms.

Splicing is catalyzed by a complex molecular machine, the spliceosome, which recognizes splice consensus sequences in nascent pre-mRNAs. The resulting splicing reaction generates mature mRNAs by removing intronic and joining exonic sequences. The catalytic cycle is initiated by recruitment of the U1 and U2 small nuclear ribonucleoprotein (snRNP) subunits to the 5’ splice site and the branch point upstream of the 3’ splice site, respectively. Upon joining of further subunits (U4-U6 snRNPs) and extensive remodeling, a catalytically active higher-order complex is formed. Alternative splicing is commonly regulated by differential recruitment of the U1 and U2 snRNPs. In most cases, such modulation occurs by auxiliary RNA-binding proteins (RBPs) which promote or inhibit U1 or U2 snRNP recruitment by binding to intronic or exonic cis-regulatory elements (1, 3, 4). Spliceosome assembly may occur by two conceptually different mechanisms: In ‘intron definition’, the U1 and U2 snRNPs directly assemble across the intron to form a catalytically competent spliceosome. Alternatively, a cross-exon complex of U1 and U2 snRNPs forms first in a process termed ‘exon definition’ and is then converted into the catalytic cross-intron complex. The simpler intron definition scenario is thought to be the default splicing mechanism for short introns (< 200 bp) that allow for efficient cross-intron spliceosome complex formation (5, 6). Accordingly, intron definition is prevalent in lower organisms such as budding yeast and Drosophila that often display just one or few short introns per gene (6–8). In contrast, exon definition seems to be required for splicing of most mammalian genes, as these typically contain long introns and short exons (6–8). The predominant role of exon definition in mammals is supported by splice site mutation effects on splicing outcomes and by the co-evolution of cis-regulatory elements across exons (6, 8). Furthermore, mathematical models accurately described human splicing kinetics when assuming an exon definition mechanism (9, 10).

Here, we study how intron and exon definition affect the precision and efficiency of alternative splicing regulation. We compare both mechanisms using mathematical modeling, study their functional implications and test the models against comprehensive high-throughput mutagenesis data. As a model system, we use a cancer-relevant human alternatively spliced exon in the RON receptor tyrosine kinase gene, in which the flanking introns are short (87 and 80 nt), implying that both intron and exon definition scenarios are possible (5, 6). Remarkably, our data includes measurements of all arising isoforms, including the ones that exhibit retention of one or both introns. We find that only exon definition quantitatively explains concerted isoform changes upon sequence mutations, and that the changes in the retention isoforms are crucial to distinguish between the two splicing scenarios. We further show that the more complex exon definition pathway provides additional benefits beyond spliceosome assembly across long introns. Our analysis indicates that exon definition greatly simplifies alternative splicing regulation compared to intron definition and efficiently prevents the generation of intron retention products, which are potentially toxic to cells. The presented model provides a framework for the systems-level analysis of complex splice isoform patterns, and offers insights into the mechanistic principles of alternative splicing regulation.

## METHODS

### Extraction and clustering of single point mutation effects on splicing from random mutagenesis screen

We recently established a high-throughput screen of randomly mutated minigenes to decode the cis-regulatory landscape that determines splicing of the alternative exon (AE) 11 in the proto-oncogene *RON* (*MST*1*R*). Experimental details and data have been published in (11). Here we briefly summarize the linear regression model which we used in this previous work to identify single point mutation effects. Furthermore, we describe new data analyses performed in the present paper.

The minigene considered included *RON* exons 10, 11 and 12 together with the intermediate introns. The five splice sites included in the minigene had a normal wildtype strength, as judged by the MaxEnt 5’ and 3’ scores (Fig. 1A and (12)).

**Figure 1:**
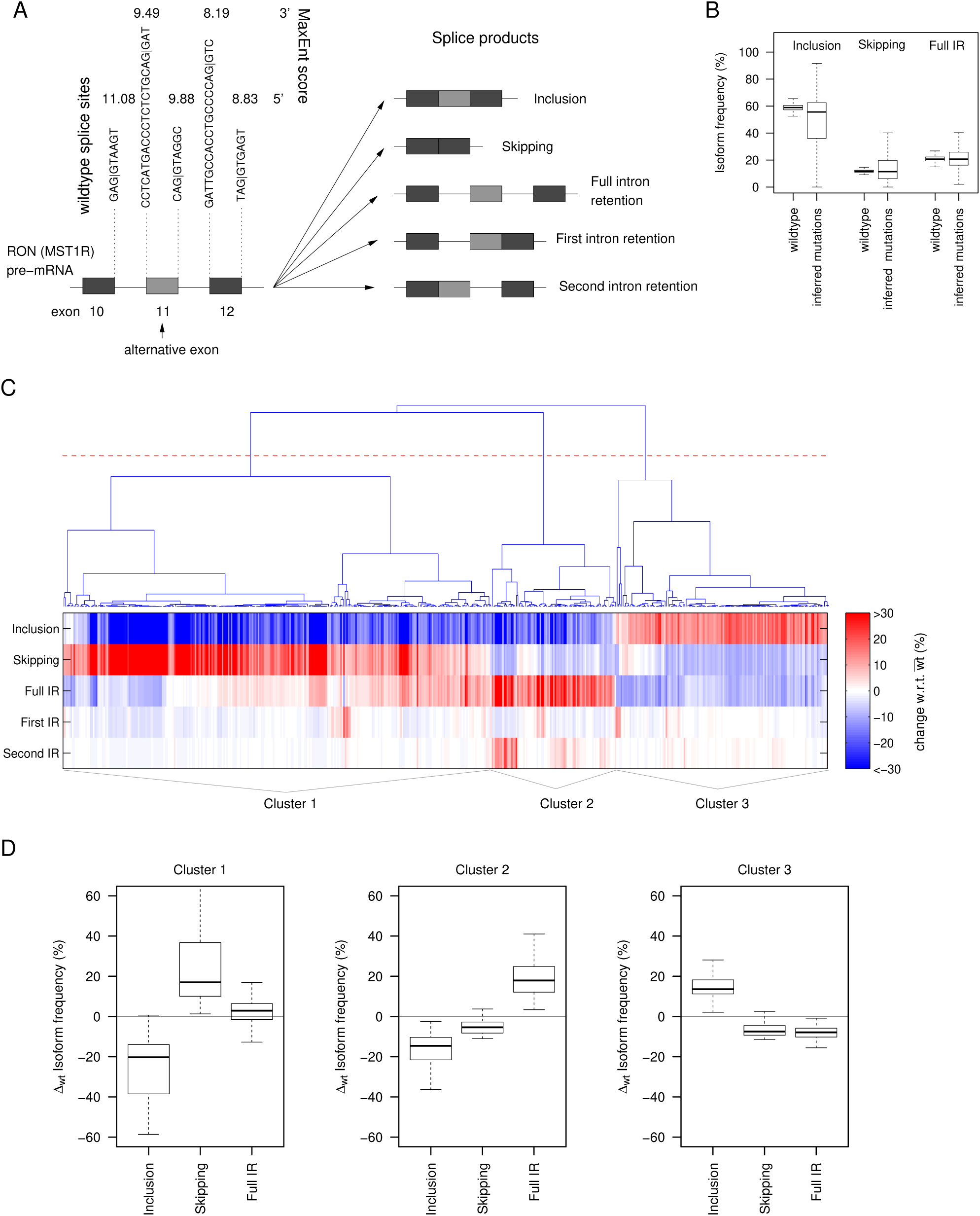
Sequence mutations in a three-exon minigene containing *RON* AE exon 11 induce concerted changes in the distribution of splice isoforms. A The studied three-exon-minigene (704 bp) contains *RON* AE exon 11 and the complete adjacent introns and constitutive exons 10 and 12. The wildtype sequences of the 3’ and 5’ splice sites included in the minigene are shown together with the corresponding splice site scores. Using next-generation sequencing, five different splice products were quantified (as % of all splice products) for wildtype minigenes as well as for single point mutations (see Methods and (11)). B Point mutations induce strong changes in the splice isoform distribution, as visible by the much broader isoform frequency distributions of the mutated minigenes compared to the population of 500 unmutated wildtype (wt) minigenes. Full IR: full intron retention. C Heatmap of splice isoform difference between mutant and wildtype is plotted for 510 point mutations (columns) with a strong effect on the splicing (more than 10% change in at least one isoform frequency with respect to wildtype). Mutations are sorted using hierarchical clustering (Euclidean distance) and three main clusters are defined (using the red line in the dendrogram as a threshold). D Same data as in C, represented as boxplots summarizing the isoform distribution of each cluster for the three main isoforms. Mutations in clusters 1 and 3 induce anti-correlated changes in inclusion and skipping. In cluster 1, these changes are most pronounced in absolute terms, and intron retention is only slightly changed compared to wt. Cluster 3 shows weaker changes and altered intron retention, though in opposite direction. Mutations assigned to cluster 2 decrease both inclusion and skipping and simultaneously increase full intron retention.

Most mutated minigenes in the random mutagenesis screen contained more than one point mutation. As a result, for most mutations only a combined effect on five splice isoforms (AE inclusion, AE skipping, first IR, second IR and full IR, see Fig. 1A) was measured. Linear regression modeling allowed us to infer the effect of single mutations on the splicing outcomes (see Fig. 1B). The key assumption of the model was that mutation effects on isoform ratios (e.g. skipping/inclusion) add up in logarithmic space (or equivalently, that mutations have multiplicative effects on isoform ratios). The predictive power of the regression model for individual point mutation effects on the five splice isoforms was confirmed in two ways: (i) RT-PCR measurements of single mutation effects that were not part of the model training dataset showed an excellent agreement with the model predictions (Fig. 2d in (11)); (ii) In a cross-validation approach, we found that a model fitted to subsets of the original data could accurately predict single mutation minigenes excluded from the training dataset, as soon as the corresponding mutation occurred in at least 5 minigenes (in combination with other mutations) in the training dataset (Fig. 2c in (11)). Based on these findings, we concluded that single point mutation effects on splice isoforms as inferred by regression can be used as a basis for kinetic modeling of splicing decisions. Given the high reproducibility of mutation effects in the initial screen (11), we restricted all analyses described below to the first RNA-seq replicate in HEK293T cells (11).

**Figure 2:**
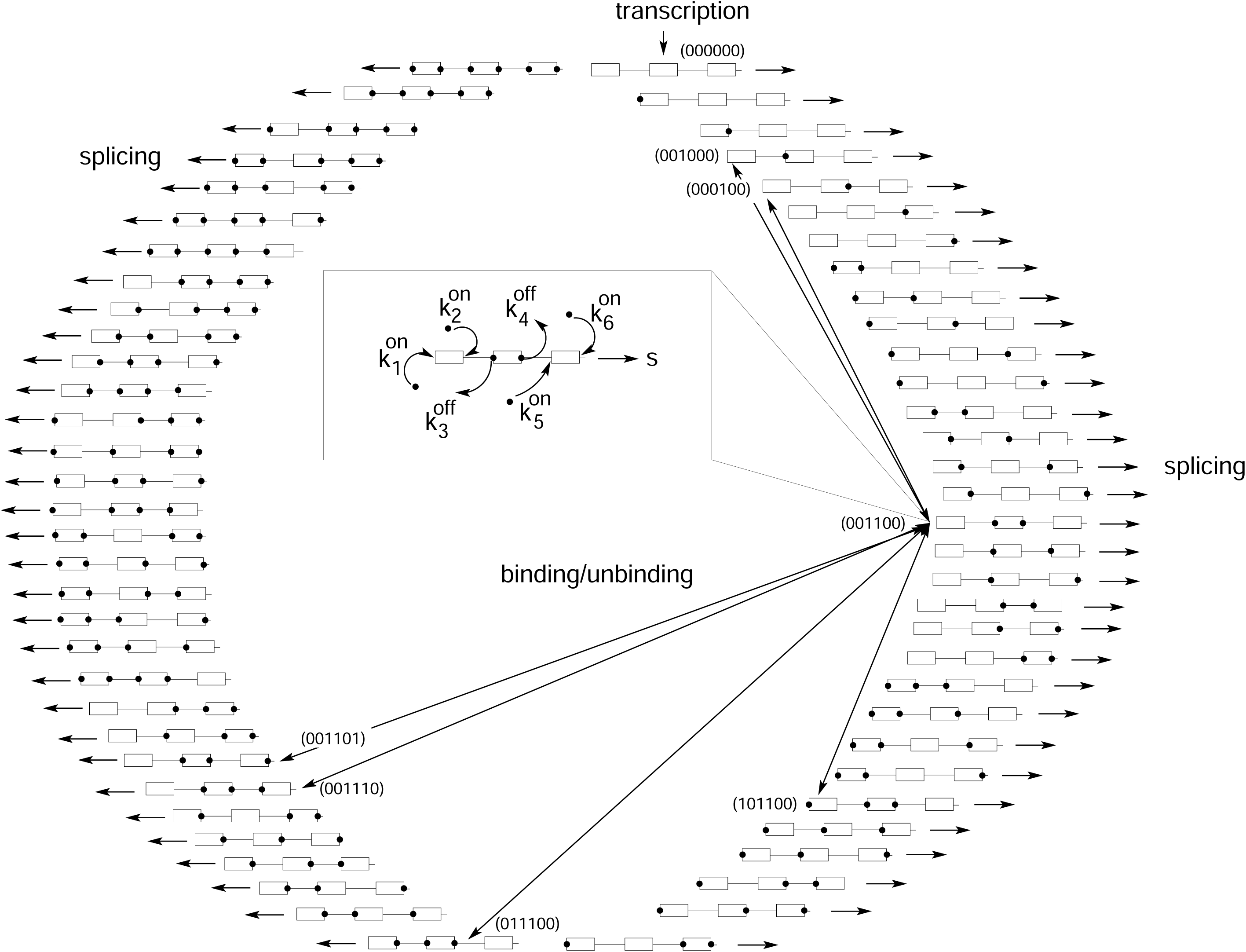
Kinetic model for the derivation of the steady state spliceosome binding probabilities. Each of the 6 splice sites present in the minigene can be bound (1, black circles on pre-mRNA schematic) or unbound (0) by spliceosomal subunits, leading to a total of 2^6^ binding states on the pre-mRNA. The unbound pre-mRNA, denoted by (000000), is produced by transcription, and then spliceosome binding reactions eventually take place. An unbound splice site *i* can be bound at a rate 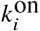 by the spliceosome, and a bound site *i* can unbind again at a rate 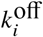. For a better overview, these binding reactions are mostly omitted from the scheme, and only exemplary transitions reflecting the possible reactions for the state (001100), where only both sites of the AE are bound, are shown in the inset and by arrows. Generally, each binding state interacts with 6 different other states in the kinetic reaction scheme. Additionally, splicing of all binding states occurs at rate *s*.

In Fig. 1C, single point mutation effects on the frequencies of five canonical isoforms (AE inclusion, AE skipping, first IR, second IR and full IR) were clustered to identify recurrent patterns in mutation-induced splicing changes. In total, 1942 single mutation effects on splicing were inferred from the random mutagenesis screen (and the splice isoform distributions of all these mutation effects are shown in Fig. 1B). The majority of these mutations induce however only small changes in the isoform distribution compared to wildtype. We therefore selected mutations that induce more than 10% change in at least one of the five canonical isoforms. For each of these 510 mutations, the vector of changes w.r.t. wildtype for the five canonical isoforms was built. These vectors were classified using complete linkage hierarchical clustering and the cosine distance as a similarity measure. Note that for clustering and plotting of Fig. 1C isoform changes of larger than ±30% were bounded to ±30%, while the isoform distributions in Fig. 1D do not include such bounding and therefore show larger maximal changes in isoform frequencies.

### Kinetic model of binding and steady state distribution of binding states

Alternative splicing is commonly regulated by differential recruitment of the U1 and U2 snRNPs to the 5’ and 3’ splice sites. Such differential recruitment affects splicing decision and thus splice isoform distributions (see Fig. 1). To mechanistically understand the emergence of the splice outcome from the binding kinetics, we built an ODE-based mathematical model (sketched in Fig. 2). We described the U1 and U2 binding kinetics and derived the resulting steady state distribution of binding states. Two different splice mechanisms (intron and exon definition, see Fig. 3A) are implemented to connect the binding states to splicing decision and measured splice outcomes.

**Figure 3:**
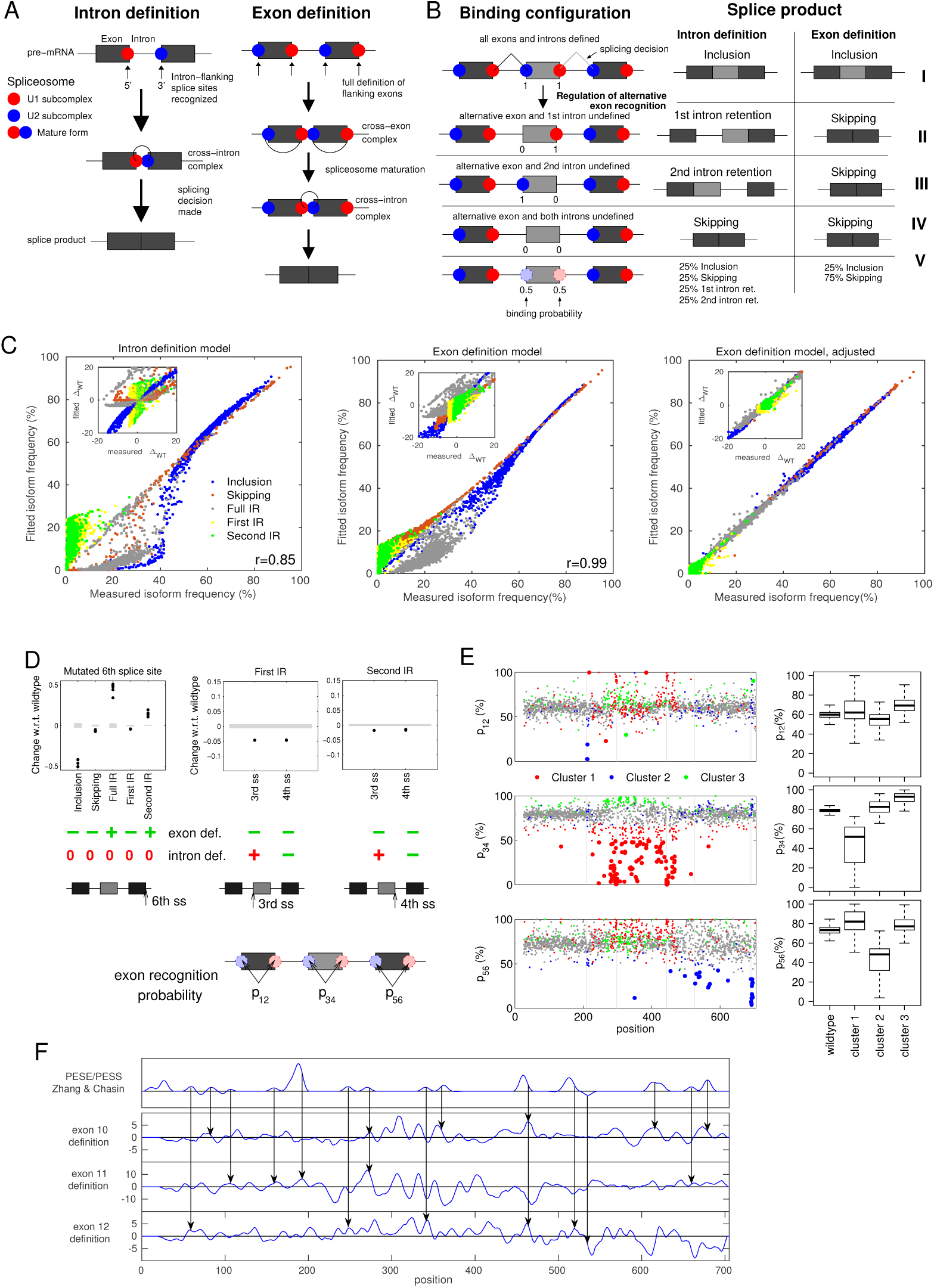
The exon definition model quantitatively explains isoform changes in the mutagenesis screen. A Intron definition model: an intron is spliced out, as soon as its 3’ and 5’ ends splice sites are simultaneously bound by the U1 and U2 snRNP spliceosomal subcomplexes. Exon definition model: Full definition of flanking exons is additionally required for the splicing of an intron, as transitory cross-exon complexes are involved in spliceosome maturation. B Different spliceosome binding configurations (I-IV) of the U1 and U2 subcomplexes may result in distinct splice products in the intron and exon definition models, respectively. In configuration V, binding at the 3’ and 5’ ends of the AE is assumed to take place with 50% probability, giving rise to a equimolar mixture of binding states I-IV. C The exon definition model (middle) shows a better quantitative agreement with the mutagenesis data when compared to the intron definition model (left), as judged by the scatter of model fit against measurements of all five splice isoforms for 1854 point mutations. The performance of the exon definition model is further improved by allowing five global parameters (common to all point mutations) to model under-representation of long intron retention products due to metabolic instability and/or sequencing biases (right). The insets compare model fit (y-axis) vs. data (x-axis) as the difference in splicing outcomes between point mutations and wildtype (zero: no change relative to wt). In terms of directionality of changes, the exon definition model provides a better qualitative match to the measured mutation effects. D Defined point mutations in the 3rd, 4th and 6th splice sites (see bottom sketches) allow for a categorical discrimination between the intron and exon definition mechanisms. Shown are measured splice isoform differences (relative to wt) for minigenes harboring individual point mutations in the indicated splice sites (dots) alongside with the wildtype standard deviation (gray shadows). Directionalities of mutation effects according to the intron and exon definition models – as derived from analytical calculations (Supplementary Material, Section 1) – are indicated below (green for matches with the data, red for contradicting results). E Landscapes and corresponding boxplots showing the exon recognition probabilities (*p*_12_, *p*_34_ and *p*_56_, see scheme) expressed as % recognition for point mutations along the minigene sequence (x-axis) according to the best-fit adjusted exon definition model. The dot color indicates to which cluster a mutation was assigned (see legend and Fig.1; mutations with weak effects, not included in clustering are plotted in gray). Mutations with a recognition probability shift of more than 20% relative to wildtype are highlighted in bold (only the strongest effect being highlighted for each mutation). Mutations in cluster 2 mainly affect the recognition probability of constitutive exons (*p*_12_ or *p*_56_), while mutations in the other two clusters mainly affect alternative exon recognition (*p*_23_), although in different direction and to a different extent. F Landscape of potential ESE/ESS as predicted by the Human Splicing Finder, Version 3.1 (top panel) and lanscapes of the density of strong effects (| z-score| >2) on the recognition probabilities of the three exons (bottom panels). Arrows point to agreements in the peak position and directionality between HSF predictions and our model results.

The minigene analyzed in the paper contains three exons and corresponding introns, and thus altogether 6 splice sites (see main text and Fig. 1). Combinatorial binding of spliceosome subunits (U1 or U2) to these sites results in a total of 2^6^ possible binding states. Fig. 2 shows the considered binding states and examples of possible transitions between them. We can assign each binding state a binary vector with entries 1 at the bound sites and 0 at unbound positions. The completely unbound state (000000) is produced by transcription and can be bound by the spliceosome. Each state can turn into another state by binding of the spliceosome at free splice sites or unbinding of the occupied splice sites. Since we have 6 binding positions, each state can switch to one of 6 other states by binding or unbinding. Additionally, we assume splicing can take place, and that spliced transcripts are not available for further binding/unbinding. A set of linear ordinary differential equations describing the kinetics of the concentration of transcripts in each binding state can be derived. For example, if we denote by *n*_001100_ the concentration of transcripts were only both splice sites of the AE are bound, the temporal evolution of *n*_001100_ will follow:

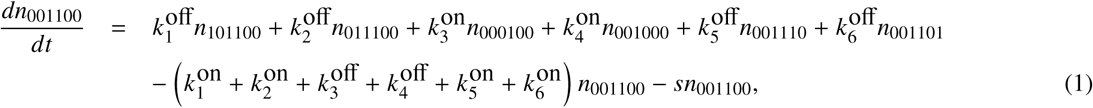

where 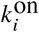 and 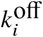 are the binding and unbinding rates at splice site *i* and *s* is the splicing rate (see Fig. 2 and inset). If splicing is slow relative to spliceosome binding and unbinding (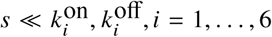, rapid-equilibrium assumption), we can neglect the splicing term in Eq. 1 and can simplify the steady state solution of the ODE system to

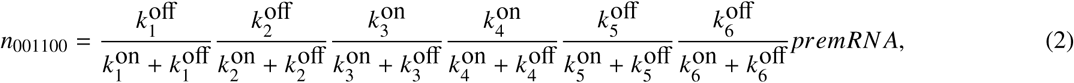

and similar structured terms for all other binding states, where *premRNA* = ∑*n*_*k*_ is the total amount of the unspliced transcripts. The special solution 2 can be verified by substitution in Eq. 1 and neglecting splicing terms. We therefore introduce a steady state recognition probability

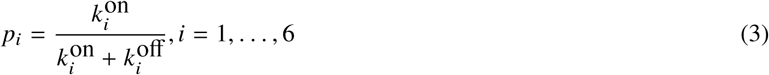

for each splice site *i*, and express the general steady state solution in the form

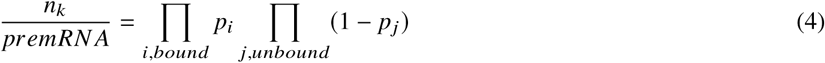

where the first product is taken over all bound sites in state *k* and the second product over all unbound sites is state *k*.

In the above derivation, we have assumed that binding and unbinding of the spliceosome to the pre-mRNA occur post-transcriptionally. However, Eq. 4 is obtained also in the case of co-transcriptional binding/unbinding of the spliceosome, as long as the final splicing reaction takes place post-transcriptionally. By considering the binding of the spliceosome to a particular splice site *i* during the corresponding window of opportunity, we get

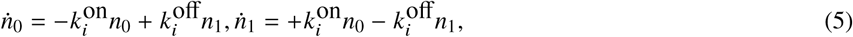

where *n*_0_(*t*) and *n*_1_(*t*) are the number of transcripts with the splice site *i* unbound or bound, respectively, and we neglect possible splicing reactions. If we assume that the time window of opportunity for recognition of this splice site starts at *t* = 0 and ends at *t* = τ_*i*_, we can solve the corresponding initial value problem with *n*_0_(0) = *N*_*i*_, *n*_1_(0) = 0, *N*_*i*_ being the total number of transcripts considered. Since we have *n*_0_(*t*) + *n*_1_(*t*) = *const*. = *N*_*i*_, we get by substituting *n*_1_ = *N*_*i*_ − *n*_0_ in Eqs. 5

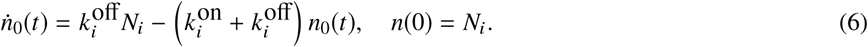

Eqs. 6 can be solved by variation of the constants, leading to the solution

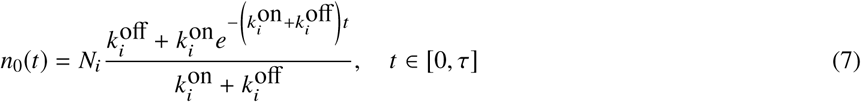

and subsequently

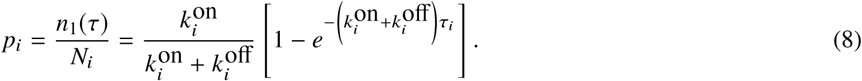

Thus, in the co-transcriptional case, the final probability for a splice site to be bound depends additionally on the length of the corresponding time window of opportunity. Co-transcriptional spliceosome binding/unbinding introduces therefore a correction term in Eqs. 3.

In the general case when splicing is not slow compared to binding/unbinding, Eq. 4 does not hold true. A more complex model considering the competition of splicing reactions and spliceosome binding/unbinding would include a higher number of model parameters, and could not be calibrated properly based on the present data. The approximations made above seem however to be reasonable for the particular splicing decision considered in this paper, since the resulting model is capable to describe our data.

### Splice outcome distribution for the intron definition and exon definition mechanisms

To connect the measured splicing outcome to the binding rates 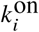 and 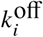, the splice isoform generated from each binding state (i.e., the splicing decision) has to be identified. For six states (Fig. S1A in the Supplementary Material), the splicing outcome is identical for both intron and exon definition mechanisms. For example, when all sites are recognized (state 111111), both introns are spliced out, leading to the inclusion isoform. Similarly, if all except the two flanking splice sites of the alternative exon are recognized, the only possible outcome is skipping. Furthermore, for 32 other binding states no splicing can occur, since no matching 3’ and 5’ splice sites are recognized (Supplementary Material, Fig. S1B). Thus, the only possible splicing outcome of these states is full intron retention. For the remaining 26 partially bound states in Fig. S1C, the splicing outcome depends of the splicing mechanism considered. We have tested two different model variants based on intron and exon recognition mechanisms of splicing (Fig. 3A). For the intron recognition mechanism, we assume that an intron is spliced out, as soon as its 3’ and 5’ sites are recognized. If splicing could occur either across an intron or across a longer sequence containing both introns and the alternative exons (skipping), we assume that the former reaction is much more efficient and neglect the latter. In the exon definition mechanism, an intron is spliced out only if additionally, the adjacent exons are fully defined. The resulting splicing reactions for both mechanisms are indicated in Fig. S1C. The color codes the splice outcome for each binding state. By adding the steady state probabilities *n*_*k*_ */premRNA* for all binding states leading to a certain splice isoform and using Eqs. 4, we get the splice isoform distribution in both models. For the intron definition model, we find the following splice isoform frequencies

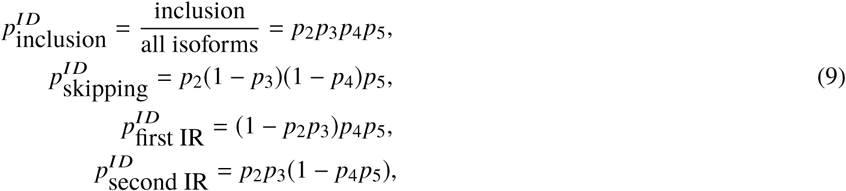

where *p*_*i*_, *i* = 1, …, 6 are the recognition probabilities of the different splice sites defined in Eqs. 3. The splice isoform distribution resulting for the exon definition model reads

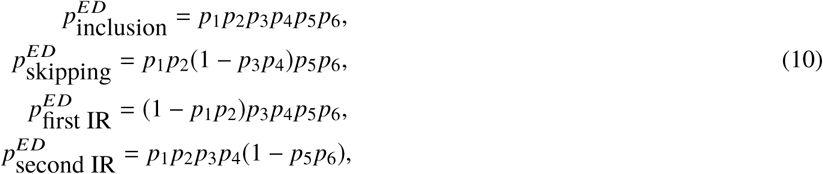

For both models, all frequencies add up to one, implying that the full intron retention probability can be calculated as

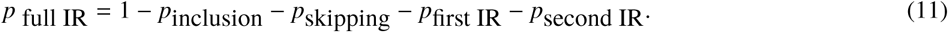

Note that in the exon definition model, *p*_1_ and *p*_2_, *p*_3_ and *p*_4_, as well as *p*_5_ and *p*_6_ appear in all formulae only together. We therefore can introduce

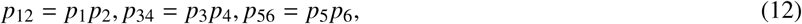

that give the steady state recognition probabilities of the three exons, and reformulate Eqs. 11:

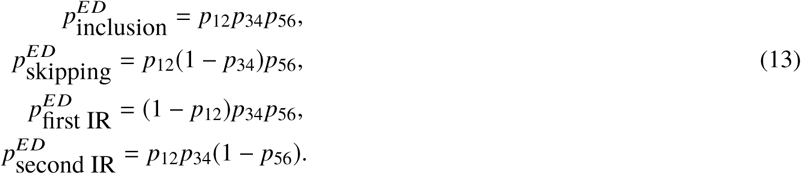

Remarkably, these splice isoform distributions are robust to the precise implementation of the exon definition mechanism: in our model, we assume that U1 and U2 snRNP independently recognize splice sites, and that cross-exon and cross-intron complexes form only later during spliceosome maturation (i.e., they influence only the splicing decision made for a given binding configuration). Alternatively, exon definition may already occur at the level of initial U1 and U2 snRNP binding, because both subunits cooperate across exons during splice site recognition (8, 13). Interestingly, the same isoform distribution formulae as derived above are obtained if we assume highly cooperative binding of U1 and U2 across exons: then, the binding space in Fig. 2 reduces to only 2^3^ states, where each of the three exons are either completely bound or completely unbound, each exon having the overall recognition probability given in Eqs. 12. The binding state distribution are found as above in Eqs. 4, now depending on the recognition probabilities of the exons as a whole given in Eqs. 12. Assuming that splicing can occur between two bound exons, the distribution of the splice outcome in Eqs. 14 is recovered.

## RESULTS

### Mutations in the *RON* minigene induce concerted splice isoform changes

Using high-throughput mutagenesis and next-generation sequencing, we recently quantified the splice products originating from a splicing reporter minigene of the *RON* gene for 1942 single point mutations (11). The three-exon minigene covers *RON* alternative exon 11 (147 nt), the two flanking introns (87 and 80 nt, respectively) as well as constitutive exons 10 and 12 (210 and 166 nt, respectively; Fig. 1A). In HEK293T cells, the major splice product for the unmutated wildtype minigene is exon 11 inclusion (59%), followed by full intron retention (21%), exon 11 skipping (12%), first intron retention (4%) and second intron retention (4%) (Figs. 1A and B). 510 out of the 1942 single point mutations quantified in our study induced significant changes of >10% in the relative abundance of any isoform w.r.t. wildtype (Fig. 1C). Using hierarchical clustering, we sorted these mutations according to their effect on all isoform frequencies, and found three types of splice isoform changes (Fig. 1D): in cluster 1, mutations induced anti-correlated changes in exon 11 inclusion and skipping, with little changes in intron retention isoforms. The remaining mutations additionally affected intron retention, either together with correlated changes in exon 11 inclusion and skipping (cluster 2), or with anti-correlated changes in inclusion and skipping (cluster 3). Taken together, these results indicate that mutation effects in *RON* converge on a small set of splice isoform patterns and may contain information about the underlying regulatory mechanisms.

### Mathematical modeling discriminates intron and exon definition

We turned to mathematical modeling to mechanistically explain mutation-induced changes in splice isoforms. We assumed that mutations influence the splice site recognition by the spliceosome, and modeled the binding of spliceosomes to the pre-mRNA (Figs. 2, 3A and 3B). For simplicity, we only described the initial binding events, i.e., U1 and U2 snRNP binding to the 5’ and 3’ splice sites, respectively. Subsequent spliceosome maturation steps were not modeled explicitly, and it was assumed that splicing decisions are made based on the initial U1 and U2 snRNP recognition patterns (see below). In our model, each U1 or U2 snRNP binding step to one of the six splice sites in the three-exon minigene is characterized by a recognition probability *p*_*i*_. Note that the experimentally measured *RON* minigene lacked the first splice site of exon 10. In the model, we assumed a recognition probability *p*_1_ = 1 to mimick that an exon definition complex forms between mRNA cap-structure and second splice site. We assumed that U1 and U2 snRNP binding is fast compared to the subsequent spliceosome maturation and splicing catalysis. Then, the probabilities *p*_*i*_ are given by 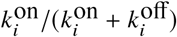, where 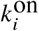 and 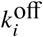 are the binding and unbinding rates of U1 or U2 snRNP to splice site *i* (see Methods). For each pre-mRNA molecule, multiple splice sites can be occupied at a time and depending on the individual recognition probabilities (*p*_*i*_) such simultaneous binding may occur in different combinations. We describe the combinatorial nature of spliceosome binding by combining the individual recognition probabilities *p*_*i*_ into joint probabilities, one for each of the 64 (2^6^) possible U1 and U2 snRNP binding configurations (Fig. 2). For instance, the joint probability of all six splice sites being simultaneously occupied is given by the product *p*_1_ … *p*_6_, and this term changes to (1 − *p*_1_)*p*_2_ … *p*_6_ if the first splice site is not occupied.

In the next step, we assigned a splicing outcome to each of the 64 binding states, and summed up the probabilities over all binding states yielding the same splicing outcome (Methods and Supplementary Material, Fig. S1). We thereby describe the frequency of five splice isoforms as a function of six splice-site recognition parameters (*p*_*i*_). By fitting this model to the measured mutation-induced isoform changes, we infer how mutations affect spliceosomal recognition of splice sites (see below). In two alternative model variants, we implemented splicing decisions based on intron definition and exon definition mechanisms (Figs. 3A and 3B): For the intron definition model, it was assumed that an intron can be spliced out as soon as its flanking 5’ and 3’ splice sites are simultaneously occupied by U1 and U2 snRNPs (Fig. 3A, left). If multiple competing splicing reactions are possible in a binding configuration, we assumed that splicing occurs across the shortest distance (Fig. 3B and S1). The exon definition model involves an additional layer of regulation: before catalytic cross-intron complexes can form, transitory cross-exon U1-U2 snRNP complexes are required to stabilize initial U1/U2 snRNP binding to splice sites (Fig. 3A, right). We implemented this additional requirement for cross-exon complexes by assuming that an intron can only be spliced if all splice sites flanking the adjacent exons are occupied (‘defined’). For example, splicing of the first intron requires full definition of neighboring exons 10 and 11, i.e., simultaneous recognition of splice sites 1-4 in the three-exon minigene (Fig. 3B and S1). Importantly, 26 out of 64 binding configurations generate distinct splicing outcomes in the exon and intron definition models (Fig. S1). Hence, we expect that concerted isoform changes in our mutagenesis dataset (Fig. 1) will discriminate between intron and exon definition mechanisms.

### High-throughput mutagenesis data supports the exon definition model

To investigate whether our mutagenesis data evidence intron and/or exon definition, we separately fitted these model variants to the measured frequencies of five splice isoforms for the wildtype sequence and 1854 single point mutations. We used 1854 out of 1942 single point mutation for fitting, and excluded 88 which resulted in > 5% non-canonical isoforms (see Table S1). During fitting, we assumed that mutations affect the recognition of one or multiple splice sites. In exon definition, the U1 and U2 snRNPs affect splicing only if they are simultaneously bound to both splice sites of an exon. Therefore, splicing outcomes depend only on three effective parameters (*p*_12_, *p*_34_, *p*_56_), each reflecting the recognition probability of the complete exon. Thus, in exon definition there are three free parameters per mutation variant, whereas intron definition results in four independent parameters (see Methods and Supplementary Material).

Despite its lower degree of freedom, the exon definition model provides an overall better fit to the mutagenesis data when compared to the intron definition model (Pearson correlation coefficients = 0.94 vs. 0.85, respectively; Fig. 3C, left and middle panels). The fit quality can be further improved if we additionally allow five global parameters (shared between all mutation variants) to accommodate that long intron retention products may be under-represented in the RNA sequencing library due to metabolic instability (faster degradation of unspliced transcripts (14)) and/or sequencing biases (such as PCR amplification or clustering problems for long fragments on the Illumina flowcell). Taken together, these quantitative results favor exon definition as the predominant mechanism of *RON* splicing. Qualitative arguments based on the algebraic sign of mutation-induced splice isoform changes further disfavor the intron definition model: First, isoform changes in the best-fit intron definition model frequently occur in opposite direction compared to the data, whereas this is not the case for the best-fit exon definition model (Fig. 3C, insets). Second, using analytical calculations, we show that the direction of isoform changes for splice site mutations completely abolishing spliceosome binding is fully consistent with exon definition, but frequently disagrees with intron definition (Fig. 3D, Fig. S2 and Supplementary Material, Section 1). This is particularly evident for mutations of the last splice site (5’ splice site of exon 12) which induce characteristic changes in all splice isoforms (Fig. 3D, left panel). Here, the exon definition model perfectly describes the isoform changes observed in the data, while the intron definition model predicts that the 6^*th*^ splice site does not contribute at all. Notably, the measurement of partial and full intron retention isoforms turns out to be essential for the discrimination of intron and exon definition, since it increases considerably the number of experimentally accessible opposite splice site mutation effects.

Taken together, these results strongly support that *RON* exons 10-12 are spliced via the exon definition mechanism, even though they are flanked by two short introns. Thus, in human cells exon definition may be the preferred and more efficient splicing mechanism, even if the gene structure (intron length) permits the simpler intron definition mode. Currently, it is difficult to generalize this finding to other human genes, since mutagenesis data including quantification of intron retention isoforms are not available in the literature.

Notably, our conclusions concerning *RON* splicing are robust to the precise implementation of the exon definition mechanism: in our model, we assume that U1 and U2 snRNP independently recognize splice sites, and that cross-exon and cross-intron complexes form only later during spliceosome maturation. Alternatively, exon definition may already occur at the level of initial U1 and U2 snRNP binding, because both subunits cooperate across exons during splice site recognition (9, 13). We find that both scenarios lead to the same splice isoform probability equations, implying that our fitting results also apply for strong cross-exon cooperation of U1 and U2 snRNP binding (see Methods).

### Modeling infers spliceosome relocation upon mutations and RBP knockdowns

To further validate the biological plausibility of the exon definition model, we analyzed how the exon recognition probabilities (*p*_12_, *p*_34_, *p*_56_) are perturbed by point mutations in the best-fit model. In line with the intuitive expectation, we find that strong changes in exon recognition require point mutations to be located either within or in close vicinity to the respective exon (Fig. 3E). For the outer constitutive exons, strong mutation effects are mostly confined to the corresponding splice sites, whereas the alternative exon is additionally regulated by a large number of non-splice-site mutations. This reflects the extensive regulation of alternative (but not constitutive) exons by nearby cis-regulatory elements. The recognition probability landscapes also provide plausible explanations for the concerted splice isoform changes we had identified by clustering (Fig. 1): Concerted changes in exon 11 inclusion and skipping (cluster 2) are explained by changes in constitutive exon recognition (*p*_12_ and *p*_56_). On the contrary, any type of anticorrelated change in exon 11 inclusion and skipping (clusters 1, 3) is assigned to perturbed AE recognition (parameter *p*_34_), *p*_34_ being affected with opposing directionality in each of the clusters (decreased *p*_34_ in cluster 1 and increased *p*_34_ in cluster 3).

We have compared the density of strong mutation effects along the minigene with a prediction of potential splice site enhancers (ESE) vs. splice site silencers (ESS) based on (15). To this end, we have calculated the number of mutations with strong effects (| z-score| > 2) on each exon definition probability in a gliding 8-nt window along the minigene, and plotted the difference of the number of enhancing and repressing effects as a running mean over a sliding 8-nt window (splice sites regions were excluded). The potential ESE and ESS based on (15) were analysed with the Human Splicing Finder (HSF), Version 3.1 (16). For the majority of peaks found with HSF, we find a peak of the same directionality in the density of strong effects on the recognition of at least one of the exons (see Fig. 3F). Besides the splice sites, the positions strongest affecting splicing in our dataset are found within the alternative exon and are related to strengthening or weakening of AE recognition due to affected *HNRNPH* binding sites, as we showed in detail in (11). A multitude of further ESE/ESS along the minigene as predicted by HSF based on the minigene sequence is summarized in Fig. S5 and Table S2.

Besides the effects of single point mutations, our model also allows us to quantify the effects of knockdowns of trans-acting RNA-binding proteins that control *RON* splicing. As a proof-of-concept, we fitted the exon definition model to human HEK293 data, in which the levels of the RBP *HNRNPH* were reduced by knockdown and splicing outcomes were subsequently quantified for the population of unmutated minigenes. As shown in Supplementary Material, Fig. S4A, we found that *HNRNPH* mainly controls the recognition of the alternative exon (*p*_34_), with minor effects on the recognition of the outer constitutive exons (*p*_12_ and *p*_56_, respectively). This agrees with our previous finding in human MCF7 cells that while *HNRNPH* binds throughout the minigene sequence, splicing control is mainly executed via a cluster of binding sites within in the alternative exon (11). In Fig. S4B, we confirm for HEK293 cells that *HNRNPH* affects splicing outcomes by binding to the AE.

We have performed knockdown of four additional factors (*PRFP*6, *PUF6*0, *SMU*1 and *SRSF*2) in HEK293 cells. All factors affect recognition of the alternative exon, *PRPF*6, *SMU*1, *SRSF*2 by decreasing and PUF60 by increasing AE recognition (see Fig. S4A). We find that *SMU*1 additionally decreases exon 12 recognition and *PRPF*6 and *SMU*1 enhance exon 10 definition (see Fig. S4A). However, in contrast to *HNRNPH*, the analysis of the interplay between mutations and knockdown effects revealed no clear synergies for these four factors. Thus, the data cannot be used to make any prediction of potential binding sites for these RBPs.

Hence, fitting the exon definition model to perturbation conditions allows us to reconstruct how RBPs affect the splice site recognition by the spliceosome. This constitutes a first step towards reconstruction and mechanistic modeling of combinatorial splicing networks, in which many RBPs jointly control splicing.

### Benefits of splicing regulation by an exon definition mechanism

To explore the biological benefits of exon definition beyond the recognition of exons flanked by long introns, we performed simulations using our splicing models. Interestingly, these simulations revealed that exon definition facilitates alternative splicing control when compared to intron definition. In our models, we simulate alternative splicing regulation by modulating the recognition probability of exon 11 at its 3’ or 5’ splice site. This mimics point mutations or the binding of regulatory RBPs nearby these splice sites. In the intron definition mechanism, splicing outcomes are very distinct, depending on whether *p*_3_ and *p*_4_ are separately or jointly regulated (Fig. 4A, left and Supplementary Material). In contrast, in the exon definition model, splicing outcomes are identical, irrespective of how the recognition of the 3’ and 5’ splice site of the alternative exon is regulated (Fig. 4A, right). Thus, only for exon definition, the alternative exon serves as a regulatory module which integrates inputs on both exon-flanking splice sites into a joint recognition probability *p*_34_ of the alternative exon 11. This modularization simplifies alternative splicing control and ensures that splicing outcomes are robust to the precise location and nature of cis-regulatory elements in the pre-mRNA sequence. Therefore, the seemingly more complex exon definition mechanism in fact simplifies alternative splicing control.

**Figure 4:**
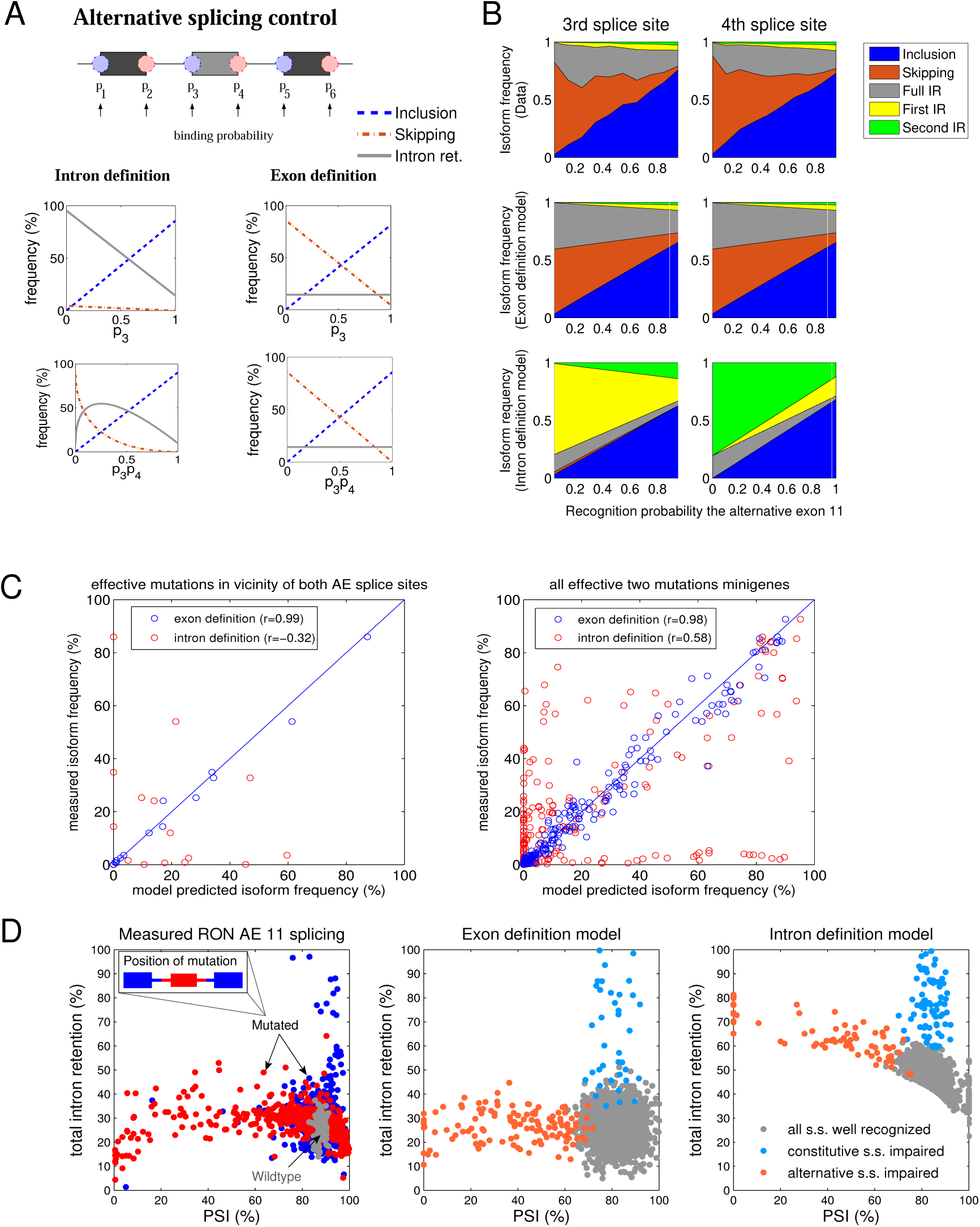
Exon definition allows for modular control of alternative splicing and prevents accumulation of intron retention products. A Simulated splice product frequency of inclusion, skipping and total intron retention (sum of first, second and full IR isoforms) in response to varying recognition of the splice sites flanking the alternative exon. Simulations of the intron and exon definition models are shown in the left and right columns, respectively. The upper plots show the splicing probabilities obtained when only the binding probability of the 3’ splice site (*p*_3_) is varied (similar plots are obtained for variation of *p*_4_, not shown). The bottom plot displays the effects of concerted variation of the two recognition probabilities (*p*_3_ and *p*_4_, x-axis: product *p*_3_*p*_4_). In the intron definition scenario, skipping is possible only if *p*_3_ and *p*_4_ are simultaneously regulated, and is accompanied by enhanced retention. For the exon definition model, separate or joint changes in and/or lead to a switch from skipping to inclusion, without accumulation of retention isoforms. B Mutagenesis data confirms the modular control of AE 11 splicing found in the exon definition scenario. Point mutations at positions in a +/- 30-nt window around the 5’ (left column) or 3’ (right column) splice sites of the AE 11 were selected and sorted according to their effect on the recognition probability of the AE in the best-fit model (adjusted exon definition model). The measured changes in the splice isoform fractions with varying mutation strength (1st row) are similar for both splice sites, and agree with simulations of the exon definition model in which *p*_34_ is systematically varied (2nd row), but disagree with the intron definition model (3rd row). See also main text and Supplementary Material, Section 3. C An exon definition model trained on single point mutation data accurately predicts the abundance of five splice isoforms for combined mutations. Measured isoform frequencies in minigenes containing two mutations are plotted against values predicted by an exon definition model fitted only to single point mutation measurements. The left panel shows three combinations of mutations in a +/- 30 nt window around the 3’ and 5’ splice sites of the AE, and the right panel shows all 45 present combinations of two arbitrary mutations throughout the minigene. Only single mutations that induce strong changes were considered (sum of absolute changes in all five isoforms > 20%). See Supplementary Material, Section 4 for details. D Alternative splicing occurs at low levels of intron retention in the mutagenesis data (left panel). Shown is the sum of all retention products as a function of the PSI-metric (PSI= AE inclusion / (AE inclusion + AE skipping)) which measures alternative splicing of exon 11. The distribution of wildtype minigenes is shown by grey dots and each colored dot represents a single point mutation. Mutations located to the outer constitutive (and adjacent intron halves) are highlighted in blue, whereas the red dots show corresponding mutation effects in or around the AE (see legend). The middle and left panels show 2,000 simulations of the exon and intron definition models, respectively. In these simulations, the splice site recognition parameters (*p*_1_, …, *p*_6_) were randomly perturbed, one randomly chosen parameter being more affected than the others to mimic the experimentally measured PSI and intron retention values (see Supplementary Material, Section 5 for details). Only exon definition allows for alternative splicing at low retention levels.

Exon definition further seems beneficial, as it prevents the accumulation of potentially toxic intron retention products during splicing regulation: using simulations and analytical calculations, we find that the sum of all intron retention products remains constant in the exon definition model if splicing is regulated by the AE recognition parameter *p*_34_ (Fig. 4A, red lines and Supplementary Material, Section 5). In these simulations, the degree of intron retention is solely determined by the recognition probabilities of the outer constitutive exons (*p*_12_ and *p*_56_, see Methods). On the contrary, the intron definition mechanism inevitably leads to a strong accumulation of retention products during alternative splicing regulation, especially if the splice site recognition probabilities *p*_3_ or *p*_4_ are regulated separately (Fig. 4A, left and Supplementary Material, Section 5). In fact, in the intron definition model, pronounced switching from inclusion to skipping isoforms is only possible if *p*_3_ and *p*_4_ are concurrently regulated. However, even in this scenario, intron retention species account for ≥ 50% of the splice products during the splicing transition (Supplementary Material, Section 5).

Using analytical calculations, we confirm that exon modularity and suppression of intron retention also occur for pre-mRNAs containing four exons (see Supplementary Material, Section 6 and Discussion). This suggests that exon definition is generally beneficial from a regulatory point of view.

### Exon definition modularizes splicing regulation

Our simulations show that exon definition modularizes splicing control and prevents the accumulation of intron retention products. To confirm the predicted modularity of alternative splicing regulation, we compared the effects of point mutations located at 3’ and 5’ ends of the alternative exon. Since the exon functions as a module in the exon definition model, we expect that these mutations should have very similar effects on the abundance of splice products. We considered all mutations within a +/- 30 nt window around the 3’ and 5’ splice sites of exon 11. To account for mutation strength, we sorted mutations according to their effect on the AE recognition probability (*p*_34_) in the best-fit model. Then, we plotted the experimentally measured splice isoform abundances as a function of the assigned mutation strength (Fig. 4B). As found in the simulations of the exon definition model, the observed mutation-strength-dependent isoform changes are almost identical for 3’- and 5’-associated mutations. Furthermore, the measured isoform patterns quantitatively agree with simulations of an exon definition model, in which the AE recognition parameter *p*_34_ is systematically varied at otherwise constant recognition probabilities (Fig. 4B, second row). In contrast, corresponding simulations of the intron definition model completely fail to match the data (Fig. 4B, third row). In further support for the exon definition model, we observe highly similar flanking mutation effects not only around the alternative exon but also for the constitutive exon 12 (Fig. S3). In contrast, the intron definition model would predict congruence of mutation effects flanking a common intron, but this behavior is not supported by the experimental data (Fig. 4B and Fig. S3). These observations confirm that exon definition allows exons to function as dominant regulatory modules in alternative splicing control.

To further support that modular exons integrate regulation at the 3’ and 5’ splice site into a joint splicing outcome, we turned to the analysis of combined mutation effects. Therefore, we fitted the adjusted exon definition model to the subset of minigenes harboring only a single mutation, and predicted splicing outcomes of minigenes with a combination of two mutations. The exon definition model accurately captures how two simultaneous mutations in the vicinity of the 3’ and 5’ splice site of exon 11 (each having a strong effect on splicing) jointly affect splicing outcomes (Fig. 4C, left panel). More generally, the exon definition model accurately predicts the combined outcome of any two mutations throughout the minigene (Fig. 4C, right panel). In contrast, a similarly trained intron definition model fails to correctly predict combined mutation effects (Fig. 4C, red dots). Taken together, we find that integration of splice-regulatory input signals in *RON* follows an exon definition scenario which has profound impact on the controllability of alternative splicing.

### Exon definition prevents the accumulation of undesired intron retention products

An important observation in our splicing simulations of the exon definition model is that intron retention products remain constant if splicing is regulated at the alternative exon (by the AE recognition parameter *p*_34_, Fig. 4A). In contrast, intron retention products inevitably accumulate during alternative splicing regulation in the intron definition model (Fig. 4A). To intuitively understand why intron and exon definition differentially affect retention products, consider discrete spliceosome binding configurations (Fig. 3B). If all six splice sites in the three-exon pre-mRNA are occupied by U1 and U2 snRNPs, the splicing outcome is exon inclusion for both mechanisms (Fig. 3B, I). In the next step, alternative splicing can be induced by reducing the recognition of one or both splice sites of the alternative exon. In exon definition, such regulation yields only exon skipping because the middle exon is always incompletely recognized and this impairs splicing of both introns (Fig. 3B, II-IV). In contrast, retention products accumulate in intron definition, as one of the introns remains defined and is therefore spliced (Fig. 3B, II-IV). Our model translates these qualitative arguments into continuous and quantitative predictions of splicing outcomes for five isoforms. For instance, it predicts that in intron definition, retention products strongly accumulate even if the recognition probability of both splice sites is reduced coordinately, e.g., to 50% (*p*_3_ = *p*_4_ = 0.5). This is because combinatorial spliceosome binding to the 3’ and 5’ splice sites results in an equally distributed mixture of four binding configurations, two of which result in retention products (Fig. 3A, V). Hence, exon definition seems superior when compared to intron definition, as it prevents the accumulation of potentially deleterious retention products.

To verify that alternative splicing of *RON* exon 11 is controlled without intron retention, we compared the predictions of our exon definition model to the experimental data. To this end, splicing of the alternative exon was quantified for each point mutation using the PSI metric (percent spliced-in; PSI = AE inclusion/(AE inclusion + AE exclusion)) and then plotted against the corresponding total intron retention level, i.e., the sum of full, first and second intron retention isoforms (Fig. 4D, left panel). In line with the exon definition model, we observed that the majority of point mutations (red and blue dots) induce shifts in alternative splicing (PSI) at almost constant intron retention levels, when compared to wildtype (grey dots). Only a minority (< 2%) of mutations show strong effects on intron retention, but these have in turn only minor effects on the PSI.

These orthogonal changes in either exon inclusion or intron retention could be explained by simulations of the exon definition model, in which we randomly perturbed one of the splice site recognition probabilities, while sampling the others close to their reference value (Fig. 4D, middle panel). The model traces changes in PSI at constant retention levels back to altered splice site recognition of alternative exon 11 (red dots), whereas intron retention enhancement at constant PSI involves reduced recognition of the outer constitutive exons (blue dots, Supplementary Material, Section 5). Consistently, we find in the experimental data that mutations with strong effect on intron retention map to the constitutive exons (Fig. 4D, left; blue dots), whereas mutations affecting PSI are located within or close to the AE (Fig. 4D, left; red dots). Simulations of the intron definition model fail to reconcile the data as the PSI cannot be modulated without accumulation of retention products (Fig. 4D, right panel; Supplementary Material, Section 5). In conclusion, modeling and comprehensive mutagenesis data suggest that alternative splicing by an exon definition mechanism prevents mis-splicing over a wide range of exon inclusion levels. This is likely to be important for *RON* protein function, since all intron retention events in this splicing decision give rise to pre-mature stop codons in the mature mRNA.

## DISCUSSION

Regulatory networks need to produce a certain outcome in a highly precise and controllable fashion. Mathematical models are valuable tools to understand design principles of cellular networks that ensure robustness and precision (17–19). To date, only a handful of mechanistic modeling studies on alternative splicing have been published. These mainly focused on the quantification of mutation effects (9, 11, 20), studied the impact of co-transcriptional splicing (10, 21) and analyzed cell-to-cell variability of the process (22, 23). Here, we approach splicing regulation from a different angle and mechanistically describe how splice site recognition by the spliceosome shapes splicing outcome. We systematically compare intron and exon definition mechanisms, and find that exon definition ensures robust, yet simple regulation of *RON* alternative splicing. Thereby, we gain general insights into the efficiency and controllability of splicing.

Using data-based modeling, we identified exon definition as the mechanism of *RON* exon 11 splicing. The prevalence of exon definition is surprising, given that the flanking introns are very short. Previous work showed that vertebrate exons flanked by short introns can switch to an intron definition mechanism if cross-exon spliceosome complexes are inhibited, e.g., by artificially lengthening the exon (6) or by the lack of exonic splice enhancer elements (24). Our data indicates that a short intron length is not sufficient to switch to intron definition in a natural human exon, and that exon definition is more efficient than intron definition in human cells (25).

Accordingly, we find that exon definition leads to a modularization of splicing regulation. Hence, regulation at one splice site of an exon is transferred to the other splice site, such that exons act as functional units. This has important consequences for the robustness and control of alternative splicing: Our simulations highlight that for pure intron definition, splicing outcomes would be very distinct if splice-regulatory inputs affect the 3’ or the 5’ splice site of the alternative exon (Fig. 4). Furthermore, exon skipping may be difficult to achieve with intron definition, unless both splice sites are coordinately regulated (Fig. 4). Accordingly, a global survey of Drosophila alternative splicing indicated that exons with short flanking introns (likely spliced by an intron definition mechanism) show a strong trend against exon skipping (24). In contrast, in the modular exon definition, inputs at the 3’ and 5’ splice site (or combinations thereof) produce the same splicing outcome. Such signal integration by exon definition is likely to be physiologically relevant, as RBPs frequently control alternative exons by binding nearby only one of the flanking splice sites (26). Arguably, a given RBP can repress or activate splicing depending on its binding position relative to an alternative exon (26). Our model does not exclude such a scenario, but predicts that the net effect of the RBP is integrated in a simple way with signals from other RBPs. In conclusion, exon definition allows for reliable splicing regulation, even though alternative exons are typically influenced by a whole battery of distinct cis-regulatory elements (4, 5).

Exon definition further prevents the accumulation of potentially non-functional intron retention products, and thereby improves the fidelity and efficiency of alternative splicing. In line with our observation that intron retention is difficult to achieve in the exon definition scenario, human splice site mutations most often cause exon skipping and rarely result in intron retention (6). If retention occurs, the mutations are typically located to short introns or affect first or terminal intron of a pre-mRNA, both of which may be more prone to splicing by an intron definition mechanism (6). Intron retention can also serve as a means of active cellular regulation of gene expression. For instance, during granulocyte differentiation, intron retention is enhanced for dozens of genes. Interestingly, this involves a switch from exon to intron definition, as splice factors which favor intron definition complexes are upregulated which promotes the retention of short introns with weak splice sites (27, 28). Our finding that alternative splicing is coupled to intron retention in the intron definition scenario may explain why exon definition is the dominant splicing mecanism in human cells. In contrast, in simpler organisms like S. Cerevisae intron definition may be more prevalent, sinces alternative splicing is largely restricted to the regulated retention of a small number of short introns (28).

In this work, we analyzed a prototypical splicing unit consisting of three exons. Most human genes contain four exons, raising the question whether the described regulatory principles also apply for these more complex scenarios. In the Supplementary Material, we analyze an extended exon definition model containing four exons and show that the inclusion frequency (PSI) of each internal exon is solely determined by its own recognition probably (Supplementary Material, Section 6). Thus, the inclusion of an exon is regulated independently of the neighboring exons. Importantly, this modularity not only involves reliable signal integration on an exon, but also ensures insulation of this exon from other alternative splicing events. In analogy to the three-exon scenario, total intron retention is solely determined by the recognition probabilities of the flanking exons and uncoupled from exon inclusion, i.e., alternative splicing regulation occurs without the accumulation of retention products. Taken together, this suggests that the regulatory benefits of exon definition described in this work continue to hold for more complex pre-mRNAs.

Genome-wide sequencing indicates that 80% of human exons are spliced co-transcriptionally while RNA polymerase is elongating the transcript (7). In this work, we assumed that the final splicing reactions occur with a delay after nascent RNA synthesis. The model holds true if exon recognition occurs co-transcriptionally, as long as the actual splicing takes place after the window of opportunity for the spliceosome binding to the last exon is closed. Only in this scenario, the binding/unbinding dynamics is directly reflected in the splicing outcome, since there is no direct competition between splicing and binding reactions. Notably, this assumption does not exclude that splicing occurs while the transcript is still attached to elongating RNA polymerase. Evidence from the literature supports that splicing of human introns occurs with a delay after nascent RNA synthesis (29) and begins only several kilobases after an intron-exon junction leaves the RNA polymerase complex, with the lag being especially pronounced for alternatively spliced exons (30). Given the median length of human exons and introns (145 and 1964 bp, respectively (1)), the splicing machinery thus likely generates splicing decisions based on sequence stretches containing multiple exons, allowing for neighboring human introns to be spliced concurrently (30, 31) or even in inverse order relative to transcription (26, 30, 31). In this scenario, all six splice sites are available in the pre-mRNA for efficient exon definition. In fact, concurrent splicing of introns further supports exon definition, as this mechanism prevents the accumulation of partially spliced retention products (Fig. 4B). In simpler organisms, splicing is tightly coupled to the exit from the RNA polymerase (30, 32). Thus, the kinetics of splicing may have co-evolved with mechanisms of splice decision making, with slower kinetics being beneficial for exon definition and thus for precise alternative splicing.

## CONCLUSION

Our modeling framework integrates the effect of sequence mutations and knockdowns of trans-acting RBPs on spliceosome recruitment and splicing outcomes (Fig. 3 and Fig. S4). Thus, it constitutes a first step towards a comprehensive network model of splicing which mechanistically describes the integration of multiple splice-regulatory inputs into a net splicing outcome. In fact, we can successfully predict how multiple point mutations jointly control splicing outcomes (Fig. 4C), and the same type of predictions are possible for combined RBP knockdowns and combination of RBP knockdown and sequence mutations. Conceptually, the modeling framework resembles thermodynamic models of transcriptional gene regulation (33–35). However, for the case of splicing, regulation is more complex compared to transcription, as both the regulators (RNA-binding proteins) and the effectors (spliceosomes) show combinatorial binding to multiple sequence elements. Owing to this high level of complexity at multiple levels of splicing regulation, we believe that mechanistic splicing models like the one presented here will be essential to fully disentangle the intricate networks of splicing regulation.

## AUTHOR CONTRIBUTIONS

ME, SL and JK conceived and designed research; ME and SL performed data analysis and modeling; SB and JK performed experiments; STS and KZ analyzed sequencing data; ME and SL wrote the paper with input from JK and KZ;

## ACKNOWLEDGMENTS

The authors would like to thank the members of all participating labs for their support and discussion. We gratefully acknowledge the Institute of Molecular Biology (IMB) Core Facilities for their support, especially the Genomics and the Bioinformatics Core Facilities. We particularly thank Anke Busch from the IMB Core Facility for help with analysis of next-sequencing data; Funding: This work was funded by a joint DFG grant (ZA 881/2-1 to K.Z., KO 4566/4-1 to J.K. and LE 3473/2-1 to S.L.). K.Z. was also supported by the Deutsche Forschungsgemeinschaft (SFB902 B13). S.L. acknowledges support by the German Federal Ministry of Research (BMBF; e:bio junior group program, FKZ: 0316196). The Institute of Molecular Biology (IMB) gGmbH is funded by the Boehringer Ingelheim Foundation;

## REFERENCES

1. Lee, Y., and D. C. Rio, 2015. Mechanisms and regulation of alternative pre-mRNA splicing. Annu. Rev. Biochem. 84:291–323.

2. Xiong, H. Y., B. Alipanahi, L. J. Lee, H. Bretschneider, D. Merico, R. K. Yuen, Y. Hua, S. Gueroussov, H. S. Najafabadi, T. Hughes, Q. Morris, Y. Barash, A. R. Krainer, N. Jojic, S. W. Scherer, B. J. Blencowe, and B. J. Frey, 2015. RNA splicing. The human splicing code reveals new insights into the genetic determinants of disease. Science 347:1254806.

3. Wang, Z., and C. B. Burge, 2008. Splicing regulation: from a parts list of regulatory elements to an integrated splicing code. RNA 14:802–13.

4. Suthandy, F. X. R., S. Ebersberger, L. Huang, A. Busch, M. Bach, H. S. Kang, J. Fallmann, D. Maticzka, R. Backofen, P. F. Stadler, K. Zarnack, M. Sattler, S. Legewie, and J. König, 2018. In vitro iCLIP-based modeling uncovers how the splicing factor U2AF2 relies on regulation by cofactors. Genome Res. 28:699–713.

5. Hertel, K. J., 2008. Combinatorial control of exon recognition. J. Biol. Chem 283:1211–5.

6. Berget, S. M., 1995. Exon recognition in vertebrate splicing. J. Biol. Chem. 270:2411–4.

7. Conti, L. D., M. Baralle, and E. Buratti, 2013. Exon and intron definition in pre-mRNA splicing. Wiley Interdiscip. Rev. RNA 4:49–60.

8. Ke, S., and L. A. Chasin, 2011. Context-depending splicing regulation: exon definition, co-occurring motif pairs and tissue specificity. RNA Biol. 8:384–8.

9. Arias, M. A., A. Lubkin, and L. A. Chasin, 2015. Splicing of designer exons informs a biophysical model for exon definition. RNA 21:213–19.

10. Davis-Turak, J., T. L. Johnson, and A. Hoffmann, 2018. Mathematical modeling identifies potential gene structure determinants of co-transcriptional control of alternative pre-mRNA splicing. Nucleic Acids Res. 46:10598–10607.

11. Braun, S., M. Enculescu, S. T. Setty, M. Cortes-Lopez, B. P. de Almeida, F. X. R. Sutandy, L. Schulz, A. Busch, M. Seiler, S. Ebersberger, N. L. Barbosa-Morais, S. Legewie, J. König, and K. Zarnack, 2018. Decoding a cancer-relevant splicing decision in the RON proto-oncogene using high-throughput mutagenesis. Nat. Commun. 9:3315.

12. Yeo, G., and C. B. Burge, 2004. Maximum entropy modeling of short sequence motifs with applications to RNA splicing signals. Journal of Computational biology 11:377–394.

13. Braun, J. E., L. J. Friedman, J. Gelles, and M. J. Moore, 2018. Synergistic assembly of human pre-spliceosomes across introns and exons. eLife 7:e37751.

14. Lykke-Andersen, S., and T. H. Jensen, 2015. Nonsense-mediated mRNA decay: an intricate machinery that shapes transcriptomes. Nat. Rev. Mol. Cell Biol. 16:665–77.

15. Zhang, X. H.-F., and L. A. Chasin, 2004. Computational definition of sequence motifs governing constitutive exon splicing. Genes & Development 18:1241–1250.

16. Desmet, F.-O., D. Hamroun, M. Lalande, G. Collod-Béroud, M. Claustres, and C. Béroud, 2009. Human Splicing Finder: an online bioinformatics tool to predict splicing signals. Nucleic Acids Research 37:e67.

17. Blüthgen, N., and S. Legewie, 2013. Robustness of signal transduction pathways. Cell. Mol. Life Sci. 70:2259–69.

18. Kamenz, J., T. Mihaljev, A. Kubis, S. Legewie, and S. Hauf, 2015. Robust ordering of anaphase events by adaptive thresholds and competing degradation pathways. Mol. Cell 60:446–59.

19. Enculescu, M., C. Metzendorf, R. Sparla, M. Hahnel, J. Bode, M. U. Muckenthaler, and S. Legewie, 2017. Modelling systemic iron regulation during dietary iron overload and acute inflammation: Role of hepcidin-independent mechanisms. PLoS Comput. Biol. 13:e1005322.

20. Baeza-Centurion, P., B. M. nana, J. M. Schmiedel, and B. L. J. Valcárcel, 2019. Combinatorial genetics reveals a scaling law for the effects of mutations on splicing. Cell 176:549–563.

21. Davis-Turak, J. C., K. Allison, M. N. Shokhirev, P. Ponomarenko, L. S. Tsimring, C. K. Glass, T. L. Johnson, and A. Hoffmann, 2015. Considering the kinetics of mRNA synthesis in the analysis of the genome and epigenome reveals determinants of co-transcriptional splicing. Nucleic Acids Res. 43:699–707.

22. Waks, Z., A. M. Klein, and P. A. Silver, 2011. Cell-to-cell variability of alternative RNA splicing. Mol. Syst. Biol. 7:506.

23. Schmidt, U., E. Basyuk, M. C. Robert, M. Yoshida, J. P. Villemin, D. Auboeuf, S. Aitken, and E. Bertrand, 2011. Real-time imaging of cotranscriptional splicing reveals a kinetic model that reduces noise: implications for alternative splicing regulation. J. Cell Biol. 193:819–29.

24. Fox-Walsh, K. L., Y. Dou, B. J. Lam, S. P. Hung, P. F. Baldi, and K. J. Hertel, 2005. The architecture of pre-mRNAs affects mechanisms of splice-site pairing. Proc. Natl. Acad. Sci. USA 102:16176–81.

25. To further exclude splicing by an intron definition mechanism, we also considered mixed intron and exon definition models, in which only a subset of the three exons acts as a functional unit, whereas the remainder affects splicing already when partially defined. Interestingly, only the full exon definition model was consistent with the mutagenesis data, further suggesting that none of the two RON introns is spliced by a direct cross-intron spliceosome complex (data not shown).

26. Witten, J. T., and J. Ule, 2011. Understanding splicing regulation through RNA splicing maps. Trends Genet. 27:89–97.

27. Wong, J. J., W. Ritchie, O. A. Ebner, M. Selbach, J. W. Wong, Y. Huang, D. Gao, N. Pinello, M. Gonzalez, K. Baidya, A. Thoeng, T. L. Khoo, C. G. Bailey, J. Holst, and J. E. Rasko, 2013. Orchestrated intron retention regulates normal granulocyte differentiation. Cell 154:583–95.

28. A. G. Jacob, C. W. J. S., 2017. Intron retention as a component of regulated gene expression programs. Hum. Genet. 136:1043–1057.

29. Nojima, T., T. Gomes, M. Carmo-Fonseca, and N. J. Proudfoot, 2016. Mammalian NET-seq analysis defines nascent RNA profiles and associated RNA processing genome-wide. Nat. Protoc. 11:413–28.

30. Drexler, H. L., K. Choquet, and L. S. Churchman, 2019. Human co-transcriptional splicing kinetics and coordination revealed by direct nascent RNA sequencing. https://www.biorxiv.org/content/10.1101/611020v2.

31. Kim, S. W., A. J. Taggart, C. Heintzelman, K. J. Cygan, C. G. Hull, J. Wang, B. Shrestha, and W. G. Fairbrother, 2017. Widespread intra-dependencies in the removal of introns from human transcripts. Nucleic Acids Res. 45:9503–9513.

32. Oesterreich, F. C., L. Herzel, K. Straube, K. Hujer, J. Howard, and K. M. Neugebauer, 2016. Splicing of nascent RNA Coincides with intron exit from RNA polymerase II. Cell 165:372–381.

33. Bintu, L., N. E. Buchler, H. G. Garcia, U. Gerland, T. Hwa, J. Kondev, and R. Phillips, 2005. Transcriptional regulation by the numbers: models. Curr. Opin. Genet. Dev. 15:116–24.

34. Casanovas, G., A. Baneji, M. U. M. F. d’Alessio, and S. Legewie, 2014. A multi-scale model of hepcidin promoter regulation reveals factors controlling systemic iron homeostasis. PLoS Comput. Biol. 10:e1003421.

35. Schulthess, P., A. Löffler, S. Vetter, L. Kreft, M. Schwarz, A. Braeuning, and N. Blüthgen, 2015. Signal integration by the CYP1A1 promoter – a quantitative study. Nucleic Acids Res. 43:5318–30.

