## Supplementary Text for "Exon definition facilitates reliable control of alternative splicing in the *RON* proto-oncogene"

**Authors:** Mihaela Enculescu[1], Simon Braun[1], Samarth Thonta-Setty[2], Kathi Zarnack[2], Julian König[1], Stefan Legewie[1]

**Affiliations:**

[1]Institute of Molecular Biology, Ackermannweg 4, 55128 Mainz, Germany.

[2]Buchmann Institute for Molecular Life Sciences, Goethe University Frankfurt, Max-von-Laue-Str. 15, 60438 Frankfurt a.M., Germany

### Content

**1 Splice site mutation effects ……………………………………………………………2**

***1.1 Effects of splice site mutations in the intron and exon definition models***

***1.2 Directionality of splice site mutations effects in the intron and exon definition models***

***1.3 Adjusted exon definition model with variable isoform-specific degradation rates***

***1.4 Effect of variable degradation rates on the directionality of changes upon splice site mutations***

### 1 Splice site mutation effects

Mutations in splice site consensus sequences impair the recognition of the splice sites by the spliceosome. Thus, they specifically lower one of the binding probabilities $p_i, i=1,\ldots,6$, suggesting that measurements of splicing outcomes for these mutations are valuable to test our models.

#### *1.1 Effects of splice site mutations in the intron and exon definition models*

Interestingly, the binding probabilities modulate the splicing outcome differently in the intron and exon definition models, as found by comparing Eqs. (5) and (6): For example, the recognition probability of the last splice site $p_6$ has an effect on all splice isoforms in the exon definition model, but does not affect splicing in the intron definition model.

Fig. S2 compares the experimentally measured effects of splice site mutations in our mutagenesis screen to the effects predicted by the intron and exon definition models (and Fig. 3D shows a subset of this analysis). Plotted are changes in the frequency of each isoform with respect to wildtype for splice site mutations at sites (2-6). Mutations affecting the first splice site were not present in our minigene library and were therefore excluded from this analysis. The experimentally measured changes are plotted as a median of the isoform changes among all mutations affecting the first two or last two positions of an intron (GT and AG splice site consensus sequences, respectively). For both models, qualitative simulations were performed (middle panels) and the parameters were set to $p_1=1$ and $p_i=0.9, i=2,\ldots,6$ in wildtype. The effect of a splice site mutation is qualitatively simulated by reducing the corresponding probability from 0.9 to 0.1. The rightmost panel in Fig. S2 shows the quantitative fit of the final adjusted exon definition model to the data (Fig. 3C, right). In general, the exon definition model (and in particular its best-fit version) agrees much better with the data than the intron definition model.

In fact, using analytical calculations (Section 2.4), one can show that the directionality of isoform changes upon splice site mutations, shown in the middle panels of Fig. S2, holds true for arbitrary values of the recognition parameters $p_i, i=2,\ldots,6$. Hence, the qualitative effects of splice site mutation on splice isoforms provide a unique footprint that distinguishes intron and exon definition models. The data (left panel) in Fig. S2 supports exon definition in contrast to intron definition in several points:

1. Mutations of the 6th splice site affect all isoforms (dark blue bars).
2. Mutations of the 3rd and 4th splice sites (red bars) induce comparable changes for each of the isoforms, supporting that the AE acts as module, as predicted by the exon definition model. In contrast, the intron definition model predicts the recognition parameters $p_3$ and $p_4$ induce opposite changes in first and second intron retention.
3. Mutation effects on inclusion and skipping correspond also quantitatively to the exon definition model, while the intron definition model leads to a much smaller amplitude of changes in skipping for all splice sites mutations.
4. Directionality of changes agrees to the exon definition model for 23 out of 25 effects, with exception of the changes in full intron retention at mutated AE sites (3rd and 4th splice sites, red bars). As shown below, a full match between exon definition model and data for all 25 effects can be achieved by additional model extensions (Sections 2.5. and 2.6).

Our data therefore strongly indicate that an exon definition mechanism controls RON AE 11 splicing.

***1.2 Directionality of splice site mutations effects in the intron and exon definition models***

Using the models in Eqs. (5) and (9) as well as Eq. (7) we can uniquely determine the directionality of changes in the splice isoform frequencies upon splice site mutations. If we denote by $\nabla p_{isoform}^{ID}$ the derivatives of the isoform frequencies in the intron definition model w.r.t. the model parameters $(p_{2,} p_{3,} p_{4,} p_5)$ , we get from Eqs. (5) and (7)

$$\begin{gathered}
\nabla p_{inclusion}^{ID} = [p_3 p_4 p_{5,} p_2 p_4 p_{5,} p_2 p_3 p_{5,} p_2 p_3 p_4], \\
\nabla p_{skipping}^{ID} = [(1-p_3)(1-p_4) p_{5,} - p_2 (1-p_4) p_{5,} - p_2 (1-p_3) p_{5,} p_2 (1-p_3)(1-p_4)], \\
\nabla p_{firstIR}^{ID} = [-p_3 p_4 p_{5,} - p_2 p_4 p_{5,} (1-p_2 p_3) p_{5,} (1-p_2 p_3) p_4], \\
\nabla p_{secondIR}^{ID} = [p_3 (1-p_4 p_5), p_2 (1-p_4 p_5), -p_2 p_3 p_{5,} - p_2 p_3 p_4], \\
\nabla p_{fullIR}^{ID} = [-p_3 (1-p_5) - p_5 (1-p_4), -p_2 (1-p_5), -p_5 (1-p_2), -p_2 (1-p_3) - p_4 (1-p_2)].
\end{gathered} \qquad (10)$$

Since the splice site recognition probabilities are bounded between 0 and 1, the sign of the all derivatives with respect to the model parameters is fixed, e.g.

$$\frac{dp_{first}^{ID}}{dp_3} = -p_2 p_4 p_5 < 0, for\, all\, 0 < p_i < 1, i = 2,4,5. \qquad (11)$$

Similarly, for the exon definition model we can denote by $\nabla p_{isoform}^{ED}$the derivatives of the single isoform frequencies w.r.t. to the exon definition probabilities $(p_{12}, p_{34}, p_{56})$. Using Eqs. (7) and (9) we find

$$\begin{gathered}
\nabla p_{inclusion}^{ED} = [p_{34} p_{56}, p_{12} p_{56}, p_{12} p_{34}], \\
\nabla p_{skipping}^{ED} = [(1-p_{34}) p_{56}, -p_{12} p_{56}, p_{12} (1-p_{34})], \\
\nabla p_{firstIR}^{ED} = [-p_{34} p_{56}, (1-p_{12}) p_{56}, (1-p_{12}) p_{34}], \\
\nabla p_{secondIR}^{ED} = [p_{34} (1-p_{56}), p_{12} (1-p_{56}), -p_{12} p_{34}], \\
\nabla p_{fullIR}^{ED} = -[p_{34} (1-p_{56}) + p_{56} (1-p_{34}), p_{12} (1-p_{56}) + p_{56} (1-p_{12}), p_{12} (1-p_{34}) + p_{34} (1-p_{12})].
\end{gathered} \qquad (12)$$

Also for the exon definition model, all derivatives have fixed sign, e.g.

$$\frac{dp_{first}^{ED}}{dp_{34}} = (1-p_{12}) p_{56} > 0, for\, all\, 0 < p_i < 1, i = 12,56. \qquad (13)$$

By comparing the signs of the single terms in Eqs. (10) and (12) we can find differences in the directionality of predicted changes for splice site mutations in the two models, and thereby discriminate both models based on experimental data (see Section 2.3). For instance, a mutation in the 3rd splice site decreases the recognition probabilities $p_3$ and $p_{34}$ in the intron and exon definition models, respectively$p_{34}$. By comparing Eqs. (11) and (13) we find that for such a mutation, first intron retention is predicted to increase in the intron definition model, but to decrease in the exon definition model. The latter is observed experimentally (Fig. S2), which supports the exon definition model.

The analytically calculated directionalities of isoform changes in response to splice site mutations for intron and exon definition mechanisms are indicated by '+' and '-' signs in the bottom of Fig. 3D.

#### 1.3 Adjusted exon definition model with isoform-specific degradation rates

The exon definition model reproduces almost completely the directionality of splice isoform changes upon splice site mutations (see Fig. S2, first and third columns). However, it fails to reproduce the measured changes in full IR, overestimates mutation effects on first and second intron retention, and slightly underestimates the effects on skipping (Fig. S2). As detailed below, these quantitative differences between model and data can be counterbalanced by assuming that different splice products degrade at different rates (see Fig. S2, right panel).

In the adjusted exon definition model, the measured splice isoforms are assumed to be produced at a rate proportional to the individual splice isoform probabilities derived in Eqs. (9), but are degraded at individual rates. This leads to the following kinetic model

$$\frac{d}{dt} inclusion = s\, p^{ED}_{inclusion}\, premRNA - d_1\, inclusion,$$
$$\frac{d}{dt} skipping = s\, p^{ED}_{skipping}\, premRNA - d_2\, skipping,$$
$$\frac{d}{dt} fullIRtot = \frac{d}{dt} premRNA + \frac{d}{dt} fullIR = trans - (s + d_0)\, premRNA$$
$$+ s\left(1 - p^{ED}_{inclusion} - p^{ED}_{skipping} - p^{ED}_{firstIR} - p^{ED}_{secondIR}\right) premRNA - d_3 (fullIRtot - premRNA), \quad (14)$$
$$\frac{d}{dt} first = s\, p^{ED}_{firstIR}\, premRNA - d_4\, firstIR,$$
$$\frac{d}{dt} secondIR = s\, p^{ED}_{secondIR}\, premRNA - d_5\, secondIR.$$

Here, *fullIRtot* is the sum of the two unspliced species, nascent pre-mRNA and mature unspliced mRNA (*fullIR*), which cannot be distinguished experimentally. The nascent pre-mRNA is produced by transcription at a rate *trans*, spliced at total rate $s$ and degraded at rate $d_0$.

The steady state found by inserting Eqs. (9) in (14) is given by

$$premRNA = \frac{trans}{s + d_0},\ inclusion = premRNA \frac{s}{d_1} p_{12} p_{34} p_{56},$$
$$skipping = premRNA \frac{s}{d_2} p_{12} (1 - p_{34}) p_{56},$$
$$fullIRtot = premRNA \left[1 + \frac{s}{d_3}\left(1 - p_{12} p_{34} - p_{12} p_{56} - p_{34} p_{56} + 2 p_{12} p_{34} p_{56}\right)\right], \quad (15)$$
$$firstIR = premRNA \frac{s}{d_4} (1 - p_{12}) p_{34} p_{56},\ secondIR = premRNA \frac{s}{d_5} p_{12} p_{34} (1 - p_{56}).$$

In these steady state equations, synthesis and degradation terms always occur as ratios (e.g., $s$ and $d_1$for inclusion). This implies that the model also accommodates that each isoform may be generated with its own splicing rate$s_i$from the pre-mRNA precursor, as this would ultimately lead to an isoform-specific value for the ratio ($s_i / d_i$), much like an isoform-specific degradation rate. Likewise, length biases in RNA sequencing – which may disfavor the detection of long isoforms such as full intron retention – are expected to scale this ratio and are therefore effectively considered in the model. Hence, the exon definition model with isoform-specific degradation takes into account various technical and biological effects leading to isoform biases and thereby allows for a more realistic description of splicing decisions.

Finally, we can connect the steady state to the measured isoform *frequencies* by normalizing Eqs. (15) with the total number of measured RNA transcripts:

$$total = inclusion + skipping + fullIRtot + firstIR + secondIR, \quad (16)$$

and find

$$p_{inclusion} = \frac{s\, p_{12} p_{34} p_{56}}{d_1 R}, \; p_{skipping} = \frac{s\, p_{12}(1-p_{34}) p_{56}}{d_2 R},$$
$$p_{fullIR} = \frac{\left[d_3 + s(1 - p_{12}p_{34} - p_{12}p_{56} - p_{34}p_{56} + 2p_{12}p_{34}p_{56})\right]}{d_3 R}, \quad (17)$$
$$p_{firstIR} = \frac{s(1-p_{12}) p_{34} p_{56}}{d_4 R}, \; p_{secondIR} = \frac{s\, p_{12} p_{34}(1-p_{56})}{d_5 R}.$$

Here, the factor R in Eqs. (17) equals the total quantified RNA to pre-mRNA ratio

$$R = \frac{total}{premRNA} = \frac{mRNA + premRNA}{premRNA} \quad (18)$$

and is equal to $(1+s/d)$if all degradation rates are the same ($d_i = d, i = 1,\ldots,5$). In this case, the simple model given in Eqs. (9) is recovered. For variable degradation rates, $R$ reads

$$R = 1 + \frac{s\, p_{12} p_{34} p_{56}}{d_1} + \frac{s\, p_{12}(1-p_{34}) p_{56}}{d_2} + \frac{s(1 - p_{12}p_{34} - p_{12}p_{56} - p_{34}p_{56} + 2p_{12}p_{34}p_{56})}{d_3}$$
$$+ \frac{s(1-p_{12}) p_{34} p_{56}}{d_4} + \frac{s\, p_{12} p_{34}(1-p_{56})}{d_5}. \quad (19)$$

#### *1.4 Effect of variable degradation rates on the directionality of changes upon splice site mutations*

The inclusion of isoform-specific degradation rates leads to a better quantitative model fit of the exon definition to the mutagenesis data (Fig. 3C). Furthermore, on a qualitative level, this model extension can fix the disagreement between the exon definition model and the experimental data with respect to the directionality of splice site mutation effects on the full IR isoform (Fig. S2). In fact, this will be proven below by showing that the consideration of variable degradation rates does not switch the directionality of splice isoform changes compared to a model with common degradation rates, except for the isoform full intron retention.

The proof of this claim can be done by looking at the structure of the adjusted model derived in Eqs. (17). When computing the derivative of the splice isoforms inclusion, skipping, first or second IR with respect to a certain exon definition probability$p_i$we find either

$$\frac{\partial p_{isoform}}{\partial p_i} = \frac{p_{isoform}}{p_i R}\left(R - p_i \frac{\partial R}{\partial p_i}\right), \quad (20)$$

if the particular isoform $p_{isoform}$ increases with $p_i$($p_{isoform} \sim p_i$) in the basic exon definition model derived in Eqs. (9), or

$$\frac{\partial p_{isoform}}{\partial p_i}=-\frac{p_{isoform}}{(1-p_i)R}\left(R+(1-p_i)\frac{\partial R}{\partial p_i}\right), \quad (21)$$

if the particular isoform decreases with $p_i$($p_{isoform} \sim (1-p_i)$) in the unadjusted model of Eqs. (9). For example, for the skipping isoform $p_{skipping}=\frac{p_{12}(1-p_{34})p_{56}}{d_2 R}$ we get:

$$\frac{\partial p_{skipping}}{\partial p_{12}}=\frac{p_{skipping}}{p_{12}R}\left(R-p_{12}\frac{\partial R}{\partial p_{12}}\right),$$
$$\frac{\partial p_{skipping}}{\partial p_{34}}=-\frac{p_{skipping}}{(1-p_{34})R}\left(R+(1-p_{34})\frac{\partial R}{\partial p_{34}}\right), \quad (22)$$
$$\frac{\partial p_{skipping}}{\partial p_{56}}=\frac{p_{skipping}}{p_{56}R}\left(R-p_{56}\frac{\partial R}{\partial p_{56}}\right).$$

The normalization factor $R$ depends on the exon definition probabilities $p_i, i=12,34,56$ like a first order polynomial, as given in Eq. (19). We thus have

$$R-p_{12}\frac{\partial R}{\partial p_{12}}=1+\frac{s}{d_3}(1-p_{34}p_{56})+\frac{p_{34}p_{56}}{d_4}>1,$$
$$R-p_{34}\frac{\partial R}{\partial p_{34}}=1+\frac{s}{d_3}(1-p_{12}p_{56})+\frac{p_{12}p_{56}}{d_2}>1, \quad (23)$$
$$R-p_{56}\frac{\partial R}{\partial p_{56}}=1+\frac{s}{d_3}(1-p_{12}p_{34})+\frac{p_{12}p_{34}}{d_5}>1.$$

Furthermore, from Eq. (19) we can calculate

$$R+(1-p_{12})\frac{\partial R}{\partial p_{12}}=1+\frac{s\,p_{34}p_{56}}{d_1}+\frac{s(1-p_{34})p_{56}}{d_2}+\frac{s(1-p_{34})(1-p_{56})}{d_3}+\frac{s\,p_{34}(1-p_{56})}{d_5}>1,$$
$$R+(1-p_{34})\frac{\partial R}{\partial p_{34}}=1+\frac{s\,p_{12}p_{56}}{d_1}+\frac{s(1-p_{12})(1-p_{56})}{d_3}+\frac{s(1-p_{12})p_{56}}{d_4}+\frac{s\,p_{12}(1-p_{56})}{d_5}>1, \quad (24)$$
$$R+(1-p_{56})\frac{\partial R}{\partial p_{56}}=1+\frac{s\,p_{12}p_{34}}{d_1}+\frac{s\,p_{12}(1-p_{34})}{d_2}+\frac{s(1-p_{12})(1-p_{34})}{d_3}+\frac{s(1-p_{12})p_{34}}{d_4}>1.$$

Note that we have used the fact that all probabilities $p_i, i=12,34,56$ are positive numbers smaller than 1. By combining Eqs. (20) and (23) we thus find

$$\frac{\partial p_{isoform}}{\partial p_i}=\frac{p_{isoform}}{p_i R}\left(R-p_i\frac{\partial R}{\partial p_i}\right)>0, \quad (25)$$

for all isoforms $p_{isoform}$ that increase with $p_i$ in the basic exon definition model derived in Eqs. (9). By combining Eqs. (21) and (24) we further find

$$\frac{\partial p_{isoform}}{\partial p_i}=-\frac{p_{isoform}}{(1-p_i)R}\left(R+(1-p_i)\frac{\partial R}{\partial p_i}\right)<0, \quad (26)$$

for all isoforms $p_{isoform}$ that decrease with $p_i$ in the unadjusted exon definition model derived in Eqs.

(9). Therefore, the introduction of individual degradation rates cannot induce a switch in the directionality of changes with respect to the exon definition probabilities, but can adjust only the amplitude of changes for inclusion, skipping, first and second intron retention.

In contrast, numeric simulations show that variable degradation rates can induce a switch of the directionality in the changes of full intron retention, and thus correct the difference between the basic exon definition model and data (see Fig. S2).

### 2 Quantitative estimation of mutations effects on splice site definition

Both intron and exon definition models as well as the adjusted exon definition model were fitted to the single mutation effects on the isoforms frequencies inferred from our RNA-seq data (see Section 1.1). Since our microscopic model does not handle non-standard isoforms (e.g., those using cryptic splice sites or those including insertion/deletion mutations), mutations that have a significant effect on the measured non-standard isoforms ('others') were filtered out from the dataset. More precisely, we keep mutations with $p_{others} < 2\, p_{others}^{wt}$, where $p_{others}$ denotes the frequency of non-standard isoforms in the mutated gene and $p_{others}^{wt}$ denotes the corresponding mean frequency in the wildtype minigenes. Furthermore, the frequencies of the five canonical isoforms (inclusion, skipping, full IR, first IR and second IR) were re-normalized, such that their total summed up frequency is one, like expected in the model.

In total, 1854 single point mutation effects and the wildtype values of five canonical splice isoforms were fitted. We assume that each mutation may affect spliceosome binding to splice sites, and thus allowed all recognition probabilities to be fitted independently for each mutation and the wildtype. For the intron definition model in Eqs. (5), we have four unknown parameters per mutation, namely the recognition probabilities of the 3’ and 5’ ends of each intron $p_2, p_3, p_4, p_5$. The recognition probability $p_6$ does not affect splicing outcomes in this model. For the exon definition model in Eqs. (9), there are only three parameters per mutation, that correspond to the joint recognition probabilities of the three exons $p_{12}, p_{34}, p_{56}$. The adjusted definition model in Eqs. (17) includes five additional parameters, i.e., the ratios of the degradation to splicing rates of the five splice products $d_1/s, \ldots, d_5/s$. These five parameters are assumed to have the same values for all mutations and the wildtype (i.e., mutations effects on degradation rates are neglected), and were thus fitted globally to all mutation and wildtype measurements.

The total number of parameters was therefore 7420, 5565 and 5570 for the intron, exon and adjusted exon definition models, respectively. Fitting was done by minimizing the squared norm of the difference vector between model (Eqs. (5), (9) and (13), respectively) and measured values using Matlab subroutine lsqnonlin. Starting parameter values were sampled using latin hypercube sampling. Minimization was repeated for 100 sets of starting parameters, which led to stable results for the optimal parameter values.

Optimization was done while restricting parameter values to [0,1] for all recognition probabilities. For the adjusted exon definition model, the degradation rates $d_1, d_2, d_4, d_5$ of the isoforms inclusion, skipping, first and second intron retention were constrained to be between 0.3 and 3.33 times the degradation rate $d_3$ of the full intron retention isoform. The ratio $d_3/s$ was at its turn constrained between 0 and 3.33. The best-fit values of the global parameters were

$d_1/s=0.059, d_2/s=0.079, d_3/s=0.197, d_4/s=0.562, d_5/s=0.648$. Thus, inclusion and skipping isoforms were more stable compared to retention isoforms as expected based on the literature (*12*). The best-fit values of the recognition probabilities were $p_{12}^{wt}=0.60, p_{34}^{wt}=0.79, p_{56}^{wt}=0.73$for the wildtype. The corresponding values for all single mutation effects are summarized in Table S1.

### 3 Analysis of alternative exon modularity

In the exon definition model, exons function as modules as the recognition parameters corresponding to the surrounding splice sites enter the overall exon recognition parameter in a multiplicative fashion (Eqs. (8)). Hence, mutations should have identical effects on splice isoforms, irrespective of whether they target the 3' and 5' splice sites of the exon, which would not be the case for the intron definition model in Eqs. (5). To confirm the predicted modularity, we analyzed mutations nearby splice sites, sorted them according to mutation strength and compared their effects on splice isoforms (Fig. 4B and Fig. S3). In line with the exon definition hypothesis, we found that 3' and 5'mutations in the alternative exon have almost identical effects on splicing outcomes.

For the plots in Fig. 4B, mutations in a +/- 30-nt window around the 5' and 3' splice sites of the AE 11 were selected (164 mutations at positions 266-325 and 157 mutations at positions 415-474, respectively). The best-fit parameters of the adjusted exon definition model in Eqs. (17) were used to sort these mutations according to their effect on the recognition probability of the alternative exon $p_{34}$ (reflecting the mutation strength), and classified the mutations into 10 uniform bins between $p_{34}=0$ and $p_{34}=1$ (e.g. the 5th bin contains mutations that lead to an exon recognition probability $p_{34}$between 0.4 and 0.5). The top plots in Fig. 4B show the experimentally measured splice isoform frequencies as the mean in each bin vs. bin center value of $p_{34}$. Mutations in the vicinity of the left and right AE splice sites induce similar effects, supporting the concept that the alternative exon is recognized as a module. The plots below show simulations of the adjusted exon definition model (middle) as well as intron definition model (bottom). For the exon definition model, the degradation to splice rates $d_1/s, \ldots, d_5/s$are fixed to the best-fit values. For both models, the recognition probabilities $p_{12}, p_{56}$and $p_2, p_5$respectively, take the best-fit values obtained for the wildtype minigenes. To mimic 5' regulation of the alternative exon, the parameter $p_3$is varied from 0 to 1 in the left plots, while keeping $p_4=0.9$. In the right column, 3' regulation is simulated by varying $p_4$ from 0 to 1 while keeping $p_3=0.9$.

Similar plots for mutations within a +/- 30-nt window around the 2nd, 5th and 6th splice sites are shown in Fig. S3. For these plots, 168 mutations within positions 180-240 (left column), 132 mutations within positions 490-550 (middle column) and 127 mutations within positions 660-720 (right column) were selected. The best-fit parameters of the adjusted exon definition model in Eqs. (17) were used to sort these mutations according to their effect on the recognition probability of the constitutive exons ($p_{12}$ in the left column and $p_{56}$ in the middle and right columns) . For the data plots (top row), mutations were classified as already described in uniform bins according to the values of $p_{12}$ and $p_{56}$, respectively. For the adjusted exon definition model (middle row) the parameters $p_{34}, p_{56}$(left column) or $p_{12}, p_{34}$(middle and right columns) were fixed at their best-fit values. For the intron definition model (bottom row) the parameters $p_3, p_4, p_5$(left column) or $p_2, p_3, p_4$(middle and right columns) were fixed at their best fit values. To mimic the regulation of the constitutive exon 10, the parameter $p_2$is varied between 0 and 1 in the left column. To mimic the regulation of the constitutive exon 12, $p_5$(middle column) or $p_6$ (right column) are varied between 0 and 1, while keeping $p_6=0.9$ or $p_5=0.9$, respectively. The available data shows similar isoform distribution for mutations on both sites of the constitutive exon 12 (top, middle and right), supporting again the modular character of splicing reflected by the exon definition model in contrast to the intron definition model. Note that data on mutated 5th splice site is partially missing (top, middle), since these mutations lead to the activation of a cryptic splice site and formation of non-canonical isoforms, that are not currently described by the model.

### 4 Prediction of combined mutation effects

To further support modular exon regulation, we assessed whether the adjusted exon definition model trained on single mutation effects (Fig. 3B, right) could successfully predict combined mutation effects. For the plots in Fig. 4C, we focused on predicting a subset of 908 minigenes in our mutagenesis data that contained only two point mutations. However, most of these individual mutations do not have a strong effect on RON splicing, so that the prediction of their combined effect would be trivial. We therefore further filtered for those single point mutations that induce more than 20% change with respect to wildtype taken as a sum of all absolute isoform changes ($\sum \left|p_i - p_i^{wt}\right| > 0.2$) and ended up with 45 double mutation minigenes that could be used to test the exon definition model.

The combined effect of two mutations was calculated based on the assumption that mutations multiplicatively affect the $k^{on}/k^{off}$ ratio of the spliceosome binding to the single exons. Thus, in similarity to Eqs. (3), we assume for each exon recognition probability in a double mutation minigene:

$$p_{combined} = \frac{k^{on}_{combined}/k^{off}_{combined}}{1 + k^{on}_{combined}/k^{off}_{combined}}, \qquad (27)$$

with

$$\frac{k^{on}_{combined}}{k^{off}_{combined}} = \left(\frac{k^{on}_{wt}}{k^{off}_{wt}}\right)^{-1} \frac{k^{on}_1}{k^{off}_1} \cdot \frac{k^{on}_2}{k^{off}_2} \qquad (28)$$

and

$$\frac{k^{on}_i}{k^{off}_i} = \frac{p_i}{1 - p_i}, i = wt, 1, 2. \qquad (29)$$

In (28) and (29) $p_{wt}, k^{on}_{wt}, k^{off}_{wt}$ are the exon definition probabilities and binding/unbinding rates fitted to the wildtype and $p_i, k^{on}_i, k^{off}_i, i = 1,2$ are the corresponding values associated with the single mutation effects. Thus, Eqs. (27-29) allow us to calculate the exon recognition probabilities in the double mutation minigene from the estimated exon recognition probabilities in the single mutation minigenes. Finally, predictions of the five isoform frequencies in minigenes containing two mutations are made using the models in Eqs. (5) and (17), respectively. These predictions (in total 225 isoform values for each model) are compared to the experimental measurements in Fig. 4C (right panel).

For the left panel in Fig. 4, the analysis was further restricted based on the positions of mutations: specifically, only mutations in a +/- 30 nt window around the 3' and 5' sites of the AE were considered (positions 275-330 and 415-475). This leads to a subset of 3 double mutation minigenes (in total 15 isoform value predictions for each model).

### 5 Simulation of mutation effects on PSI and total intron retention

We used the model to investigate how alternative splicing regulation relates to intron retention. For simplicity, we did not distinguish intron retention isoforms but considered only total intron retention, i.e., the sum of all retention products. Considering that all isoform frequencies sum up to 1, we derive an analytical expression for *totalIR* from Eqs. (9)

$$totalIR = 1 - p_{inclusion} - p_{skipping} = 1 - p_{12}\, p_{56}\,, \quad (30)$$

For the PSI of the middle exon as a measure of alternative splicing, we obtain

$$PSI = \frac{p_{inclusion}}{p_{inclusion} + p_{skipping}} = p_{34}. \quad (31)$$

Hence, alternative splicing and the degree of intron retention are controlled independently in the exon definition model (if all splice isoforms are degraded turned over with the same rates), and alternative splicing regulation occurs without the accumulation of retention products.

For the intron definition model *totalIR* and *PSI* cannot be uncoupled in the same way. Specifically, we obtain from Eq. (7):

$$totalIR = 1 - 2\, p_2 p_3 p_4 p_5 - p_2 p_5 + p_2 p_3 p_5 + p_2 p_4 p_5\,,$$
$$PSI = \frac{p_3 p_4}{2\, p_3 p_4 + 1 - p_3 - p_4}. \quad (32)$$

If both splice sites flanking the alternative exon are jointly recognized ($p_3$=$p_4$=0), or jointly not recognized ($p_3$=$p_4$=1), we have

$$totalIR = 1 - p_2 p_5\,, \quad (33)$$

which resembles Eq. (30). In contrast to the exon definition model, *totalIR* approaches 1 if only one of the splice sites is recognized ($p_3 = 1 \wedge p_4 = 0, \vee\, p_3 = 0 \wedge p_4 = 1$). Hence, both splice sites need to be regulated jointly to prevent retention in the intron definition model.

However, even in this scenario of joint regulation ($p_3 = \alpha\, p_4$), intron retention isoforms accumulate at intermediate *PSI* values, i.e., during the switch from exon skipping to inclusion. Plugging $p_3 = \alpha\, p_4$into Eqs. (32) yields

$$totalIR = 1 - \frac{2\alpha}{(1+\alpha)^2} \quad (34)$$

for $PSI = 0.5$and$p_2 = p_5 = 1$. Hence, even if the splice sites of the outer exons are perfectly recognized ($p_2 = p_5 = 1$) *totalIR* will accumulate to at least 50% of all splice products at intermediate inclusion levels (PSI=0.5). Less complete recognition of the outer exons results in even higher retention (not shown), implying alternative splicing regulation in the intron definition model inevitably results in the accumulation of retention products.

To visualize the distinct accumulation of retention products in the intron and exon definition models, Monte-Carlo simulations mimicking point mutation effects were performed (Fig. 4D, middle and right). To this end, 2000 sets of binding probabilities ($p_1, \dots, p_6$) were used. In all runs, five of the six binding probabilities ($p_2 - p_6$) are drawn from a Gauss distribution (standard deviation 5%). In 45 of the runs, one of the constitutive exons splice sites (2,5,6) is randomly chosen and its binding probability additionally perturbed (standard deviation 50%). In other 300 of the runs, one of the alternative exons splice sites (3 or 4) is randomly chosen and the corresponding binding probability is additionally perturbed (standard deviation 50%). The distributions for $p_2 - p_6$ are centered around (0.95,0.7,0.7,0.95,0.95) and (0.9, 0.9, 0.9, 0.9, 0.9) for the intron and exon definition models, respectively. These values were chosen to mimic the experimentally measured wildtype PSI and intron retention values. The value of $p_1$ is set to 1 in both models since the minigene used in the experiment does not include the 1st splice site.

### 6 Analysis of splicing for a pre-mRNA containing four exons

In this section, we theoretically analyze splicing of a four-exon gene to show that the same principles we derived above also hold for longer sequences and more complex splicing scenarios.

Consider a gene with four exons in which the recognition is denoted in a binary fashion, e.g., (1111) and (0000) for the fully bound (all exons or recognized) and empty states (no exon recognized), respectively. We define the sum of the states (1111), (1011), (1101) and (1001) as the *productive states*, as only for these cases no intron will be retained after splicing (following the same splicing rules as in Section 2).

With $p_i$ being the individual recognition probability of each exon, we obtain for the frequency of total intron retention, i.e., the sum of all retention isoforms

$$totalIR = 1 - productive\,states = 1 - p_1 p_4 \qquad (35)$$

and

$$productive\,states = p_1 p_4. \qquad (36)$$

This expression resembles Eq. (30) and shows that *totalIR* solely depends on the recognition of the outer exons.

The percent spliced-in (*PSI*) for the second and third (i.e., the inner) exons is given by

$$\begin{aligned} PSI_2 &= \frac{p_{1111} + p_{1101}}{productive\,states} = p_2, \\ PSI_3 &= \frac{p_{1111} + p_{1011}}{productive\,states} = p_3. \end{aligned} \qquad (37)$$

Thus, as for the three-exon case (Eq. (31)), alternative splicing is controlled independently of *totalIR*. Furthermore, alternative splicing regulation is modularized in two ways: (i) By assigning a total recognition probability to each exon ($p_i$), we implicitly assumed that splice signals acting on the two splice sites of this exon are integrated into a net outcome (within-exon modularity); (ii) According to Eqs. (37), the inclusion level of exon is controlled only by its own recognition probability independent of its neighboring exons. This gives rise to an additional cross-exon modularity that was not present in the three-exon case (Section 2).

Similar conclusions also hold for more complex scenarios involving more than four exons (as long as we assume that exon skipping can also occur across long sequences comprising multiple introns and exons). Taken together, these calculations support that exon definition is generally beneficial for the alternative splicing regulation, also for long pre-mRNAs with complex exon structure.

### 7 Analysis of the *HNRNPH* knockdown response and synergistic effects

In order to analyze how *HNRNPH* affects RON splicing, we quantitatively analyzed hnRNP knockdown data in HEK293 cells using our model. The distribution of the exon recognition probabilities in wildtype minigenes and the mutation effects on these probabilities were calculated by fitting the adjusted exon definition model to the *HNRNPH* knockdown data as described in Section 3. Here, the global parameters $d_1/s, \ldots, d_5/s$ were kept at the values determined for control conditions.

The distribution of the exon recognition probabilities $p_{12}, p_{34}, p_{56}$ for the wildtype minigenes is shown in Fig. S4A. The knockdown mainly increases the recognition probability of the alternative exon $p_{34}$, while the recognition probabilities of the constitutive exons $p_{12}, p_{56}$ are comparable to control values.

In order to identify which sites are most relevant for HNRNPH-dependent regulation, we inferred the splicing response of single point mutation variants upon HNRNPH knockdown using our model. We hypothesized that mutations that either weaken or reinforce an HNRNPH binding site would display positive or negative synergy with the HNRNPH knockdown. For instance, a reduced knockdown response compared to the wildtype minigene would be expected if an important HNRNPH binding site is compromised by a mutation (negative synergy). For the exon recognition probabilities $p_i, i=12,34,56$ in mutated minigenes, a z-score was calculated based on the mean and standard deviation in wildtype minigenes:

$$z_i = \frac{\log\left(p_i^{KD}/p_i^{control}\right) - wtmean}{wtstd}, \qquad (38)$$

where *wtmean* and *wtstd* are the mean and standard deviation of $\log\left(p_i^{KD}/p_i^{control}\right)$ in wildtype minigenes. Mutations with $|z_i|>5$ indicate a highly distinct knockdown response in wildtype and mutant background and were considered as a synergistic interaction which hints to the presence of a cis-regulated element targeted by *HNRNPH*. The density of such interactions in a 5-nt sliding window is plotted in Fig. S4B. The most reliable synergistic interactions are found for the recognition probability $p_{34}$ of the alternative exon with corresponding mutations located in the alternative exon or flanking introns. For a similar analysis in MCF7 cells see also Fig. 5 in (*11*).
