## Supplementary Figure 1 for "Exon definition facilitates reliable control of alternative splicing in the *RON* proto-oncogene"

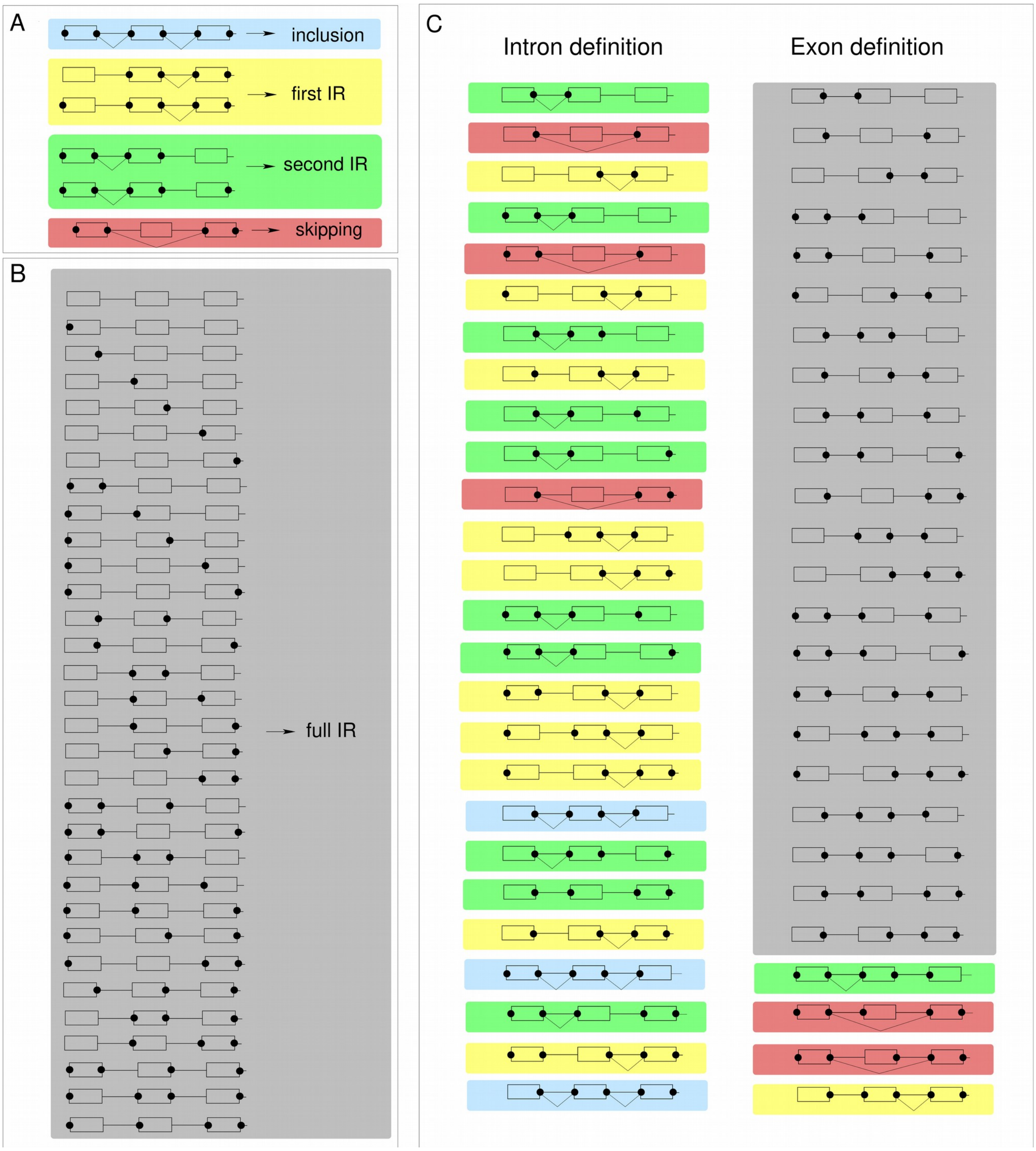


**Fig. S1. Common and distinct splicing decisions in intron vs. exon definition models. A** Spliceosomal binding states leading to identical splicing outcomes in both models. **B** Binding states with no matching 3' and 5' splice sites across introns. Since no splicing reaction can take place, these binding states lead to the full intron retention isoform in both models. **C** Binding states that lead to different splice outcomes in the intron (left column) and exon definition (right column) model. Black dots indicate U1 and U2 binding to splice sites. Splicing reactions are indicated by triangular black lines. The background color reflects the splice outcome (blue: inclusion, red: skipping, yellow: first IR, green: second IR, gray: full IR).
