## Supplementary Figure 2 for "Exon definition facilitates reliable control of alternative splicing in the *RON* proto-oncogene"

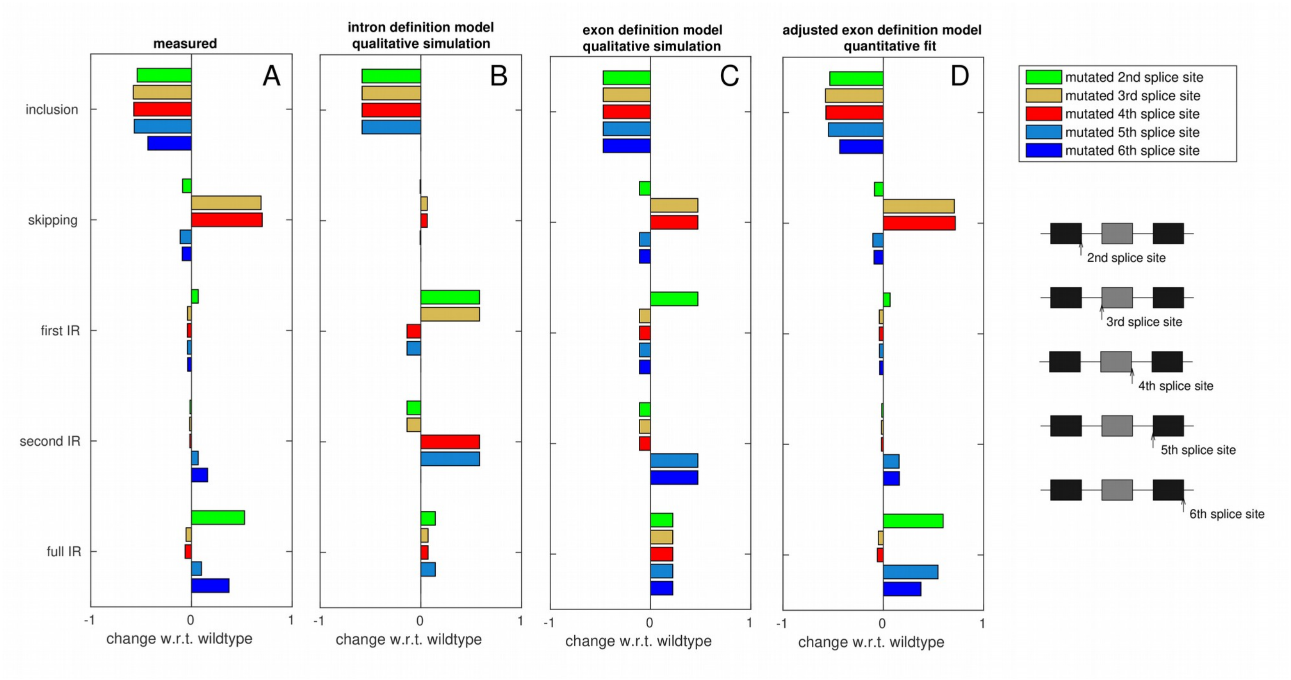


**Fig. S2. Splice site mutation effects on splice isoforms support the exon definition model. A** Bar plots showing the median isoform differences w.r.t. wildtype for directly measured minigenes containing single point mutations at different splice sites. Splice sites are defined as the first and last two positions within an intron (sites 211-212,296-297,445-446,523-524,691-692). Mutations at these positions have the strongest effect on splicing outcomes within the library. A subset of this data is shown in Fig. 3D. **B,C** Predictions of splice site mutation effects based on qualitative simulations of the intron and exon definition models, respectively. Importantly, the directionality of the simulated isoform changes hold true for arbitrary sets of splice recognition parameters ( see Sections 1.1 and 1.2 for details). The exon definition (but not the intron definition) model reproduces 23 out of 25 splice mutation effects, and only fails to describe AE splice site effects on full IR. **D** The adjusted exon definition model with isoform-specific degradation and fitted parameters (Fig. 3B, right) qualitatively and quantitatively reproduces all measured splice site effects.
