## Supplementary Figure 3 for "Exon definition facilitates reliable control of alternative splicing in the *RON* proto-oncogene"

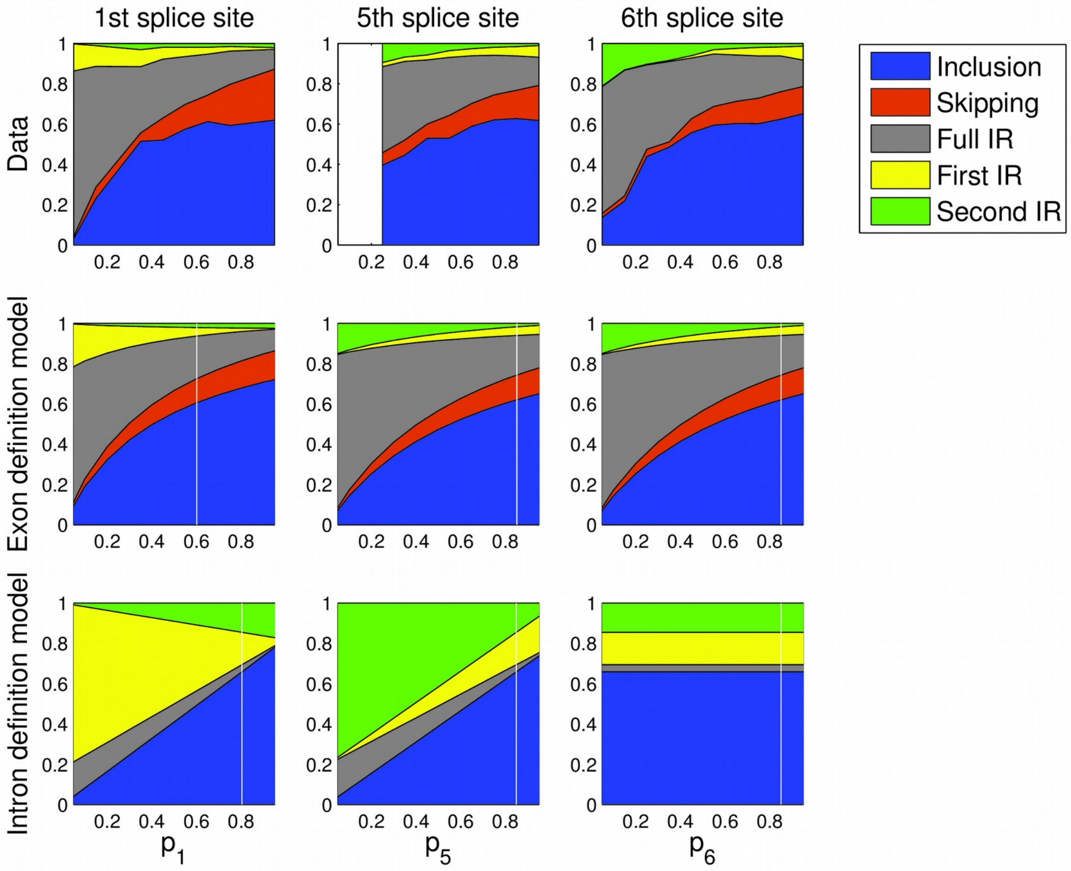


**Fig. S3. Position-dependent mutation effects on splice isoforms support the exon definition model (related to Fig. 4B).** Plots show a similar analysis as in Fig. 4B for the other splice sites. Mutations at positions in a +/- 30-nt window around the 2nd, 5th and 6th splice sites were selected. The mutations were sorted according to their effect on the recognition probability $p_{12}$(left) or $p_{56}$ (middle and right) in the best-fit model (adjusted exon definition model; Fig. 3B, right). The measured changes in the splice isoform fractions with varying mutation strength (1$^{st}$ row) are similar for the 5th and 6th splice splice sites, and agree with simulations of the exon definition model in which $p_{56}$ is systematically varied (2$^{nd}$ row), but disagree with the intron definition model (3$^{rd}$ row). Furthermore, mutations at the 2nd splice site induce similar changes as mutations at the 5th and 6th splice sites for the inclusion, skipping and full intron retention isoforms, as reflected by the exon definition model in contrast to the intron definition model. The vertical white line is located at the best-fit parameter values corresponding to wildtype minigenes.
