## Supplementary Figure 4 for "Exon definition facilitates reliable control of alternative splicing in the *RON* proto-oncogene"

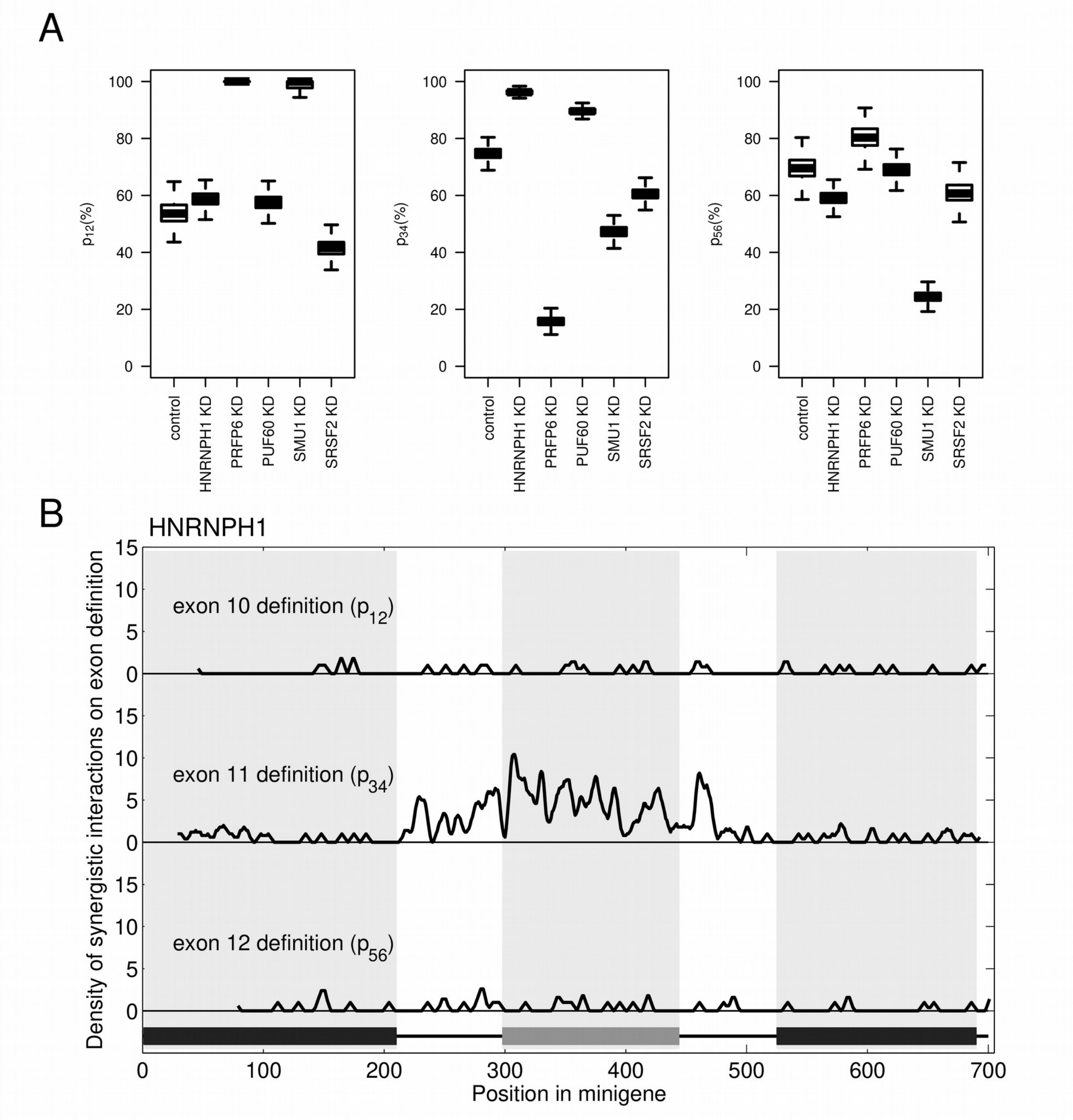


**Fig. S4. Knockdown of different RNA binding proteins affects mainly the recognition probability of the alternative exon. A** Boxplots show the distribution of the best-fit recognition probabilities $p_{12}, p_{34}, p_{56}$ of the three exons in wildtype minigenes under control and *HNRNPH1, PRPF6, PUF60, SMU1* or *SRSF2* knockdown conditions. The fit was generated using the adjusted exon definition model (see Section 7 for details). **B** *HNRNPH* may target cis-regulatory elements in the alternative exon, as evidenced by a synergistic interaction of *HNRNPH* knockdown with point mutations in this region. Maps show the density of synergistic mutation-knockdown effects on the three exon recognition probabilities. Single mutation effects on the exon recognition were determined separately for control and *HNRNPH* knockdown conditions. The difference (in logarithmic space) between control and *HNRNPH* knockdown values for single point mutations was compared to the knockdown response of wildtype minigenes and expressed as a z-score relative to the within-wildtype-variation (see Section 7 and (11)). Large z-scores indicate a synergistic interaction with the *HNRNPH* knockdown. The density of mutations with a z-score $|z| > 5$ in a 5-nt sliding window is plotted.
