## Supplementary Figure 5 for "Exon definition facilitates reliable control of alternative splicing in the *RON* proto-oncogene"

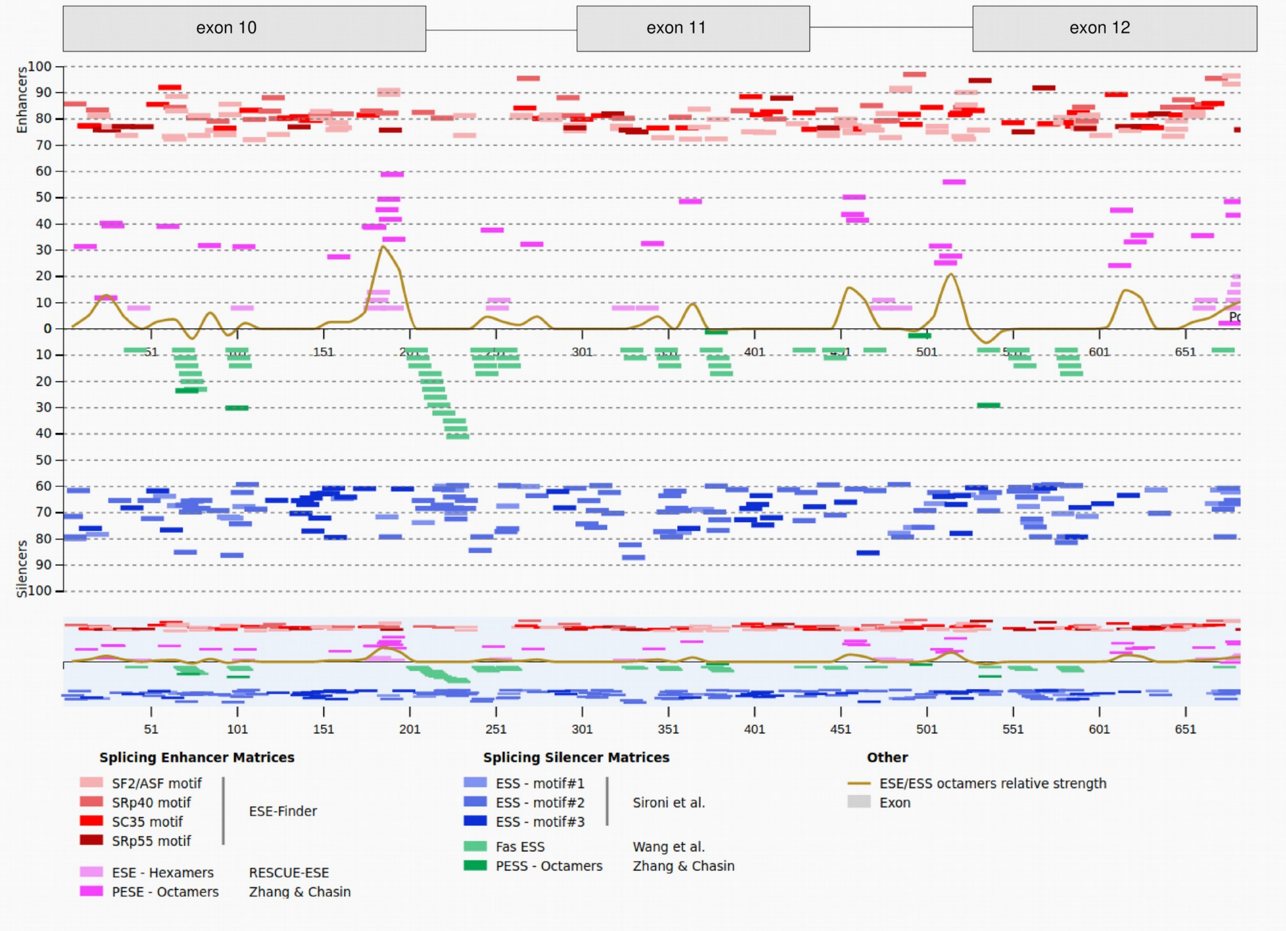


**Fig. S5 Summary of exonic splice enhancers (ESE) and exonic splice silencers (ESS) predicted for the wildtype minigene sequence.** The plot was generated with the Human Splicing Finder, Version 3.1. Results are also summarized in Table S2.
