## Supplementary material for "Exon definition facilitates reliable control of alternative splicing in the *RON* proto-oncogene": Table S1

### Best Fit Values of Model Parameters

| | Exon Definition Model | | | Intron Definition Model | | | |
|---|---|---|---|---|---|---|---|
| | $p_{12}$ | $p_{34}$ | $p_{56}$ | $p_2$ | $p_3$ | $p_4$ | $p_5$ |
| wildtype | 0.60 | 0.79 | 0.73 | 0.80 | 1.00 | 0.97 | 0.85 |
| A26T | 0.58 | 0.79 | 0.71 | 0.80 | 0.99 | 1.00 | 0.81 |
| A26G | 0.58 | 0.79 | 0.61 | 0.77 | 0.99 | 0.99 | 0.78 |
| G27T | 0.67 | 0.71 | 0.77 | 0.95 | 0.84 | 0.88 | 0.92 |
| G27C | 0.69 | 0.76 | 0.66 | 0.82 | 0.99 | 0.99 | 0.82 |
| G27A | 0.53 | 0.78 | 0.67 | 0.76 | 0.99 | 0.99 | 0.77 |
| C28T | 0.58 | 0.82 | 0.70 | 0.81 | 1.00 | 1.00 | 0.82 |
| C28G | 0.40 | 0.64 | 0.97 | 0.85 | 0.79 | 0.82 | 0.88 |
| C28A | 0.66 | 0.75 | 0.63 | 0.80 | 1.00 | 0.99 | 0.80 |
| T29G | 0.61 | 0.76 | 0.75 | 0.83 | 0.97 | 0.98 | 0.83 |
| T29C | 0.63 | 0.79 | 0.70 | 0.81 | 0.99 | 0.99 | 0.83 |
| T29A | 0.54 | 0.78 | 0.66 | 0.76 | 0.99 | 1.00 | 0.78 |
| G30A | 0.56 | 0.79 | 0.73 | 0.80 | 0.99 | 1.00 | 0.81 |
| T31G | 0.61 | 0.79 | 0.73 | 0.81 | 1.00 | 0.99 | 0.83 |
| T31C | 0.61 | 0.81 | 0.71 | 0.81 | 1.00 | 0.99 | 0.83 |
| T31A | 0.61 | 0.79 | 0.67 | 0.81 | 0.98 | 1.00 | 0.81 |
| G32T | 0.60 | 0.80 | 0.68 | 0.80 | 1.00 | 0.99 | 0.81 |
| G32C | 0.44 | 0.61 | 0.75 | 0.71 | 0.80 | 0.80 | 0.73 |
| G32A | 0.60 | 0.76 | 0.67 | 0.78 | 0.99 | 0.99 | 0.79 |
| C33T | 0.62 | 0.78 | 0.77 | 0.83 | 0.99 | 0.98 | 0.85 |
| C33G | 0.78 | 0.65 | 0.60 | 0.90 | 0.86 | 0.84 | 0.90 |
| C33A | 0.68 | 0.88 | 0.78 | 0.87 | 1.00 | 0.99 | 0.89 |
| C34T | 0.56 | 0.78 | 0.63 | 0.76 | 0.99 | 0.99 | 0.77 |
| C34A | 0.51 | 0.85 | 0.73 | 0.79 | 1.00 | 0.96 | 0.86 |
| G35T | 0.49 | 0.80 | 0.73 | 0.76 | 0.99 | 1.00 | 0.79 |
| G35A | 0.57 | 0.81 | 0.71 | 0.81 | 0.99 | 1.00 | 0.82 |
| C36T | 0.61 | 0.84 | 0.77 | 0.84 | 1.00 | 1.00 | 0.86 |
| C36G | 0.54 | 0.73 | 0.69 | 0.74 | 0.99 | 0.98 | 0.77 |
| C36A | 0.66 | 0.77 | 0.79 | 0.89 | 0.93 | 0.97 | 0.87 |
| C37T | 0.61 | 0.79 | 0.71 | 0.82 | 0.98 | 0.99 | 0.83 |
| C37A | 0.53 | 0.79 | 0.71 | 0.77 | 1.00 | 0.98 | 0.81 |
| CT38C | 0.54 | 0.79 | 0.79 | 0.80 | 0.99 | 0.98 | 0.83 |
| T38G | 0.65 | 0.73 | 0.71 | 0.79 | 1.00 | 0.99 | 0.81 |
| T38C | 0.57 | 0.79 | 0.70 | 0.79 | 0.99 | 0.99 | 0.81 |
| T38A | 0.62 | 0.80 | 0.74 | 0.82 | 1.00 | 0.99 | 0.84 |
| T39C | 0.59 | 0.78 | 0.72 | 0.80 | 0.99 | 0.98 | 0.83 |
| T39A | 0.64 | 0.73 | 0.70 | 0.81 | 0.97 | 0.98 | 0.82 |
| TC40T | 0.59 | 0.75 | 0.72 | 0.78 | 1.00 | 0.96 | 0.82 |
| C40T | 0.49 | 0.74 | 0.64 | 0.71 | 0.97 | 0.98 | 0.72 |
| C40A | 0.58 | 0.56 | 0.81 | 0.90 | 0.72 | 0.71 | 0.94 |
| C41T | 0.55 | 0.77 | 0.79 | 0.80 | 0.99 | 0.99 | 0.82 |
| C41G | 0.69 | 0.77 | 0.68 | 0.84 | 0.98 | 1.00 | 0.82 |
| C41A | 0.59 | 0.79 | 0.72 | 0.80 | 0.99 | 0.99 | 0.81 |
| T42G | 0.65 | 0.81 | 0.68 | 0.82 | 1.00 | 0.97 | 0.85 |
| T42C | 0.64 | 0.82 | 0.67 | 0.82 | 1.00 | 0.98 | 0.84 |
| T42A | 0.57 | 0.82 | 0.77 | 0.82 | 1.00 | 0.99 | 0.85 |
| G43T | 0.55 | 0.76 | 0.46 | 0.70 | 0.79 | 0.98 | 0.56 |
| G43C | 0.55 | 0.78 | 0.72 | 0.77 | 1.00 | 0.99 | 0.80 |
| G43A | 0.57 | 0.78 | 0.73 | 0.79 | 0.99 | 0.98 | 0.82 |
| A44T | 0.47 | 0.80 | 0.71 | 0.80 | 0.92 | 1.00 | 0.77 |
| A44G | 0.53 | 0.77 | 0.74 | 0.77 | 0.99 | 0.98 | 0.80 |

| | | | | | | | |
|---|---|---|---|---|---|---|---|
| A44C | 0.51 | 0.73 | 0.68 | 0.72 | 1.00 | 0.98 | 0.76 |
| A45T | 0.59 | 0.82 | 0.72 | 0.83 | 0.98 | 1.00 | 0.83 |
| A45G | 0.60 | 0.82 | 0.66 | 0.81 | 1.00 | 0.99 | 0.82 |
| T46G | 0.68 | 0.76 | 0.82 | 0.95 | 0.88 | 0.90 | 0.94 |
| T46C | 0.62 | 0.76 | 0.68 | 0.82 | 0.95 | 1.00 | 0.79 |
| T46A | 0.58 | 0.80 | 0.77 | 0.82 | 0.99 | 0.99 | 0.84 |
| A47T | 0.56 | 0.71 | 0.79 | 0.83 | 0.92 | 0.96 | 0.82 |
| A47G | 0.54 | 0.74 | 0.82 | 0.80 | 0.97 | 0.96 | 0.84 |
| T48G | 0.62 | 0.97 | 0.82 | 0.91 | 0.98 | 1.00 | 0.92 |
| T48C | 0.59 | 0.75 | 0.78 | 0.82 | 0.97 | 0.97 | 0.83 |
| T48A | 0.61 | 0.78 | 0.61 | 0.78 | 0.99 | 1.00 | 0.78 |
| G49C | 0.73 | 0.88 | 0.76 | 0.89 | 1.00 | 1.00 | 0.90 |
| G49A | 0.68 | 0.82 | 0.76 | 0.86 | 0.99 | 0.99 | 0.88 |
| T50G | 0.57 | 0.87 | 0.65 | 0.82 | 1.00 | 1.00 | 0.83 |
| T50C | 0.60 | 0.79 | 0.75 | 0.81 | 0.99 | 0.99 | 0.83 |
| T50A | 0.59 | 0.80 | 0.79 | 0.82 | 1.00 | 0.99 | 0.85 |
| TG51T | 0.56 | 0.80 | 0.60 | 0.75 | 1.00 | 0.99 | 0.77 |
| G51T | 0.52 | 0.85 | 0.74 | 0.80 | 1.00 | 1.00 | 0.83 |
| G51A | 0.63 | 0.83 | 0.68 | 0.83 | 1.00 | 1.00 | 0.84 |
| G52T | 0.58 | 0.82 | 0.77 | 0.82 | 1.00 | 1.00 | 0.84 |
| G52C | 0.62 | 0.78 | 0.71 | 0.81 | 0.99 | 1.00 | 0.82 |
| G52A | 0.65 | 0.75 | 0.74 | 0.82 | 0.99 | 0.98 | 0.84 |
| T53G | 0.49 | 0.82 | 0.74 | 0.77 | 0.99 | 1.00 | 0.81 |
| T53C | 0.58 | 0.79 | 0.73 | 0.81 | 0.99 | 0.98 | 0.84 |
| T53A | 0.58 | 0.81 | 0.83 | 0.83 | 0.99 | 0.99 | 0.85 |
| C54T | 0.59 | 0.80 | 0.73 | 0.81 | 0.99 | 0.99 | 0.83 |
| C54G | 0.63 | 0.64 | 0.87 | 0.98 | 0.78 | 0.79 | 0.99 |
| C54A | 0.51 | 0.86 | 0.68 | 0.79 | 1.00 | 1.00 | 0.81 |
| C55T | 0.55 | 0.79 | 0.74 | 0.78 | 1.00 | 0.98 | 0.82 |
| C55G | 0.54 | 0.82 | 0.84 | 0.83 | 0.98 | 1.00 | 0.85 |
| C55A | 0.52 | 0.81 | 0.72 | 0.78 | 0.99 | 0.99 | 0.80 |
| G56T | 0.54 | 0.66 | 0.96 | 0.97 | 0.78 | 0.80 | 0.98 |
| G56C | 0.36 | 0.79 | 0.91 | 0.73 | 0.99 | 1.00 | 0.78 |
| G56A | 0.60 | 0.78 | 0.70 | 0.80 | 0.99 | 0.99 | 0.81 |
| GA57G | 0.65 | 0.78 | 0.80 | 0.83 | 1.00 | 0.95 | 0.89 |
| A57T | 0.56 | 0.81 | 0.81 | 0.82 | 1.00 | 0.97 | 0.86 |
| A57G | 0.55 | 0.87 | 0.70 | 0.81 | 1.00 | 1.00 | 0.83 |
| G58T | 0.52 | 0.68 | 0.80 | 0.81 | 0.89 | 0.88 | 0.85 |
| G58A | 0.61 | 0.76 | 0.68 | 0.78 | 1.00 | 0.99 | 0.80 |
| A59T | 0.67 | 0.75 | 0.60 | 0.79 | 0.99 | 1.00 | 0.78 |
| A59G | 0.64 | 0.80 | 0.70 | 0.82 | 1.00 | 0.99 | 0.84 |
| AC60A | 0.54 | 0.73 | 0.70 | 0.77 | 0.96 | 0.98 | 0.77 |
| C60T | 0.71 | 0.83 | 0.65 | 0.85 | 1.00 | 1.00 | 0.85 |
| C60A | 0.57 | 0.75 | 0.68 | 0.76 | 1.00 | 0.98 | 0.79 |
| C61T | 0.59 | 0.78 | 0.76 | 0.82 | 0.98 | 1.00 | 0.83 |
| C61G | 0.61 | 0.94 | 0.77 | 0.87 | 1.00 | 1.00 | 0.89 |
| C61A | 0.62 | 0.74 | 0.80 | 0.86 | 0.94 | 0.93 | 0.88 |
| C62T | 0.68 | 0.77 | 0.77 | 0.89 | 0.94 | 0.99 | 0.85 |
| C62A | 0.61 | 0.80 | 0.68 | 0.80 | 1.00 | 0.99 | 0.82 |
| C63T | 0.64 | 0.78 | 0.71 | 0.82 | 0.99 | 0.99 | 0.83 |
| C63G | 0.45 | 0.73 | 1.00 | 0.97 | 0.76 | 0.83 | 0.98 |
| C63A | 0.62 | 0.86 | 0.62 | 0.82 | 1.00 | 1.00 | 0.82 |
| C63CT | 0.55 | 0.83 | 0.73 | 0.83 | 0.96 | 1.00 | 0.83 |

| | | | | | | | |
|---|---|---|---|---|---|---|---|
| C64T | 0.48 | 0.79 | 0.88 | 0.79 | 1.00 | 0.99 | 0.84 |
| C64A | 0.58 | 0.82 | 0.69 | 0.80 | 1.00 | 0.99 | 0.82 |
| C64CA | 0.66 | 0.74 | 0.78 | 0.90 | 0.90 | 0.91 | 0.91 |
| CA65C | 0.50 | 0.67 | 0.89 | 0.88 | 0.84 | 0.87 | 0.88 |
| A65T | 0.61 | 0.79 | 0.75 | 0.82 | 0.99 | 0.99 | 0.83 |
| A65G | 0.61 | 0.84 | 0.73 | 0.84 | 0.98 | 1.00 | 0.84 |
| A65C | 0.58 | 0.81 | 0.73 | 0.81 | 1.00 | 0.99 | 0.83 |
| AG66A | 0.57 | 0.77 | 0.67 | 0.77 | 0.99 | 0.99 | 0.77 |
| G66T | 0.68 | 0.75 | 0.78 | 0.92 | 0.90 | 0.93 | 0.90 |
| G66A | 0.63 | 0.79 | 0.73 | 0.82 | 1.00 | 0.98 | 0.84 |
| G67T | 0.67 | 0.77 | 0.79 | 0.90 | 0.92 | 0.97 | 0.87 |
| G67C | 0.82 | 0.61 | 0.79 | 1.00 | 0.78 | 0.78 | 1.00 |
| G67A | 0.69 | 0.79 | 0.76 | 0.86 | 0.98 | 0.97 | 0.87 |
| G68A | 0.70 | 0.81 | 0.78 | 0.86 | 0.99 | 0.98 | 0.88 |
| A69T | 0.59 | 0.85 | 0.70 | 0.82 | 1.00 | 1.00 | 0.84 |
| A69G | 0.57 | 0.82 | 0.67 | 0.80 | 0.99 | 1.00 | 0.81 |
| A69C | 0.68 | 0.89 | 0.77 | 0.88 | 1.00 | 0.99 | 0.90 |
| A69AT | 0.47 | 0.80 | 0.87 | 0.81 | 0.97 | 1.00 | 0.82 |
| T70G | 0.69 | 0.86 | 0.74 | 0.87 | 1.00 | 1.00 | 0.87 |
| T70C | 0.60 | 0.81 | 0.73 | 0.82 | 0.99 | 0.99 | 0.84 |
| T70A | 0.57 | 0.86 | 0.68 | 0.81 | 1.00 | 1.00 | 0.82 |
| TG71T | 0.62 | 0.83 | 0.75 | 0.84 | 1.00 | 0.99 | 0.86 |
| G71C | 0.68 | 0.81 | 0.90 | 0.95 | 0.91 | 0.93 | 0.96 |
| G71A | 0.58 | 0.81 | 0.74 | 0.81 | 1.00 | 0.99 | 0.84 |
| G72T | 0.67 | 0.78 | 0.67 | 0.82 | 0.99 | 0.99 | 0.82 |
| G72C | 0.71 | 0.81 | 0.73 | 0.86 | 0.99 | 0.99 | 0.86 |
| G72A | 0.63 | 0.79 | 0.69 | 0.81 | 0.99 | 0.99 | 0.82 |
| G73T | 0.70 | 0.78 | 0.81 | 0.91 | 0.93 | 0.93 | 0.92 |
| G73C | 0.67 | 0.89 | 0.80 | 0.89 | 0.99 | 1.00 | 0.89 |
| G73A | 0.70 | 0.78 | 0.80 | 0.93 | 0.91 | 0.91 | 0.93 |
| T74G | 0.43 | 0.70 | 0.99 | 0.85 | 0.87 | 0.83 | 0.96 |
| T74C | 0.63 | 0.77 | 0.72 | 0.82 | 0.99 | 0.97 | 0.83 |
| T74A | 0.61 | 0.79 | 0.68 | 0.81 | 0.98 | 0.99 | 0.82 |
| T74TG | 0.66 | 0.76 | 0.73 | 0.82 | 0.99 | 0.98 | 0.84 |
| G75T | 0.55 | 0.81 | 0.79 | 0.81 | 1.00 | 1.00 | 0.84 |
| G75A | 0.63 | 0.79 | 0.75 | 0.89 | 0.92 | 1.00 | 0.84 |
| G76T | 0.64 | 0.78 | 0.68 | 0.80 | 1.00 | 0.96 | 0.84 |
| G76C | 0.66 | 0.80 | 0.72 | 0.83 | 1.00 | 0.98 | 0.85 |
| G76A | 0.66 | 0.77 | 0.76 | 0.83 | 0.99 | 0.98 | 0.86 |
| C77T | 0.60 | 0.83 | 0.66 | 0.81 | 1.00 | 1.00 | 0.82 |
| C77G | 0.56 | 0.89 | 0.44 | 0.74 | 0.97 | 0.99 | 0.71 |
| C77A | 0.62 | 0.83 | 0.70 | 0.83 | 1.00 | 0.99 | 0.84 |
| CA78C | 0.56 | 0.76 | 0.76 | 0.78 | 1.00 | 0.98 | 0.81 |
| A78T | 0.58 | 0.81 | 0.75 | 0.81 | 1.00 | 0.99 | 0.83 |
| A78G | 0.59 | 0.80 | 0.73 | 0.81 | 0.99 | 1.00 | 0.83 |
| A78C | 0.65 | 0.68 | 0.75 | 0.91 | 0.84 | 0.86 | 0.90 |
| AG79A | 0.72 | 0.82 | 0.72 | 0.85 | 1.00 | 0.97 | 0.88 |
| G79T | 0.69 | 0.77 | 0.75 | 0.85 | 0.97 | 0.97 | 0.86 |
| G79C | 0.53 | 0.76 | 0.67 | 0.74 | 0.99 | 1.00 | 0.76 |
| G79A | 0.70 | 0.79 | 0.71 | 0.84 | 0.99 | 0.99 | 0.85 |
| G80T | 0.70 | 0.79 | 0.78 | 0.87 | 0.97 | 0.97 | 0.88 |
| G80C | 0.60 | 0.71 | 0.84 | 0.92 | 0.86 | 0.86 | 0.93 |
| G80A | 0.71 | 0.81 | 0.74 | 0.86 | 0.99 | 0.97 | 0.88 |

| | | | | | | | |
|---|---|---|---|---|---|---|---|
| G81T | 0.64 | 0.72 | 0.63 | 0.77 | 1.00 | 0.99 | 0.78 |
| G81A | 0.66 | 0.79 | 0.78 | 0.86 | 0.97 | 0.98 | 0.86 |
| A82T | 0.56 | 0.80 | 0.75 | 0.80 | 1.00 | 0.99 | 0.82 |
| A82G | 0.64 | 0.80 | 0.71 | 0.82 | 1.00 | 0.99 | 0.84 |
| A82C | 0.58 | 0.80 | 0.74 | 0.80 | 1.00 | 0.99 | 0.83 |
| A83T | 0.56 | 0.79 | 0.81 | 0.81 | 0.99 | 0.99 | 0.84 |
| A83G | 0.61 | 0.79 | 0.75 | 0.82 | 0.99 | 0.99 | 0.84 |
| T84G | 0.69 | 0.83 | 0.71 | 0.85 | 1.00 | 1.00 | 0.86 |
| T84C | 0.63 | 0.80 | 0.72 | 0.82 | 0.99 | 0.99 | 0.83 |
| T84A | 0.69 | 0.78 | 0.70 | 0.83 | 1.00 | 0.99 | 0.84 |
| C85T | 0.59 | 0.67 | 0.69 | 0.82 | 0.88 | 0.91 | 0.80 |
| C85A | 0.48 | 0.75 | 0.79 | 0.78 | 0.95 | 0.99 | 0.78 |
| T86G | 0.72 | 0.70 | 0.52 | 0.79 | 0.94 | 0.96 | 0.76 |
| T86C | 0.56 | 0.72 | 0.73 | 0.78 | 0.97 | 0.97 | 0.79 |
| T86A | 0.52 | 0.71 | 0.75 | 0.75 | 0.98 | 0.93 | 0.81 |
| G87T | 0.54 | 0.72 | 0.70 | 0.75 | 0.97 | 0.97 | 0.77 |
| G87A | 0.58 | 0.79 | 0.73 | 0.81 | 0.98 | 0.99 | 0.82 |
| A88T | 0.51 | 0.72 | 0.77 | 0.76 | 0.98 | 0.97 | 0.79 |
| A88G | 0.50 | 0.76 | 0.76 | 0.75 | 0.99 | 1.00 | 0.78 |
| A88C | 0.64 | 0.75 | 1.00 | 1.00 | 0.83 | 0.86 | 1.00 |
| G89T | 0.61 | 0.73 | 0.74 | 0.80 | 0.97 | 0.93 | 0.85 |
| G89A | 0.62 | 0.83 | 0.66 | 0.82 | 1.00 | 1.00 | 0.83 |
| T90G | 0.48 | 0.76 | 0.71 | 0.73 | 0.99 | 0.99 | 0.76 |
| T90C | 0.60 | 0.76 | 0.79 | 0.89 | 0.90 | 1.00 | 0.83 |
| T90A | 0.58 | 0.77 | 0.77 | 0.81 | 0.99 | 0.99 | 0.83 |
| G91C | 0.50 | 0.76 | 0.86 | 0.80 | 0.97 | 0.99 | 0.82 |
| G91A | 0.58 | 0.79 | 0.70 | 0.80 | 0.99 | 0.99 | 0.81 |
| GC92G | 0.69 | 0.77 | 0.74 | 0.86 | 0.96 | 0.97 | 0.86 |
| C92T | 0.61 | 0.77 | 0.68 | 0.79 | 0.99 | 1.00 | 0.80 |
| C92A | 0.72 | 0.81 | 0.75 | 0.86 | 1.00 | 0.97 | 0.89 |
| C93T | 0.58 | 0.78 | 0.71 | 0.79 | 0.99 | 0.99 | 0.81 |
| C93A | 0.48 | 0.80 | 0.83 | 0.80 | 0.98 | 1.00 | 0.83 |
| C94T | 0.59 | 0.78 | 0.73 | 0.80 | 0.99 | 0.97 | 0.83 |
| C94G | 0.52 | 0.75 | 0.82 | 0.79 | 0.97 | 0.97 | 0.82 |
| C94A | 0.68 | 0.84 | 0.73 | 0.86 | 1.00 | 1.00 | 0.87 |
| G95T | 0.60 | 0.81 | 0.71 | 0.82 | 1.00 | 1.00 | 0.83 |
| G95C | 0.85 | 0.84 | 0.76 | 0.93 | 0.96 | 0.98 | 0.91 |
| G95A | 0.58 | 0.81 | 0.69 | 0.80 | 1.00 | 1.00 | 0.82 |
| G95GA | 0.55 | 0.81 | 0.75 | 0.81 | 0.99 | 1.00 | 0.83 |
| GA96G | 0.44 | 0.66 | 0.83 | 0.71 | 0.95 | 0.90 | 0.78 |
| A96T | 0.60 | 0.75 | 0.68 | 0.78 | 0.99 | 0.98 | 0.80 |
| A96G | 0.58 | 0.81 | 0.72 | 0.81 | 1.00 | 1.00 | 0.82 |
| A96C | 0.64 | 0.82 | 0.69 | 0.83 | 1.00 | 1.00 | 0.84 |
| AG97A | 0.64 | 0.81 | 0.70 | 0.82 | 1.00 | 0.99 | 0.84 |
| G97T | 0.66 | 0.78 | 0.73 | 0.83 | 0.99 | 0.99 | 0.84 |
| G97C | 0.63 | 0.70 | 0.75 | 0.83 | 0.93 | 0.87 | 0.91 |
| G97A | 0.61 | 0.80 | 0.74 | 0.82 | 1.00 | 1.00 | 0.84 |
| G98T | 0.66 | 0.77 | 0.76 | 0.85 | 0.97 | 0.96 | 0.87 |
| G98C | 0.77 | 0.89 | 0.92 | 0.91 | 1.00 | 0.98 | 0.95 |
| G98A | 0.64 | 0.80 | 0.77 | 0.85 | 0.97 | 1.00 | 0.85 |
| G99T | 0.62 | 0.82 | 0.78 | 0.83 | 1.00 | 0.98 | 0.86 |
| G99C | 0.57 | 0.79 | 0.79 | 0.81 | 0.99 | 0.98 | 0.84 |
| G99A | 0.64 | 0.77 | 0.78 | 0.86 | 0.95 | 0.96 | 0.86 |

| | | | | | | | |
|---|---|---|---|---|---|---|---|
| G100T | 0.56 | 0.81 | 0.70 | 0.79 | 1.00 | 1.00 | 0.81 |
| G100C | 0.61 | 0.78 | 0.71 | 0.80 | 1.00 | 0.99 | 0.82 |
| G100A | 0.66 | 0.82 | 0.68 | 0.83 | 1.00 | 1.00 | 0.84 |
| A101T | 0.59 | 0.77 | 0.78 | 0.81 | 0.99 | 0.98 | 0.84 |
| A101G | 0.59 | 0.78 | 0.73 | 0.80 | 0.99 | 0.99 | 0.82 |
| T102C | 0.57 | 0.81 | 0.77 | 0.81 | 1.00 | 0.99 | 0.84 |
| T102A | 0.55 | 0.79 | 0.78 | 0.81 | 0.99 | 0.99 | 0.83 |
| TG103T | 0.68 | 0.85 | 0.77 | 0.87 | 0.99 | 1.00 | 0.88 |
| G103T | 0.67 | 0.77 | 0.59 | 0.79 | 0.99 | 1.00 | 0.78 |
| G103A | 0.58 | 0.84 | 0.66 | 0.80 | 1.00 | 0.99 | 0.82 |
| G104A | 0.59 | 0.80 | 0.73 | 0.84 | 0.96 | 1.00 | 0.83 |
| GA105G | 0.58 | 0.85 | 0.67 | 0.81 | 0.99 | 1.00 | 0.82 |
| A105T | 0.63 | 0.81 | 0.71 | 0.83 | 0.99 | 0.99 | 0.84 |
| A105G | 0.57 | 0.82 | 0.69 | 0.80 | 1.00 | 0.99 | 0.82 |
| A105C | 0.61 | 0.76 | 0.53 | 0.73 | 1.00 | 0.97 | 0.75 |
| G106T | 0.65 | 0.78 | 0.77 | 0.83 | 1.00 | 0.98 | 0.86 |
| G106C | 0.53 | 0.73 | 0.97 | 0.98 | 0.82 | 0.88 | 0.95 |
| G106A | 0.59 | 0.79 | 0.73 | 0.80 | 1.00 | 0.98 | 0.84 |
| C107T | 0.64 | 0.79 | 0.65 | 0.81 | 0.99 | 1.00 | 0.81 |
| C107G | 0.51 | 0.83 | 0.81 | 0.80 | 1.00 | 0.97 | 0.86 |
| C107A | 0.60 | 0.78 | 0.74 | 0.80 | 1.00 | 1.00 | 0.82 |
| T108G | 0.67 | 0.83 | 0.71 | 0.84 | 1.00 | 1.00 | 0.85 |
| T108C | 0.62 | 0.84 | 0.74 | 0.84 | 1.00 | 0.99 | 0.86 |
| T108A | 0.62 | 0.79 | 0.69 | 0.82 | 0.98 | 0.99 | 0.82 |
| G109T | 0.55 | 0.74 | 0.67 | 0.75 | 0.99 | 1.00 | 0.77 |
| G109C | 0.64 | 0.77 | 0.74 | 0.83 | 0.98 | 0.97 | 0.85 |
| G109A | 0.56 | 0.77 | 0.78 | 0.81 | 0.98 | 0.99 | 0.82 |
| C110T | 0.62 | 0.87 | 0.76 | 0.91 | 0.93 | 1.00 | 0.87 |
| C110A | 0.59 | 0.84 | 0.66 | 0.81 | 0.99 | 1.00 | 0.82 |
| T111G | 0.59 | 0.79 | 0.81 | 0.82 | 0.99 | 0.98 | 0.85 |
| T111C | 0.57 | 0.81 | 0.73 | 0.81 | 1.00 | 0.99 | 0.83 |
| T111A | 0.60 | 0.82 | 0.75 | 0.82 | 0.99 | 1.00 | 0.84 |
| TG112T | 0.61 | 0.78 | 0.77 | 0.83 | 0.97 | 1.00 | 0.83 |
| G112T | 0.65 | 0.79 | 0.65 | 0.80 | 1.00 | 0.99 | 0.82 |
| G112C | 0.71 | 0.85 | 0.80 | 0.88 | 1.00 | 0.99 | 0.89 |
| G112A | 0.55 | 0.76 | 0.86 | 0.82 | 0.98 | 0.95 | 0.87 |
| G113T | 0.65 | 0.75 | 0.76 | 0.84 | 0.96 | 0.94 | 0.87 |
| G113C | 0.78 | 0.78 | 0.75 | 0.94 | 0.91 | 0.90 | 0.95 |
| G113A | 0.62 | 0.80 | 0.79 | 0.83 | 0.99 | 0.99 | 0.86 |
| C114T | 0.68 | 0.81 | 0.70 | 0.86 | 0.97 | 1.00 | 0.84 |
| C114G | 0.61 | 0.79 | 0.81 | 0.83 | 0.99 | 0.98 | 0.86 |
| C114A | 0.73 | 0.77 | 0.79 | 0.97 | 0.87 | 0.88 | 0.96 |
| CT115C | 0.66 | 0.83 | 0.68 | 0.83 | 1.00 | 0.99 | 0.85 |
| T115G | 0.75 | 0.63 | 0.63 | 0.92 | 0.81 | 0.83 | 0.89 |
| T115C | 0.62 | 0.82 | 0.78 | 0.84 | 1.00 | 0.99 | 0.86 |
| T115A | 0.57 | 0.80 | 0.75 | 0.81 | 0.99 | 1.00 | 0.82 |
| T116C | 0.58 | 0.82 | 0.74 | 0.82 | 0.99 | 1.00 | 0.83 |
| T116A | 0.64 | 0.81 | 0.71 | 0.83 | 1.00 | 1.00 | 0.84 |
| T117C | 0.64 | 0.77 | 0.76 | 0.85 | 0.96 | 1.00 | 0.83 |
| T117A | 0.59 | 0.72 | 0.75 | 0.79 | 0.98 | 1.00 | 0.79 |
| TA118T | 0.57 | 0.84 | 0.76 | 0.83 | 0.98 | 1.00 | 0.85 |
| A118T | 0.51 | 0.79 | 0.81 | 0.79 | 0.99 | 0.99 | 0.82 |
| A118G | 0.65 | 0.82 | 0.73 | 0.84 | 1.00 | 0.99 | 0.85 |

| | | | | | | | |
|---|---|---|---|---|---|---|---|
| A118C | 0.66 | 0.74 | 0.74 | 0.86 | 0.93 | 0.95 | 0.86 |
| C119T | 0.60 | 0.82 | 0.70 | 0.81 | 1.00 | 0.96 | 0.86 |
| C119G | 0.52 | 0.75 | 0.84 | 0.81 | 0.96 | 0.98 | 0.82 |
| C119A | 0.56 | 0.81 | 0.54 | 0.74 | 0.99 | 1.00 | 0.73 |
| A120T | 0.59 | 0.80 | 0.70 | 0.80 | 0.99 | 0.99 | 0.82 |
| A120G | 0.57 | 0.81 | 0.72 | 0.81 | 0.99 | 1.00 | 0.82 |
| A120C | 0.52 | 0.81 | 0.89 | 0.85 | 0.96 | 1.00 | 0.86 |
| C121T | 0.56 | 0.78 | 0.74 | 0.79 | 0.99 | 0.99 | 0.82 |
| C121G | 0.48 | 0.81 | 0.87 | 0.80 | 0.99 | 1.00 | 0.83 |
| C121A | 0.56 | 0.79 | 0.71 | 0.79 | 0.99 | 0.98 | 0.81 |
| T122G | 0.62 | 0.82 | 0.77 | 0.84 | 1.00 | 0.99 | 0.86 |
| T122C | 0.62 | 0.77 | 0.80 | 0.83 | 0.98 | 0.97 | 0.86 |
| T122A | 0.64 | 0.78 | 0.71 | 0.82 | 0.99 | 0.98 | 0.85 |
| G123T | 0.62 | 0.81 | 0.77 | 0.84 | 0.99 | 0.99 | 0.86 |
| G123C | 0.58 | 0.83 | 0.71 | 0.81 | 1.00 | 1.00 | 0.82 |
| G123A | 0.58 | 0.82 | 0.69 | 0.80 | 1.00 | 1.00 | 0.82 |
| C124T | 0.60 | 0.83 | 0.78 | 0.87 | 0.95 | 1.00 | 0.85 |
| C124G | 0.53 | 0.75 | 0.81 | 0.80 | 0.97 | 0.99 | 0.81 |
| C124A | 0.64 | 0.79 | 0.69 | 0.82 | 0.99 | 0.99 | 0.83 |
| C125T | 0.59 | 0.77 | 0.79 | 0.82 | 0.99 | 0.98 | 0.84 |
| C125A | 0.62 | 0.82 | 0.70 | 0.83 | 0.99 | 1.00 | 0.83 |
| T126G | 0.83 | 0.75 | 0.56 | 0.85 | 0.94 | 0.92 | 0.85 |
| T126C | 0.62 | 0.76 | 0.70 | 0.80 | 0.99 | 0.99 | 0.81 |
| T126A | 0.46 | 0.80 | 0.75 | 0.75 | 0.99 | 0.99 | 0.79 |
| G127T | 0.57 | 0.78 | 0.70 | 0.79 | 0.99 | 0.99 | 0.81 |
| G127A | 0.59 | 0.88 | 0.86 | 0.88 | 0.97 | 1.00 | 0.89 |
| G128T | 0.62 | 0.87 | 0.68 | 0.87 | 0.97 | 1.00 | 0.85 |
| G128C | 0.67 | 0.81 | 0.78 | 0.86 | 0.98 | 1.00 | 0.87 |
| G128A | 0.65 | 0.84 | 0.71 | 0.84 | 1.00 | 0.99 | 0.86 |
| C129T | 0.59 | 0.82 | 0.76 | 0.82 | 0.99 | 0.99 | 0.84 |
| C129G | 0.64 | 0.74 | 0.82 | 0.91 | 0.89 | 0.92 | 0.91 |
| C129A | 0.63 | 0.76 | 0.77 | 0.83 | 0.98 | 0.97 | 0.85 |
| CT130C | 0.69 | 0.90 | 0.81 | 0.89 | 1.00 | 0.98 | 0.92 |
| T130G | 0.65 | 0.75 | 0.77 | 0.89 | 0.91 | 0.99 | 0.83 |
| T130C | 0.64 | 0.75 | 0.81 | 0.89 | 0.91 | 0.93 | 0.89 |
| T130A | 0.56 | 0.80 | 0.72 | 0.79 | 1.00 | 1.00 | 0.82 |
| T131G | 0.44 | 0.79 | 0.82 | 0.76 | 0.99 | 0.99 | 0.79 |
| T131C | 0.59 | 0.77 | 0.67 | 0.78 | 0.99 | 0.99 | 0.79 |
| T131A | 0.61 | 0.75 | 0.72 | 0.78 | 1.00 | 0.94 | 0.85 |
| T132C | 0.62 | 0.79 | 0.74 | 0.82 | 1.00 | 0.99 | 0.84 |
| T132A | 0.58 | 0.84 | 0.67 | 0.80 | 0.99 | 1.00 | 0.81 |
| TC133T | 0.75 | 0.89 | 0.65 | 0.88 | 1.00 | 1.00 | 0.87 |
| C133T | 0.58 | 0.79 | 0.69 | 0.79 | 0.99 | 1.00 | 0.80 |
| C133G | 0.67 | 0.65 | 0.97 | 1.00 | 0.79 | 0.81 | 1.00 |
| C133A | 0.64 | 0.74 | 0.85 | 0.96 | 0.85 | 0.88 | 0.95 |
| G134T | 0.66 | 0.86 | 0.68 | 0.85 | 1.00 | 1.00 | 0.86 |
| G134C | 0.64 | 0.82 | 0.69 | 0.83 | 0.99 | 1.00 | 0.83 |
| G134A | 0.59 | 0.80 | 0.72 | 0.80 | 1.00 | 0.99 | 0.83 |
| C135T | 0.59 | 0.72 | 0.71 | 0.78 | 0.98 | 0.94 | 0.82 |
| C135G | 0.36 | 0.43 | 1.00 | 0.82 | 0.20 | 0.32 | 0.72 |
| C135A | 0.52 | 0.92 | 0.95 | 0.86 | 0.99 | 0.99 | 0.92 |
| T136C | 0.59 | 0.78 | 0.80 | 0.81 | 1.00 | 0.98 | 0.85 |
| T136A | 0.55 | 0.78 | 0.72 | 0.78 | 1.00 | 0.99 | 0.80 |

| | | | | | | | |
|---|---|---|---|---|---|---|---|
| T137G | 0.62 | 0.78 | 0.76 | 0.81 | 1.00 | 0.97 | 0.85 |
| T137C | 0.60 | 0.80 | 0.70 | 0.80 | 1.00 | 1.00 | 0.82 |
| T137A | 0.58 | 0.82 | 0.75 | 0.82 | 1.00 | 0.99 | 0.84 |
| TC138T | 0.63 | 0.78 | 0.68 | 0.81 | 0.98 | 0.99 | 0.81 |
| C138T | 0.58 | 0.77 | 0.69 | 0.78 | 1.00 | 0.93 | 0.85 |
| C139T | 0.56 | 0.78 | 0.70 | 0.78 | 0.99 | 1.00 | 0.80 |
| C139A | 0.58 | 0.79 | 0.72 | 0.80 | 0.99 | 0.98 | 0.83 |
| T140C | 0.60 | 0.78 | 0.73 | 0.81 | 0.99 | 1.00 | 0.82 |
| T140A | 0.59 | 0.78 | 0.74 | 0.80 | 0.99 | 0.99 | 0.82 |
| A141T | 0.56 | 0.80 | 0.74 | 0.80 | 0.99 | 1.00 | 0.82 |
| A141G | 0.61 | 0.79 | 0.77 | 0.82 | 0.99 | 0.98 | 0.85 |
| A141C | 0.60 | 0.73 | 0.79 | 0.82 | 0.96 | 0.93 | 0.87 |
| A141AC | 0.56 | 0.81 | 0.73 | 0.80 | 1.00 | 0.99 | 0.83 |
| AC142A | 0.65 | 0.81 | 0.70 | 0.83 | 1.00 | 1.00 | 0.84 |
| C142T | 0.60 | 0.80 | 0.73 | 0.82 | 0.99 | 1.00 | 0.82 |
| C142G | 0.57 | 0.77 | 0.78 | 0.79 | 1.00 | 0.98 | 0.83 |
| C142A | 0.64 | 0.67 | 0.77 | 0.90 | 0.85 | 0.84 | 0.93 |
| C143T | 0.62 | 0.73 | 0.64 | 0.77 | 0.99 | 0.99 | 0.78 |
| C143G | 0.55 | 0.76 | 0.76 | 0.79 | 0.98 | 0.98 | 0.81 |
| C143A | 0.65 | 0.77 | 0.74 | 0.83 | 0.98 | 1.00 | 0.83 |
| C144T | 0.61 | 0.74 | 0.78 | 0.80 | 0.99 | 0.97 | 0.84 |
| C144G | 0.81 | 0.82 | 0.61 | 0.86 | 0.99 | 1.00 | 0.84 |
| C144A | 0.70 | 0.82 | 0.80 | 0.87 | 0.98 | 0.99 | 0.87 |
| C145T | 0.63 | 0.80 | 0.68 | 0.81 | 1.00 | 0.99 | 0.83 |
| C145G | 0.62 | 0.77 | 0.75 | 0.82 | 0.98 | 0.98 | 0.83 |
| C145A | 0.67 | 0.77 | 0.62 | 0.80 | 1.00 | 0.99 | 0.80 |
| C146T | 0.62 | 0.82 | 0.72 | 0.82 | 1.00 | 1.00 | 0.84 |
| C146A | 0.60 | 0.84 | 0.75 | 0.83 | 1.00 | 0.99 | 0.86 |
| CA147C | 0.62 | 0.62 | 0.66 | 0.79 | 0.89 | 0.85 | 0.84 |
| A147T | 0.60 | 0.81 | 0.73 | 0.82 | 0.99 | 1.00 | 0.83 |
| A147G | 0.59 | 0.80 | 0.76 | 0.82 | 0.99 | 0.99 | 0.84 |
| A147C | 0.62 | 0.80 | 0.74 | 0.82 | 1.00 | 0.98 | 0.85 |
| A147AC | 0.58 | 0.83 | 0.77 | 0.82 | 1.00 | 0.99 | 0.85 |
| AC148A | 0.64 | 0.80 | 0.75 | 0.83 | 1.00 | 1.00 | 0.84 |
| C148T | 0.66 | 0.79 | 0.74 | 0.84 | 0.98 | 0.99 | 0.85 |
| C148G | 0.62 | 0.82 | 0.74 | 0.83 | 0.99 | 0.99 | 0.85 |
| C148A | 0.73 | 0.88 | 0.70 | 0.89 | 1.00 | 1.00 | 0.89 |
| C149T | 0.62 | 0.79 | 0.76 | 0.82 | 1.00 | 0.99 | 0.84 |
| C149G | 0.49 | 0.79 | 0.80 | 0.77 | 1.00 | 0.99 | 0.81 |
| C149A | 0.58 | 0.76 | 0.71 | 0.79 | 0.99 | 0.98 | 0.81 |
| C150T | 0.60 | 0.74 | 0.79 | 0.89 | 0.89 | 0.98 | 0.83 |
| C150G | 0.78 | 0.79 | 0.61 | 0.85 | 0.98 | 0.99 | 0.83 |
| C150A | 0.65 | 0.78 | 0.73 | 0.88 | 0.93 | 1.00 | 0.83 |
| C151T | 0.55 | 0.74 | 0.77 | 0.78 | 0.98 | 0.98 | 0.80 |
| C151CT | 0.58 | 0.87 | 0.64 | 0.81 | 1.00 | 0.96 | 0.86 |
| A152T | 0.59 | 0.77 | 0.73 | 0.80 | 0.99 | 0.99 | 0.82 |
| A152G | 0.58 | 0.78 | 0.76 | 0.80 | 1.00 | 0.99 | 0.83 |
| A152C | 0.68 | 0.81 | 0.81 | 0.86 | 0.99 | 0.99 | 0.88 |
| AT153A | 0.68 | 0.86 | 0.76 | 0.86 | 1.00 | 0.99 | 0.88 |
| T153C | 0.61 | 0.77 | 0.70 | 0.80 | 0.99 | 0.99 | 0.81 |
| T153A | 0.61 | 0.75 | 0.78 | 0.81 | 0.99 | 0.99 | 0.83 |
| C154T | 0.58 | 0.83 | 0.70 | 0.81 | 0.99 | 1.00 | 0.83 |
| C154G | 0.59 | 0.75 | 0.68 | 0.78 | 0.98 | 0.98 | 0.79 |

| | | | | | | | |
|---|---|---|---|---|---|---|---|
| C154A | 0.56 | 0.81 | 0.72 | 0.80 | 1.00 | 0.99 | 0.82 |
| C155T | 0.57 | 0.81 | 0.72 | 0.81 | 0.99 | 0.99 | 0.83 |
| C155G | 0.62 | 0.82 | 0.75 | 0.83 | 1.00 | 0.99 | 0.85 |
| C155A | 0.61 | 0.76 | 0.73 | 0.80 | 0.99 | 0.98 | 0.83 |
| CA156C | 0.60 | 0.86 | 0.67 | 0.82 | 1.00 | 1.00 | 0.84 |
| A156T | 0.60 | 0.82 | 0.70 | 0.81 | 1.00 | 0.99 | 0.83 |
| A156G | 0.58 | 0.78 | 0.73 | 0.79 | 1.00 | 0.99 | 0.82 |
| A156C | 0.69 | 0.81 | 0.78 | 0.87 | 0.98 | 0.98 | 0.88 |
| AC157A | 0.60 | 0.81 | 0.71 | 0.81 | 1.00 | 0.99 | 0.83 |
| C157T | 0.65 | 0.82 | 0.73 | 0.85 | 0.98 | 1.00 | 0.85 |
| C157G | 0.62 | 0.81 | 0.67 | 0.81 | 1.00 | 0.99 | 0.82 |
| C157A | 0.62 | 0.73 | 0.56 | 0.71 | 1.00 | 0.90 | 0.79 |
| C158T | 0.57 | 0.79 | 0.72 | 0.80 | 1.00 | 1.00 | 0.81 |
| C158G | 0.56 | 0.76 | 0.71 | 0.77 | 0.99 | 0.98 | 0.79 |
| C158A | 0.62 | 0.77 | 0.71 | 0.82 | 0.97 | 0.99 | 0.82 |
| C159T | 0.61 | 0.80 | 0.64 | 0.80 | 0.99 | 1.00 | 0.80 |
| C159G | 0.67 | 0.81 | 0.57 | 0.80 | 1.00 | 0.99 | 0.80 |
| C159A | 0.72 | 0.92 | 0.76 | 0.89 | 1.00 | 1.00 | 0.91 |
| CA160C | 0.61 | 0.86 | 0.77 | 0.85 | 1.00 | 0.99 | 0.87 |
| A160T | 0.67 | 0.77 | 0.71 | 0.85 | 0.96 | 1.00 | 0.82 |
| A160G | 0.70 | 0.75 | 0.75 | 0.90 | 0.92 | 0.94 | 0.88 |
| A160C | 0.60 | 0.84 | 0.71 | 0.83 | 1.00 | 0.99 | 0.85 |
| G161T | 0.56 | 0.83 | 0.72 | 0.81 | 1.00 | 0.98 | 0.84 |
| G161C | 0.50 | 0.75 | 0.84 | 0.79 | 0.97 | 0.98 | 0.81 |
| G161A | 0.54 | 0.79 | 0.77 | 0.79 | 0.99 | 0.99 | 0.82 |
| T162G | 0.60 | 0.84 | 0.66 | 0.82 | 0.99 | 1.00 | 0.82 |
| T162C | 0.64 | 0.80 | 0.73 | 0.82 | 1.00 | 0.99 | 0.84 |
| T162A | 0.60 | 0.79 | 0.73 | 0.82 | 0.98 | 0.99 | 0.83 |
| G163T | 0.52 | 0.84 | 0.67 | 0.78 | 1.00 | 1.00 | 0.80 |
| G163C | 0.60 | 0.69 | 0.67 | 0.77 | 0.95 | 0.97 | 0.77 |
| G163A | 0.59 | 0.80 | 0.75 | 0.81 | 1.00 | 1.00 | 0.84 |
| C164T | 0.60 | 0.79 | 0.73 | 0.82 | 0.98 | 1.00 | 0.82 |
| C164G | 0.55 | 0.92 | 0.73 | 0.84 | 1.00 | 1.00 | 0.86 |
| C164A | 0.78 | 0.78 | 0.71 | 0.89 | 0.96 | 0.94 | 0.90 |
| C165T | 0.54 | 0.76 | 0.73 | 0.77 | 0.99 | 1.00 | 0.79 |
| C165G | 0.77 | 0.88 | 0.57 | 0.87 | 1.00 | 1.00 | 0.85 |
| C165A | 0.66 | 0.77 | 0.92 | 0.99 | 0.85 | 0.87 | 0.99 |
| CA166C | 0.47 | 0.76 | 1.00 | 0.90 | 0.87 | 0.91 | 0.93 |
| A166T | 0.66 | 0.80 | 0.68 | 0.83 | 0.99 | 1.00 | 0.83 |
| A166G | 0.64 | 0.79 | 0.74 | 0.83 | 0.99 | 0.99 | 0.84 |
| A166C | 0.58 | 0.82 | 0.70 | 0.80 | 1.00 | 0.99 | 0.82 |
| A167T | 0.59 | 0.75 | 0.78 | 0.83 | 0.95 | 0.99 | 0.81 |
| A167G | 0.58 | 0.83 | 0.65 | 0.80 | 1.00 | 1.00 | 0.81 |
| AC168A | 0.46 | 0.72 | 0.76 | 0.72 | 0.98 | 0.98 | 0.75 |
| C168T | 0.63 | 0.82 | 0.66 | 0.82 | 1.00 | 1.00 | 0.83 |
| C168G | 0.65 | 0.82 | 0.68 | 0.83 | 1.00 | 1.00 | 0.83 |
| C168A | 0.61 | 0.85 | 0.69 | 0.90 | 0.93 | 1.00 | 0.85 |
| C169T | 0.62 | 0.82 | 0.69 | 0.82 | 1.00 | 1.00 | 0.83 |
| C169G | 0.55 | 0.72 | 0.83 | 0.79 | 0.98 | 0.93 | 0.86 |
| C169A | 0.64 | 0.80 | 0.76 | 0.84 | 0.99 | 1.00 | 0.85 |
| T170G | 0.71 | 0.79 | 0.79 | 0.91 | 0.93 | 0.98 | 0.88 |
| T170C | 0.61 | 0.77 | 0.73 | 0.82 | 0.97 | 0.99 | 0.82 |
| T170A | 0.61 | 0.79 | 0.71 | 0.81 | 1.00 | 0.97 | 0.84 |

| | | | | | | | |
|---|---|---|---|---|---|---|---|
| A171T | 0.65 | 0.78 | 0.74 | 0.82 | 1.00 | 0.99 | 0.84 |
| A171G | 0.60 | 0.76 | 0.84 | 0.85 | 0.96 | 0.95 | 0.88 |
| A171C | 0.71 | 0.64 | 0.77 | 0.99 | 0.77 | 0.78 | 0.98 |
| G172T | 0.59 | 0.78 | 0.93 | 0.95 | 0.87 | 0.92 | 0.94 |
| G172A | 0.61 | 0.77 | 0.75 | 0.83 | 0.97 | 0.98 | 0.83 |
| GT173G | 0.90 | 0.94 | 0.93 | 0.97 | 0.98 | 1.00 | 0.96 |
| T173G | 0.61 | 0.82 | 0.71 | 0.82 | 1.00 | 0.99 | 0.84 |
| T173C | 0.62 | 0.80 | 0.71 | 0.81 | 1.00 | 0.99 | 0.83 |
| T173A | 0.60 | 0.79 | 0.79 | 0.82 | 0.99 | 0.99 | 0.85 |
| T174G | 0.57 | 0.67 | 0.52 | 0.63 | 0.78 | 0.69 | 0.71 |
| T174C | 0.55 | 0.77 | 0.70 | 0.76 | 1.00 | 0.94 | 0.83 |
| T174A | 0.65 | 0.82 | 0.75 | 0.84 | 1.00 | 0.99 | 0.86 |
| C175T | 0.60 | 0.82 | 0.63 | 0.79 | 1.00 | 1.00 | 0.80 |
| C175G | 0.52 | 0.75 | 0.87 | 0.82 | 0.95 | 0.95 | 0.86 |
| C175A | 0.69 | 0.97 | 0.72 | 0.90 | 1.00 | 1.00 | 0.91 |
| C176T | 0.69 | 0.79 | 0.70 | 0.84 | 0.99 | 0.97 | 0.86 |
| C176A | 0.67 | 0.80 | 0.63 | 0.81 | 1.00 | 0.99 | 0.82 |
| A177T | 0.55 | 0.81 | 0.64 | 0.77 | 0.99 | 1.00 | 0.79 |
| A177G | 0.65 | 0.80 | 0.63 | 0.81 | 0.99 | 1.00 | 0.81 |
| A177C | 0.62 | 0.63 | 0.73 | 0.89 | 0.82 | 0.83 | 0.88 |
| C178T | 0.71 | 0.78 | 0.74 | 0.86 | 0.98 | 0.98 | 0.87 |
| C178G | 0.73 | 0.82 | 0.82 | 0.88 | 0.99 | 0.97 | 0.91 |
| C178A | 0.58 | 0.83 | 0.70 | 0.81 | 1.00 | 0.99 | 0.84 |
| T179G | 0.62 | 0.74 | 0.77 | 0.81 | 0.99 | 0.96 | 0.85 |
| T179C | 0.60 | 0.80 | 0.77 | 0.82 | 1.00 | 0.94 | 0.89 |
| T179A | 0.57 | 0.83 | 0.73 | 0.81 | 1.00 | 1.00 | 0.83 |
| G180T | 0.48 | 0.82 | 0.68 | 0.74 | 1.00 | 0.99 | 0.78 |
| G180A | 0.48 | 0.75 | 0.69 | 0.73 | 0.98 | 1.00 | 0.73 |
| A181T | 0.55 | 0.79 | 0.65 | 0.76 | 1.00 | 0.96 | 0.81 |
| A181G | 0.68 | 0.79 | 0.68 | 0.83 | 0.99 | 0.99 | 0.84 |
| A181C | 0.53 | 0.77 | 1.00 | 0.96 | 0.85 | 0.89 | 0.96 |
| A182T | 0.45 | 0.78 | 0.92 | 0.79 | 0.99 | 0.99 | 0.84 |
| A182G | 0.59 | 0.84 | 0.67 | 0.81 | 0.99 | 0.99 | 0.83 |
| A182C | 0.57 | 0.84 | 0.52 | 0.75 | 0.99 | 0.98 | 0.75 |
| G183T | 0.53 | 0.85 | 0.89 | 0.84 | 0.99 | 0.99 | 0.87 |
| G183C | 0.48 | 0.88 | 0.63 | 0.75 | 1.00 | 0.99 | 0.78 |
| G183A | 0.58 | 0.77 | 0.72 | 0.80 | 0.98 | 1.00 | 0.80 |
| C184T | 0.67 | 0.80 | 0.88 | 0.96 | 0.89 | 0.95 | 0.92 |
| C184A | 0.94 | 0.70 | 0.70 | 1.00 | 0.85 | 0.83 | 1.00 |
| C185T | 0.77 | 0.77 | 0.79 | 0.95 | 0.91 | 0.89 | 0.97 |
| C185A | 0.70 | 0.72 | 0.66 | 0.82 | 0.98 | 0.97 | 0.82 |
| T186G | 0.35 | 0.96 | 0.84 | 0.77 | 1.00 | 1.00 | 0.82 |
| T186C | 0.62 | 0.78 | 0.73 | 0.82 | 0.99 | 0.99 | 0.83 |
| T186A | 0.63 | 0.80 | 0.71 | 0.82 | 0.99 | 0.98 | 0.84 |
| G187T | 0.69 | 0.85 | 0.58 | 0.84 | 1.00 | 0.99 | 0.84 |
| G187A | 0.50 | 0.72 | 0.84 | 0.78 | 0.97 | 0.95 | 0.82 |
| A188T | 0.55 | 0.80 | 0.72 | 0.83 | 0.94 | 1.00 | 0.81 |
| A188G | 0.59 | 0.81 | 0.66 | 0.80 | 1.00 | 1.00 | 0.82 |
| A188C | 0.62 | 0.80 | 0.72 | 0.83 | 0.98 | 1.00 | 0.83 |
| AG189A | 0.58 | 0.82 | 0.62 | 0.79 | 1.00 | 1.00 | 0.80 |
| G189T | 0.56 | 0.85 | 0.66 | 0.80 | 1.00 | 1.00 | 0.82 |
| G189C | 0.60 | 0.80 | 0.67 | 0.80 | 0.99 | 1.00 | 0.81 |
| G189A | 0.55 | 0.78 | 0.80 | 0.80 | 0.99 | 0.99 | 0.83 |

| | | | | | | | |
|---|---|---|---|---|---|---|---|
| G190T | 0.53 | 0.86 | 0.65 | 0.78 | 1.00 | 0.97 | 0.83 |
| G190A | 0.64 | 0.84 | 0.80 | 0.85 | 1.00 | 0.99 | 0.88 |
| A191T | 0.52 | 0.80 | 0.71 | 0.78 | 0.99 | 0.99 | 0.80 |
| A191G | 0.56 | 0.84 | 0.65 | 0.79 | 0.99 | 0.99 | 0.81 |
| A191C | 0.57 | 0.87 | 0.65 | 0.81 | 1.00 | 1.00 | 0.82 |
| G192T | 0.66 | 0.84 | 0.75 | 0.85 | 1.00 | 0.99 | 0.87 |
| G192C | 0.62 | 0.79 | 0.65 | 0.80 | 1.00 | 1.00 | 0.80 |
| G192A | 0.59 | 0.76 | 0.75 | 0.80 | 0.99 | 0.99 | 0.81 |
| C193T | 0.66 | 0.81 | 0.71 | 0.84 | 0.99 | 1.00 | 0.84 |
| C193G | 0.58 | 0.83 | 0.73 | 0.82 | 0.99 | 1.00 | 0.84 |
| C193A | 0.65 | 0.82 | 0.64 | 0.85 | 0.96 | 1.00 | 0.82 |
| A194T | 0.59 | 0.77 | 0.69 | 0.78 | 1.00 | 0.97 | 0.81 |
| A194G | 0.60 | 0.81 | 0.73 | 0.82 | 0.99 | 0.99 | 0.83 |
| T195G | 0.61 | 0.81 | 0.68 | 0.81 | 1.00 | 1.00 | 0.83 |
| T195C | 0.59 | 0.79 | 0.75 | 0.82 | 0.99 | 0.99 | 0.83 |
| T195A | 0.65 | 0.78 | 0.77 | 0.83 | 0.99 | 1.00 | 0.84 |
| G196T | 0.61 | 0.79 | 0.76 | 0.84 | 0.97 | 1.00 | 0.84 |
| G196C | 0.63 | 0.84 | 0.69 | 0.83 | 0.99 | 1.00 | 0.84 |
| G196A | 0.63 | 0.82 | 0.74 | 0.83 | 1.00 | 0.98 | 0.86 |
| C197T | 0.52 | 0.79 | 0.80 | 0.79 | 0.99 | 0.99 | 0.82 |
| C197G | 0.45 | 0.82 | 0.80 | 0.77 | 0.99 | 1.00 | 0.80 |
| C197A | 0.63 | 0.79 | 0.69 | 0.82 | 0.98 | 0.99 | 0.83 |
| C198T | 0.56 | 0.78 | 0.70 | 0.78 | 0.99 | 1.00 | 0.80 |
| C198G | 0.75 | 0.80 | 0.84 | 0.98 | 0.89 | 0.93 | 0.95 |
| C198A | 0.54 | 0.73 | 0.73 | 0.77 | 0.97 | 0.98 | 0.79 |
| A199T | 0.61 | 0.83 | 0.72 | 0.82 | 1.00 | 1.00 | 0.84 |
| A199G | 0.65 | 0.81 | 0.66 | 0.82 | 1.00 | 1.00 | 0.82 |
| A199C | 0.61 | 0.83 | 0.75 | 0.83 | 0.99 | 1.00 | 0.85 |
| AT200A | 0.41 | 0.87 | 0.97 | 0.81 | 0.99 | 0.99 | 0.87 |
| T200C | 0.65 | 0.81 | 0.74 | 0.83 | 1.00 | 0.99 | 0.85 |
| T200A | 0.63 | 0.79 | 0.71 | 0.82 | 0.99 | 0.99 | 0.83 |
| T201G | 0.68 | 0.72 | 0.85 | 0.99 | 0.83 | 0.84 | 0.99 |
| T201C | 0.53 | 0.78 | 0.88 | 0.82 | 0.98 | 0.97 | 0.86 |
| T201A | 0.63 | 0.79 | 0.79 | 0.83 | 0.99 | 0.98 | 0.86 |
| A202T | 0.65 | 0.79 | 0.74 | 0.83 | 0.99 | 1.00 | 0.84 |
| A202G | 0.60 | 0.77 | 0.74 | 0.81 | 0.98 | 0.98 | 0.83 |
| A202C | 0.63 | 0.74 | 0.72 | 0.83 | 0.96 | 0.99 | 0.82 |
| A203T | 0.54 | 0.78 | 0.83 | 0.82 | 0.97 | 0.98 | 0.84 |
| A203G | 0.57 | 0.78 | 0.63 | 0.76 | 1.00 | 0.99 | 0.78 |
| G204T | 0.55 | 0.93 | 0.64 | 0.83 | 1.00 | 1.00 | 0.85 |
| G204C | 0.52 | 0.82 | 0.81 | 0.80 | 1.00 | 1.00 | 0.83 |
| G204A | 0.62 | 0.77 | 0.93 | 0.98 | 0.85 | 0.89 | 0.96 |
| T205G | 0.68 | 0.76 | 0.86 | 0.97 | 0.87 | 0.88 | 0.98 |
| T205C | 0.62 | 0.82 | 0.79 | 0.84 | 0.99 | 0.98 | 0.87 |
| T205A | 0.66 | 0.80 | 0.68 | 0.83 | 1.00 | 0.99 | 0.84 |
| T206G | 0.57 | 0.80 | 0.65 | 0.78 | 1.00 | 1.00 | 0.79 |
| T206C | 0.61 | 0.77 | 0.77 | 0.82 | 0.99 | 0.99 | 0.83 |
| T206A | 0.57 | 0.80 | 0.73 | 0.80 | 0.99 | 1.00 | 0.82 |
| T207G | 0.57 | 0.79 | 0.73 | 0.80 | 1.00 | 1.00 | 0.81 |
| T207C | 0.63 | 0.84 | 0.77 | 0.84 | 1.00 | 0.99 | 0.87 |
| T207A | 0.64 | 0.85 | 0.67 | 0.84 | 1.00 | 0.99 | 0.86 |
| G208T | 0.57 | 0.75 | 0.72 | 0.79 | 0.99 | 0.99 | 0.81 |
| G208A | 0.65 | 0.79 | 0.75 | 0.83 | 1.00 | 0.93 | 0.90 |

| | | | | | | | |
|---|---|---|---|---|---|---|---|
| A209T | 0.61 | 0.79 | 0.68 | 0.82 | 0.97 | 1.00 | 0.81 |
| A209G | 0.53 | 0.81 | 0.63 | 0.76 | 1.00 | 0.99 | 0.78 |
| A209C | 0.65 | 0.85 | 0.69 | 0.85 | 1.00 | 1.00 | 0.86 |
| G210T | 0.47 | 0.82 | 0.51 | 0.64 | 0.95 | 0.95 | 0.65 |
| G210C | 0.60 | 0.84 | 0.63 | 0.81 | 1.00 | 1.00 | 0.83 |
| G210A | 0.49 | 0.79 | 0.64 | 0.73 | 0.98 | 0.99 | 0.75 |
| G211T | 0.03 | 0.73 | 0.49 | 0.07 | 0.23 | 0.37 | 0.35 |
| T212C | 0.19 | 0.76 | 0.56 | 0.37 | 0.34 | 0.37 | 0.52 |
| T212A | 0.02 | 0.66 | 0.63 | 0.07 | 0.26 | 0.31 | 0.51 |
| A213T | 0.40 | 0.82 | 0.49 | 0.53 | 0.65 | 0.79 | 0.46 |
| A213G | 0.58 | 0.79 | 0.66 | 0.78 | 1.00 | 0.99 | 0.80 |
| A214T | 0.53 | 0.85 | 0.67 | 0.79 | 1.00 | 1.00 | 0.81 |
| A214G | 0.53 | 0.79 | 0.64 | 0.75 | 0.99 | 1.00 | 0.77 |
| A214C | 0.57 | 0.90 | 0.72 | 0.84 | 1.00 | 1.00 | 0.87 |
| G215C | 0.41 | 0.82 | 0.54 | 0.56 | 0.99 | 0.93 | 0.62 |
| G215A | 0.46 | 0.85 | 0.51 | 0.93 | 0.69 | 1.00 | 0.66 |
| T216G | 0.48 | 0.89 | 0.48 | 0.71 | 0.95 | 0.98 | 0.69 |
| T216C | 0.59 | 0.80 | 0.66 | 0.83 | 0.96 | 1.00 | 0.81 |
| T216A | 0.53 | 0.79 | 0.64 | 0.75 | 0.99 | 0.99 | 0.77 |
| G217T | 0.60 | 0.72 | 0.68 | 0.78 | 0.97 | 0.96 | 0.79 |
| G217C | 0.59 | 0.75 | 0.87 | 0.92 | 0.87 | 0.88 | 0.93 |
| G217A | 0.63 | 0.84 | 0.68 | 0.83 | 1.00 | 1.00 | 0.84 |
| GT218G | 1.00 | 0.59 | 0.92 | 1.00 | 0.81 | 0.80 | 1.00 |
| T218G | 0.52 | 0.93 | 0.74 | 0.83 | 1.00 | 1.00 | 0.88 |
| T218C | 0.54 | 0.80 | 0.73 | 0.78 | 1.00 | 0.99 | 0.81 |
| T218A | 0.65 | 0.78 | 0.70 | 0.82 | 1.00 | 0.99 | 0.83 |
| TA219T | 0.64 | 0.77 | 0.64 | 0.80 | 0.98 | 0.99 | 0.80 |
| A219T | 0.60 | 0.80 | 0.66 | 0.81 | 0.99 | 1.00 | 0.81 |
| A219G | 0.65 | 0.84 | 0.72 | 0.84 | 1.00 | 1.00 | 0.85 |
| A220T | 0.62 | 0.68 | 0.83 | 0.95 | 0.82 | 0.83 | 0.95 |
| A220G | 0.55 | 0.81 | 0.74 | 0.81 | 0.98 | 1.00 | 0.82 |
| A220C | 0.69 | 0.80 | 0.77 | 0.87 | 0.97 | 1.00 | 0.85 |
| AG221A | 0.63 | 0.66 | 0.69 | 0.81 | 0.92 | 0.85 | 0.89 |
| G221T | 0.62 | 0.73 | 0.70 | 0.78 | 0.99 | 0.99 | 0.79 |
| G221A | 0.57 | 0.79 | 0.60 | 0.76 | 1.00 | 0.97 | 0.79 |
| G222T | 0.57 | 0.83 | 0.67 | 0.80 | 1.00 | 0.99 | 0.81 |
| G222A | 0.50 | 0.83 | 0.64 | 0.76 | 1.00 | 1.00 | 0.78 |
| G223T | 0.71 | 0.77 | 0.81 | 0.94 | 0.90 | 0.91 | 0.94 |
| G223A | 0.56 | 0.76 | 0.70 | 0.77 | 1.00 | 0.98 | 0.80 |
| A224T | 0.59 | 0.73 | 0.74 | 0.79 | 0.98 | 0.97 | 0.82 |
| A224G | 0.57 | 0.73 | 0.86 | 0.88 | 0.89 | 0.90 | 0.90 |
| T225C | 0.65 | 0.77 | 0.80 | 0.84 | 0.98 | 0.95 | 0.89 |
| TA226T | 0.76 | 0.50 | 0.83 | 1.00 | 0.64 | 0.65 | 1.00 |
| A226T | 0.51 | 0.73 | 0.81 | 0.78 | 0.97 | 0.96 | 0.81 |
| A226G | 0.65 | 0.77 | 0.82 | 0.90 | 0.92 | 0.93 | 0.90 |
| A226C | 0.73 | 0.59 | 0.65 | 0.94 | 0.76 | 0.78 | 0.91 |
| A226AT | 0.47 | 0.85 | 0.74 | 0.77 | 1.00 | 1.00 | 0.81 |
| A226AG | 0.59 | 0.85 | 0.72 | 0.82 | 1.00 | 1.00 | 0.85 |
| AG227A | 0.59 | 0.79 | 0.50 | 0.73 | 0.98 | 0.99 | 0.72 |
| G227T | 0.52 | 0.78 | 0.80 | 0.78 | 1.00 | 0.99 | 0.81 |
| G227C | 0.50 | 0.34 | 0.51 | 0.64 | 0.16 | 0.16 | 0.65 |
| G227A | 0.47 | 0.73 | 0.81 | 0.75 | 0.98 | 0.98 | 0.78 |
| G228T | 0.65 | 0.59 | 0.53 | 0.68 | 0.45 | 0.39 | 0.78 |

| | | | | | | | |
|---|---|---|---|---|---|---|---|
| G228A | 0.49 | 0.62 | 0.74 | 0.75 | 0.80 | 0.74 | 0.84 |
| G228GT | 0.68 | 0.72 | 0.66 | 0.82 | 0.96 | 0.97 | 0.81 |
| G229T | 0.60 | 0.62 | 0.67 | 0.83 | 0.82 | 0.90 | 0.77 |
| G229A | 0.47 | 0.63 | 0.71 | 0.85 | 0.58 | 0.75 | 0.68 |
| G230T | 0.55 | 0.61 | 0.79 | 0.88 | 0.77 | 0.80 | 0.87 |
| G230C | 0.51 | 0.68 | 0.67 | 0.73 | 0.91 | 0.95 | 0.72 |
| G230A | 0.57 | 0.71 | 0.70 | 0.77 | 0.96 | 0.95 | 0.79 |
| C231T | 0.55 | 0.80 | 0.74 | 0.79 | 1.00 | 1.00 | 0.82 |
| C231A | 0.58 | 0.81 | 0.82 | 0.86 | 0.96 | 1.00 | 0.84 |
| CA232C | 0.54 | 0.83 | 0.96 | 0.86 | 0.98 | 0.98 | 0.89 |
| A232T | 0.59 | 0.77 | 0.79 | 0.83 | 0.97 | 0.99 | 0.83 |
| A232G | 0.59 | 0.83 | 0.74 | 0.82 | 0.99 | 1.00 | 0.84 |
| A232C | 0.62 | 0.71 | 0.77 | 0.88 | 0.89 | 0.90 | 0.88 |
| AG233A | 0.64 | 0.70 | 0.68 | 0.78 | 0.98 | 0.99 | 0.78 |
| G233T | 0.49 | 0.65 | 0.88 | 0.91 | 0.79 | 0.84 | 0.88 |
| G233C | 0.58 | 0.69 | 0.70 | 0.78 | 0.94 | 0.94 | 0.79 |
| G233A | 0.58 | 0.64 | 0.85 | 0.96 | 0.77 | 0.82 | 0.92 |
| G234T | 0.62 | 0.58 | 0.81 | 0.97 | 0.73 | 0.76 | 0.94 |
| G234C | 0.67 | 0.69 | 0.79 | 0.96 | 0.83 | 0.87 | 0.93 |
| G234A | 0.56 | 0.58 | 0.85 | 0.91 | 0.76 | 0.72 | 0.98 |
| G235T | 0.64 | 0.65 | 0.73 | 0.86 | 0.87 | 0.83 | 0.92 |
| G235A | 0.58 | 0.70 | 0.75 | 0.81 | 0.93 | 0.97 | 0.80 |
| A236T | 0.61 | 0.79 | 0.81 | 0.83 | 0.99 | 0.97 | 0.87 |
| A236G | 0.57 | 0.80 | 0.87 | 0.83 | 1.00 | 0.97 | 0.87 |
| A236C | 0.57 | 0.73 | 0.55 | 0.75 | 0.90 | 0.92 | 0.74 |
| C237T | 0.65 | 0.79 | 0.70 | 0.82 | 0.99 | 0.99 | 0.83 |
| C237G | 0.55 | 0.83 | 0.70 | 0.79 | 0.99 | 1.00 | 0.81 |
| A238T | 0.58 | 0.80 | 0.77 | 0.82 | 0.98 | 1.00 | 0.83 |
| A238G | 0.62 | 0.80 | 0.77 | 0.83 | 0.99 | 0.97 | 0.86 |
| A238C | 0.50 | 0.83 | 0.89 | 0.84 | 0.96 | 1.00 | 0.84 |
| G239T | 0.57 | 0.70 | 0.80 | 0.82 | 0.93 | 0.89 | 0.88 |
| G239C | 0.55 | 0.76 | 0.75 | 0.78 | 0.99 | 0.99 | 0.80 |
| G239A | 0.61 | 0.75 | 0.73 | 0.81 | 0.98 | 0.97 | 0.83 |
| T240C | 0.62 | 0.76 | 0.74 | 0.80 | 1.00 | 0.95 | 0.86 |
| T240A | 0.61 | 0.78 | 0.70 | 0.80 | 0.99 | 0.99 | 0.82 |
| T241G | 0.67 | 0.82 | 0.83 | 0.87 | 0.99 | 0.99 | 0.88 |
| T241C | 0.64 | 0.82 | 0.72 | 0.83 | 1.00 | 0.96 | 0.88 |
| T241A | 0.60 | 0.72 | 0.74 | 0.79 | 0.98 | 0.98 | 0.81 |
| TG242T | 0.58 | 0.84 | 0.77 | 0.83 | 1.00 | 1.00 | 0.86 |
| G242T | 0.49 | 0.76 | 0.87 | 0.81 | 0.96 | 0.98 | 0.83 |
| G242C | 0.51 | 0.84 | 0.74 | 0.79 | 1.00 | 1.00 | 0.82 |
| G242A | 0.54 | 0.83 | 0.59 | 0.77 | 0.98 | 1.00 | 0.77 |
| G243T | 0.57 | 0.87 | 0.63 | 0.80 | 1.00 | 1.00 | 0.82 |
| G243A | 0.63 | 0.84 | 0.61 | 0.82 | 1.00 | 1.00 | 0.82 |
| G244T | 0.57 | 0.90 | 0.70 | 0.85 | 0.97 | 1.00 | 0.86 |
| G244A | 0.58 | 0.86 | 0.67 | 0.82 | 1.00 | 1.00 | 0.84 |
| G245T | 0.62 | 0.83 | 0.75 | 0.85 | 0.98 | 1.00 | 0.85 |
| G245C | 0.44 | 0.68 | 0.78 | 0.69 | 0.98 | 0.91 | 0.77 |
| G245A | 0.60 | 0.80 | 0.68 | 0.80 | 1.00 | 1.00 | 0.81 |
| G245GT | 1.00 | 0.02 | 1.00 | 0.99 | 0.01 | 0.02 | 0.99 |
| GA246G | 0.61 | 0.56 | 0.73 | 0.80 | 0.62 | 0.55 | 0.91 |
| A246T | 0.63 | 0.77 | 0.73 | 0.82 | 0.98 | 0.98 | 0.84 |
| A246G | 0.56 | 0.48 | 0.89 | 0.84 | 0.32 | 0.35 | 0.81 |

| | | | | | | | |
|---|---|---|---|---|---|---|---|
| T247G | 0.62 | 0.77 | 0.71 | 0.80 | 1.00 | 0.98 | 0.83 |
| T247C | 0.52 | 0.74 | 0.89 | 0.81 | 0.96 | 0.97 | 0.84 |
| T247A | 0.53 | 0.81 | 0.76 | 0.80 | 1.00 | 1.00 | 0.82 |
| C248T | 0.58 | 0.79 | 0.77 | 0.82 | 0.99 | 0.99 | 0.83 |
| C248G | 0.31 | 0.68 | 1.00 | 0.86 | 0.68 | 0.74 | 0.88 |
| C248A | 0.59 | 0.77 | 0.84 | 0.85 | 0.96 | 0.96 | 0.87 |
| T249G | 0.56 | 0.73 | 0.81 | 0.83 | 0.94 | 0.97 | 0.83 |
| T249C | 0.59 | 0.73 | 0.73 | 0.79 | 0.98 | 0.97 | 0.81 |
| T249A | 0.59 | 0.76 | 0.75 | 0.80 | 0.99 | 0.99 | 0.81 |
| G250T | 0.52 | 0.81 | 0.73 | 0.78 | 0.99 | 0.99 | 0.81 |
| G250C | 0.57 | 0.81 | 0.71 | 0.80 | 1.00 | 1.00 | 0.82 |
| G250A | 0.57 | 0.80 | 0.74 | 0.80 | 1.00 | 0.99 | 0.82 |
| GA251G | 0.60 | 0.69 | 0.64 | 0.76 | 0.95 | 0.96 | 0.76 |
| A251T | 0.57 | 0.78 | 0.69 | 0.78 | 0.99 | 0.99 | 0.80 |
| A251G | 0.59 | 0.76 | 0.72 | 0.80 | 0.97 | 1.00 | 0.80 |
| A251C | 0.54 | 0.78 | 0.66 | 0.76 | 1.00 | 0.99 | 0.79 |
| A252T | 0.56 | 0.72 | 0.74 | 0.82 | 0.92 | 1.00 | 0.78 |
| A252G | 0.56 | 0.62 | 0.88 | 0.94 | 0.77 | 0.78 | 0.96 |
| A252C | 0.42 | 0.83 | 0.82 | 0.76 | 1.00 | 1.00 | 0.81 |
| A253T | 0.60 | 0.80 | 0.72 | 0.81 | 1.00 | 0.99 | 0.83 |
| A253G | 0.61 | 0.73 | 0.72 | 0.81 | 0.96 | 0.98 | 0.81 |
| A253C | 0.71 | 0.78 | 0.78 | 0.88 | 0.95 | 0.91 | 0.93 |
| G254T | 0.55 | 0.68 | 0.57 | 0.75 | 0.85 | 0.97 | 0.66 |
| G254C | 0.61 | 0.77 | 0.64 | 0.79 | 0.99 | 0.99 | 0.80 |
| G254A | 0.60 | 0.84 | 0.64 | 0.81 | 1.00 | 1.00 | 0.82 |
| T255G | 0.49 | 0.61 | 0.85 | 0.91 | 0.73 | 0.81 | 0.84 |
| T255C | 0.57 | 0.85 | 0.73 | 0.82 | 1.00 | 1.00 | 0.84 |
| T255A | 0.67 | 0.85 | 0.68 | 0.86 | 0.99 | 1.00 | 0.85 |
| A256T | 0.61 | 0.83 | 0.75 | 0.83 | 1.00 | 0.99 | 0.86 |
| A256G | 0.57 | 0.87 | 0.78 | 0.84 | 1.00 | 1.00 | 0.87 |
| AG257A | 0.53 | 0.81 | 0.72 | 0.78 | 1.00 | 1.00 | 0.81 |
| G257T | 0.69 | 0.83 | 0.77 | 0.86 | 0.99 | 1.00 | 0.87 |
| G257A | 0.61 | 0.84 | 0.68 | 0.82 | 1.00 | 1.00 | 0.83 |
| G258T | 0.53 | 0.82 | 0.79 | 0.80 | 1.00 | 0.99 | 0.84 |
| G258A | 0.58 | 0.70 | 0.76 | 0.79 | 0.96 | 0.94 | 0.83 |
| G259T | 0.60 | 0.94 | 0.75 | 0.87 | 1.00 | 1.00 | 0.88 |
| G259C | 0.40 | 0.90 | 0.99 | 0.82 | 0.98 | 0.99 | 0.89 |
| G259A | 0.65 | 0.81 | 0.68 | 0.83 | 0.99 | 1.00 | 0.83 |
| G260C | 0.73 | 0.72 | 0.63 | 0.83 | 0.96 | 0.97 | 0.82 |
| G260A | 0.63 | 0.73 | 0.72 | 0.81 | 0.97 | 0.97 | 0.82 |
| GC261G | 0.43 | 0.84 | 0.82 | 0.77 | 1.00 | 0.99 | 0.82 |
| C261T | 0.59 | 0.78 | 0.73 | 0.80 | 0.99 | 0.99 | 0.81 |
| C261G | 0.54 | 0.73 | 0.77 | 0.82 | 0.93 | 0.99 | 0.79 |
| C261A | 0.79 | 0.10 | 0.94 | 0.83 | 0.04 | 0.07 | 0.96 |
| C262T | 0.59 | 0.83 | 0.77 | 0.83 | 1.00 | 0.99 | 0.85 |
| C262G | 0.72 | 0.84 | 1.00 | 1.00 | 0.88 | 0.91 | 1.00 |
| C262A | 0.63 | 0.76 | 0.70 | 0.80 | 1.00 | 0.99 | 0.82 |
| A263T | 0.64 | 0.82 | 0.72 | 0.83 | 1.00 | 0.94 | 0.89 |
| A263G | 0.64 | 0.78 | 0.69 | 0.81 | 0.99 | 1.00 | 0.81 |
| A263C | 0.64 | 0.88 | 0.86 | 0.87 | 1.00 | 0.99 | 0.89 |
| G264T | 0.69 | 0.91 | 0.77 | 0.89 | 1.00 | 1.00 | 0.90 |
| G264C | 0.61 | 0.94 | 0.76 | 0.87 | 1.00 | 1.00 | 0.89 |
| G264A | 0.65 | 0.87 | 0.71 | 0.85 | 1.00 | 1.00 | 0.86 |

| | | | | | | | |
|---|---|---|---|---|---|---|---|
| C265T | 0.53 | 0.81 | 0.78 | 0.80 | 1.00 | 0.99 | 0.83 |
| C265G | 0.45 | 0.92 | 0.77 | 0.80 | 1.00 | 1.00 | 0.84 |
| C265A | 0.64 | 0.82 | 0.74 | 0.84 | 0.98 | 1.00 | 0.84 |
| C266T | 0.70 | 0.94 | 0.75 | 0.90 | 1.00 | 1.00 | 0.91 |
| C266G | 0.23 | 0.63 | 1.00 | 0.97 | 0.34 | 0.67 | 0.68 |
| C266A | 0.36 | 0.81 | 0.81 | 0.71 | 0.99 | 0.99 | 0.76 |
| T267C | 0.51 | 0.84 | 0.85 | 0.81 | 1.00 | 0.95 | 0.89 |
| T267A | 0.71 | 0.93 | 0.67 | 0.89 | 1.00 | 1.00 | 0.89 |
| TA268T | 0.47 | 0.91 | 0.80 | 0.81 | 1.00 | 1.00 | 0.86 |
| A268T | 0.54 | 0.90 | 0.84 | 0.85 | 1.00 | 1.00 | 0.89 |
| A268G | 0.47 | 0.87 | 0.82 | 0.80 | 1.00 | 1.00 | 0.84 |
| A268C | 0.82 | 0.74 | 0.83 | 1.00 | 0.86 | 0.86 | 1.00 |
| C269T | 0.55 | 0.88 | 0.73 | 0.82 | 1.00 | 1.00 | 0.85 |
| C269G | 0.50 | 0.78 | 0.83 | 0.79 | 0.98 | 1.00 | 0.81 |
| C269A | 0.87 | 0.96 | 0.64 | 0.93 | 1.00 | 1.00 | 0.90 |
| T270G | 0.56 | 0.89 | 0.71 | 0.82 | 1.00 | 1.00 | 0.85 |
| T270C | 0.61 | 0.81 | 0.67 | 0.81 | 0.99 | 0.99 | 0.83 |
| T270A | 0.70 | 0.87 | 0.75 | 0.87 | 1.00 | 0.99 | 0.89 |
| TG271T | 0.40 | 0.84 | 0.81 | 0.75 | 0.99 | 1.00 | 0.79 |
| G271T | 0.59 | 0.93 | 0.77 | 0.86 | 1.00 | 1.00 | 0.89 |
| G271C | 0.65 | 0.87 | 0.64 | 0.87 | 0.96 | 1.00 | 0.84 |
| G271A | 0.75 | 0.93 | 0.72 | 0.90 | 1.00 | 0.99 | 0.91 |
| G272T | 0.63 | 0.94 | 0.80 | 0.88 | 1.00 | 1.00 | 0.90 |
| G272C | 0.50 | 0.73 | 1.00 | 0.97 | 0.81 | 0.85 | 0.99 |
| G272A | 0.77 | 0.96 | 0.68 | 0.90 | 1.00 | 1.00 | 0.90 |
| C273T | 0.64 | 0.84 | 0.67 | 0.83 | 1.00 | 1.00 | 0.83 |
| C273G | 0.57 | 0.73 | 0.77 | 0.80 | 0.97 | 0.96 | 0.82 |
| C273A | 0.74 | 0.92 | 0.69 | 0.89 | 1.00 | 1.00 | 0.89 |
| T274G | 0.79 | 0.94 | 0.57 | 0.90 | 1.00 | 1.00 | 0.87 |
| T274C | 0.64 | 0.83 | 0.72 | 0.85 | 0.98 | 1.00 | 0.85 |
| T274A | 0.63 | 0.80 | 0.71 | 0.82 | 1.00 | 0.98 | 0.85 |
| TG275T | 0.41 | 0.64 | 0.86 | 0.69 | 0.93 | 0.78 | 0.88 |
| G275T | 0.62 | 0.84 | 0.72 | 0.84 | 0.99 | 1.00 | 0.84 |
| G275C | 0.71 | 0.90 | 0.65 | 0.88 | 1.00 | 1.00 | 0.87 |
| G275A | 0.61 | 0.81 | 0.83 | 0.84 | 1.00 | 0.98 | 0.88 |
| G276T | 0.56 | 0.85 | 0.80 | 0.83 | 1.00 | 0.99 | 0.86 |
| G276C | 0.54 | 0.76 | 0.65 | 0.75 | 0.98 | 0.99 | 0.76 |
| G276A | 0.64 | 0.94 | 0.84 | 0.89 | 1.00 | 0.99 | 0.91 |
| T277C | 0.46 | 0.72 | 0.88 | 0.77 | 0.96 | 0.98 | 0.79 |
| T277A | 0.53 | 0.72 | 0.76 | 0.77 | 0.96 | 0.97 | 0.79 |
| C278T | 0.53 | 0.85 | 0.84 | 0.82 | 1.00 | 0.99 | 0.86 |
| C278G | 0.51 | 0.58 | 0.97 | 0.96 | 0.72 | 0.75 | 0.96 |
| C278A | 0.53 | 0.76 | 0.84 | 0.83 | 0.94 | 1.00 | 0.82 |
| C279T | 0.63 | 0.67 | 0.69 | 0.80 | 0.93 | 0.88 | 0.86 |
| C279A | 0.59 | 0.77 | 0.72 | 0.79 | 0.99 | 0.99 | 0.80 |
| CT280C | 0.47 | 0.12 | 0.96 | 0.85 | 0.03 | 0.14 | 0.74 |
| T280G | 0.77 | 0.03 | 0.69 | 0.91 | 0.01 | 0.03 | 0.74 |
| T280C | 0.40 | 0.21 | 0.95 | 0.80 | 0.07 | 0.12 | 0.69 |
| T280A | 0.52 | 0.19 | 0.96 | 0.76 | 0.07 | 0.09 | 0.84 |
| C281T | 0.53 | 0.75 | 0.81 | 0.79 | 0.98 | 0.99 | 0.81 |
| C281A | 0.53 | 0.83 | 0.82 | 0.85 | 0.96 | 1.00 | 0.84 |
| A282T | 0.57 | 0.46 | 0.68 | 0.63 | 0.24 | 0.19 | 0.86 |
| A282G | 0.62 | 0.10 | 0.75 | 0.91 | 0.03 | 0.05 | 0.67 |

| | | | | | | | |
|---|---|---|---|---|---|---|---|
| A282C | 0.38 | 0.27 | 0.78 | 0.91 | 0.06 | 0.16 | 0.51 |
| T283C | 0.45 | 0.75 | 0.79 | 0.75 | 0.97 | 0.99 | 0.77 |
| T283A | 0.73 | 0.06 | 0.71 | 0.89 | 0.02 | 0.04 | 0.74 |
| G284T | 0.78 | 0.94 | 0.79 | 0.92 | 1.00 | 0.99 | 0.93 |
| G284A | 0.59 | 0.87 | 0.72 | 0.83 | 1.00 | 1.00 | 0.85 |
| A285T | 0.51 | 0.82 | 0.80 | 0.80 | 1.00 | 1.00 | 0.83 |
| A285G | 0.63 | 0.52 | 0.82 | 0.93 | 0.68 | 0.65 | 0.99 |
| AC286A | 0.43 | 0.68 | 0.54 | 0.61 | 0.39 | 0.41 | 0.62 |
| C286T | 0.63 | 0.55 | 0.73 | 0.88 | 0.68 | 0.67 | 0.91 |
| C286G | 0.59 | 0.46 | 0.70 | 0.75 | 0.22 | 0.23 | 0.75 |
| C286A | 0.60 | 0.80 | 0.67 | 0.80 | 1.00 | 0.99 | 0.81 |
| C287T | 0.59 | 0.97 | 0.86 | 0.89 | 1.00 | 1.00 | 0.92 |
| C287G | 0.64 | 0.76 | 0.77 | 0.89 | 0.92 | 0.99 | 0.84 |
| C287A | 0.56 | 0.87 | 0.74 | 0.82 | 1.00 | 0.99 | 0.85 |
| C288T | 0.63 | 0.91 | 0.75 | 0.86 | 1.00 | 1.00 | 0.88 |
| C288A | 0.59 | 0.65 | 0.68 | 0.81 | 0.86 | 0.84 | 0.84 |
| T289C | 0.66 | 0.15 | 0.91 | 0.76 | 0.06 | 0.06 | 0.93 |
| T289A | 0.60 | 0.22 | 0.79 | 0.76 | 0.08 | 0.09 | 0.80 |
| C290T | 0.72 | 0.80 | 0.63 | 0.83 | 0.99 | 0.99 | 0.83 |
| C290A | 0.62 | 0.37 | 0.83 | 0.76 | 0.19 | 0.18 | 0.86 |
| T291C | 0.62 | 0.18 | 0.82 | 0.94 | 0.06 | 0.10 | 0.69 |
| T291A | 0.80 | 0.06 | 0.77 | 0.98 | 0.02 | 0.03 | 0.73 |
| C292T | 0.76 | 0.93 | 0.65 | 0.89 | 1.00 | 1.00 | 0.88 |
| C292G | 0.71 | 0.11 | 0.75 | 0.81 | 0.04 | 0.04 | 0.81 |
| C292A | 0.68 | 0.10 | 0.81 | 0.82 | 0.03 | 0.04 | 0.82 |
| T293G | 0.98 | 0.03 | 1.00 | 0.95 | 0.01 | 0.04 | 0.98 |
| T293C | 0.61 | 0.87 | 0.73 | 0.84 | 1.00 | 1.00 | 0.86 |
| G294T | 0.59 | 0.49 | 0.80 | 0.81 | 0.30 | 0.32 | 0.79 |
| G294C | 0.67 | 0.64 | 0.65 | 0.87 | 0.84 | 0.89 | 0.82 |
| G294A | 0.65 | 0.86 | 0.80 | 0.87 | 1.00 | 1.00 | 0.88 |
| C295T | 0.63 | 0.41 | 0.92 | 0.94 | 0.20 | 0.27 | 0.75 |
| C295G | 0.92 | 0.01 | 0.96 | 0.90 | 0.01 | 0.00 | 0.97 |
| C295A | 0.90 | 0.01 | 0.93 | 0.92 | 0.01 | 0.01 | 0.92 |
| CA296C | 0.95 | 0.01 | 0.90 | 0.91 | 0.01 | 0.00 | 0.95 |
| A296T | 0.96 | 0.01 | 0.94 | 0.92 | 0.00 | 0.00 | 0.96 |
| A296G | 0.95 | 0.01 | 0.94 | 1.00 | 0.00 | 0.01 | 0.88 |
| A296C | 0.96 | 0.01 | 0.89 | 0.92 | 0.00 | 0.00 | 0.93 |
| G297T | 0.95 | 0.01 | 0.99 | 0.95 | 0.01 | 0.00 | 0.94 |
| G297C | 0.95 | 0.01 | 0.96 | 0.99 | 0.00 | 0.01 | 0.90 |
| G297A | 0.92 | 0.01 | 0.96 | 0.93 | 0.01 | 0.00 | 0.94 |
| G298T | 0.92 | 0.03 | 0.79 | 0.89 | 0.01 | 0.01 | 0.88 |
| G298C | 0.88 | 0.05 | 0.75 | 0.80 | 0.02 | 0.02 | 0.93 |
| G298A | 0.73 | 0.12 | 0.79 | 0.79 | 0.05 | 0.05 | 0.88 |
| A299T | 0.66 | 0.87 | 0.68 | 0.84 | 1.00 | 1.00 | 0.85 |
| A299G | 0.90 | 0.07 | 0.84 | 0.86 | 0.03 | 0.03 | 0.93 |
| T300C | 0.82 | 0.89 | 0.80 | 0.91 | 1.00 | 0.99 | 0.92 |
| T300A | 0.59 | 0.90 | 0.63 | 0.83 | 1.00 | 1.00 | 0.84 |
| A301T | 0.69 | 0.81 | 0.70 | 0.84 | 0.99 | 1.00 | 0.85 |
| A301G | 0.57 | 0.77 | 0.70 | 0.79 | 0.98 | 0.99 | 0.80 |
| T302C | 0.60 | 0.87 | 0.75 | 0.84 | 1.00 | 1.00 | 0.86 |
| T302A | 0.59 | 0.75 | 0.81 | 0.82 | 0.97 | 0.96 | 0.85 |
| T303C | 0.71 | 0.95 | 0.78 | 0.91 | 0.99 | 1.00 | 0.91 |
| T303A | 0.65 | 0.90 | 0.70 | 0.86 | 0.99 | 1.00 | 0.86 |

| | | | | | | | |
|---|---|---|---|---|---|---|---|
| TG304T | 0.62 | 0.91 | 0.75 | 0.86 | 1.00 | 1.00 | 0.87 |
| G304T | 0.69 | 0.97 | 0.88 | 0.92 | 1.00 | 1.00 | 0.94 |
| G304A | 0.66 | 0.94 | 0.68 | 0.88 | 1.00 | 1.00 | 0.88 |
| G305T | 0.73 | 0.97 | 0.71 | 0.91 | 0.99 | 1.00 | 0.90 |
| G305C | 0.55 | 0.97 | 0.80 | 0.86 | 1.00 | 1.00 | 0.88 |
| G305A | 0.72 | 0.98 | 0.74 | 0.91 | 1.00 | 1.00 | 0.92 |
| G306T | 0.76 | 0.97 | 0.80 | 0.92 | 1.00 | 1.00 | 0.93 |
| G306C | 0.75 | 0.98 | 0.79 | 0.92 | 1.00 | 1.00 | 0.93 |
| G306A | 0.71 | 0.98 | 0.78 | 0.91 | 1.00 | 1.00 | 0.92 |
| C307T | 0.56 | 0.26 | 0.95 | 0.91 | 0.09 | 0.14 | 0.73 |
| C307G | 0.96 | 0.04 | 0.74 | 0.88 | 0.01 | 0.01 | 0.89 |
| C307A | 0.57 | 0.64 | 0.76 | 0.89 | 0.81 | 0.90 | 0.82 |
| T308G | 0.58 | 0.56 | 0.80 | 0.90 | 0.73 | 0.72 | 0.94 |
| T308C | 0.70 | 0.95 | 0.76 | 0.91 | 0.99 | 1.00 | 0.90 |
| T308A | 0.58 | 0.82 | 0.69 | 0.80 | 0.99 | 1.00 | 0.82 |
| TG309T | 0.58 | 0.93 | 0.74 | 0.88 | 0.97 | 1.00 | 0.87 |
| G309T | 0.71 | 0.97 | 0.51 | 0.87 | 1.00 | 1.00 | 0.84 |
| G309C | 0.79 | 0.99 | 0.78 | 0.94 | 1.00 | 1.00 | 0.94 |
| G309A | 0.59 | 0.89 | 0.64 | 0.82 | 1.00 | 1.00 | 0.83 |
| G310T | 0.74 | 0.97 | 0.73 | 0.91 | 1.00 | 0.96 | 0.95 |
| G310A | 0.70 | 0.97 | 0.83 | 0.91 | 1.00 | 0.98 | 0.94 |
| G311T | 0.83 | 0.92 | 0.96 | 0.94 | 0.99 | 0.99 | 0.96 |
| G311A | 0.83 | 0.98 | 0.74 | 0.93 | 1.00 | 1.00 | 0.92 |
| C312T | 0.53 | 0.45 | 0.94 | 0.81 | 0.28 | 0.29 | 0.82 |
| C312A | 0.59 | 0.79 | 0.72 | 0.80 | 1.00 | 0.99 | 0.82 |
| G313T | 0.60 | 0.37 | 0.85 | 0.89 | 0.15 | 0.20 | 0.73 |
| G313C | 0.49 | 0.83 | 0.85 | 0.80 | 1.00 | 0.99 | 0.84 |
| G313A | 0.55 | 0.46 | 0.85 | 0.84 | 0.24 | 0.29 | 0.75 |
| C314A | 0.54 | 0.63 | 0.80 | 0.83 | 0.84 | 0.77 | 0.93 |
| T315G | 0.57 | 0.58 | 0.78 | 0.88 | 0.76 | 0.74 | 0.92 |
| T315C | 0.65 | 0.89 | 0.75 | 0.86 | 1.00 | 1.00 | 0.88 |
| T315A | 0.55 | 0.65 | 0.78 | 0.85 | 0.85 | 0.87 | 0.84 |
| G316T | 0.62 | 0.56 | 0.80 | 0.97 | 0.69 | 0.75 | 0.92 |
| G316C | 0.84 | 0.52 | 0.69 | 0.98 | 0.66 | 0.66 | 0.97 |
| G316A | 0.67 | 0.64 | 0.66 | 0.84 | 0.87 | 0.85 | 0.87 |
| T317C | 0.62 | 0.93 | 0.79 | 0.87 | 1.00 | 0.92 | 0.96 |
| T317A | 0.51 | 0.72 | 0.73 | 0.72 | 1.00 | 0.96 | 0.77 |
| G318T | 0.63 | 0.90 | 0.66 | 0.85 | 1.00 | 1.00 | 0.86 |
| G318C | 0.69 | 0.93 | 0.69 | 0.89 | 0.99 | 1.00 | 0.89 |
| G318A | 0.65 | 0.68 | 0.67 | 0.78 | 0.97 | 0.92 | 0.83 |
| G319T | 0.60 | 0.69 | 0.73 | 0.81 | 0.94 | 0.93 | 0.83 |
| G319C | 0.42 | 0.76 | 0.91 | 0.77 | 0.98 | 0.98 | 0.81 |
| G319A | 0.69 | 0.85 | 0.74 | 0.86 | 1.00 | 0.99 | 0.87 |
| C320T | 0.59 | 0.36 | 0.91 | 0.78 | 0.19 | 0.19 | 0.86 |
| C320G | 0.88 | 0.03 | 0.95 | 0.92 | 0.01 | 0.02 | 0.92 |
| C320A | 0.50 | 0.69 | 0.72 | 0.73 | 0.96 | 0.98 | 0.73 |
| T321G | 0.53 | 0.75 | 0.76 | 0.77 | 0.98 | 0.99 | 0.79 |
| T321C | 0.63 | 0.89 | 0.75 | 0.86 | 1.00 | 1.00 | 0.87 |
| T321A | 0.53 | 0.62 | 0.82 | 0.85 | 0.81 | 0.77 | 0.92 |
| G322T | 0.51 | 0.36 | 0.99 | 0.85 | 0.17 | 0.23 | 0.78 |
| G322C | 0.54 | 0.63 | 0.79 | 0.97 | 0.72 | 0.89 | 0.80 |
| G322A | 0.53 | 0.37 | 0.99 | 0.88 | 0.17 | 0.22 | 0.76 |
| A323T | 0.58 | 0.70 | 0.80 | 0.85 | 0.90 | 0.90 | 0.88 |

| | | | | | | | |
|---|---|---|---|---|---|---|---|
| A323G | 0.61 | 0.48 | 0.84 | 0.79 | 0.34 | 0.32 | 0.87 |
| A323C | 0.66 | 0.82 | 0.73 | 0.84 | 1.00 | 0.98 | 0.87 |
| C324T | 0.60 | 0.55 | 0.82 | 0.93 | 0.71 | 0.71 | 0.95 |
| C324A | 0.62 | 0.60 | 0.94 | 1.00 | 0.75 | 0.76 | 1.00 |
| T325G | 0.30 | 0.98 | 1.00 | 0.78 | 0.97 | 1.00 | 0.91 |
| T325C | 0.70 | 0.94 | 0.86 | 0.91 | 1.00 | 0.99 | 0.94 |
| T325A | 0.58 | 0.63 | 0.73 | 0.81 | 0.87 | 0.82 | 0.87 |
| TG326T | 0.54 | 0.72 | 0.89 | 0.89 | 0.88 | 0.89 | 0.92 |
| G326T | 0.52 | 0.74 | 0.80 | 0.79 | 0.97 | 0.98 | 0.81 |
| G326C | 0.61 | 0.86 | 0.69 | 0.84 | 1.00 | 1.00 | 0.85 |
| G326A | 0.60 | 0.70 | 0.75 | 0.82 | 0.93 | 0.93 | 0.84 |
| T327C | 0.72 | 0.96 | 0.74 | 0.91 | 1.00 | 1.00 | 0.91 |
| T327A | 0.59 | 0.85 | 0.73 | 0.83 | 0.99 | 1.00 | 0.84 |
| G328T | 0.66 | 0.85 | 0.67 | 0.86 | 0.99 | 1.00 | 0.85 |
| G328C | 0.77 | 0.92 | 0.71 | 0.90 | 1.00 | 1.00 | 0.90 |
| G328A | 0.65 | 0.78 | 0.74 | 0.83 | 0.99 | 0.99 | 0.84 |
| T329G | 0.53 | 0.52 | 0.92 | 0.81 | 0.56 | 0.49 | 0.96 |
| T329C | 0.69 | 0.96 | 0.77 | 0.90 | 1.00 | 1.00 | 0.91 |
| T329A | 0.66 | 0.92 | 0.74 | 0.88 | 1.00 | 1.00 | 0.89 |
| TG330T | 0.65 | 0.96 | 0.86 | 0.90 | 1.00 | 1.00 | 0.92 |
| G330T | 0.77 | 0.97 | 0.65 | 0.92 | 0.99 | 1.00 | 0.90 |
| G330C | 0.86 | 0.97 | 0.63 | 0.94 | 0.97 | 1.00 | 0.90 |
| G330A | 0.52 | 0.86 | 0.57 | 0.75 | 1.00 | 0.99 | 0.77 |
| G331T | 0.66 | 0.96 | 0.78 | 0.89 | 1.00 | 1.00 | 0.91 |
| G331C | 0.76 | 0.97 | 0.75 | 0.92 | 1.00 | 1.00 | 0.91 |
| G331A | 0.71 | 0.97 | 0.60 | 0.88 | 1.00 | 1.00 | 0.88 |
| G332C | 0.58 | 0.93 | 0.86 | 0.87 | 1.00 | 1.00 | 0.90 |
| G332A | 0.78 | 0.98 | 0.80 | 0.93 | 0.99 | 1.00 | 0.93 |
| T333C | 0.67 | 0.94 | 0.73 | 0.88 | 1.00 | 0.99 | 0.90 |
| T333A | 0.68 | 0.93 | 0.72 | 0.88 | 1.00 | 1.00 | 0.89 |
| A334T | 0.56 | 0.79 | 0.66 | 0.77 | 0.99 | 0.99 | 0.79 |
| A334G | 0.58 | 0.82 | 0.79 | 0.83 | 1.00 | 0.99 | 0.86 |
| T335C | 0.60 | 0.83 | 0.77 | 0.83 | 1.00 | 1.00 | 0.85 |
| T335A | 0.56 | 0.69 | 0.72 | 0.74 | 0.99 | 0.96 | 0.78 |
| C336T | 0.68 | 0.19 | 0.91 | 0.83 | 0.07 | 0.09 | 0.86 |
| C336A | 0.89 | 0.83 | 0.64 | 0.90 | 0.97 | 0.97 | 0.89 |
| A337T | 0.58 | 0.74 | 0.80 | 0.82 | 0.97 | 0.97 | 0.84 |
| A337G | 0.64 | 0.90 | 0.72 | 0.86 | 1.00 | 1.00 | 0.88 |
| A337C | 0.59 | 0.75 | 0.67 | 0.79 | 0.97 | 1.00 | 0.78 |
| A338T | 0.57 | 0.79 | 0.72 | 0.80 | 0.99 | 0.99 | 0.81 |
| A338G | 0.61 | 0.67 | 0.73 | 0.85 | 0.88 | 0.89 | 0.85 |
| A338C | 0.59 | 0.79 | 0.78 | 0.82 | 1.00 | 0.99 | 0.84 |
| C339T | 0.61 | 0.20 | 0.94 | 0.84 | 0.08 | 0.09 | 0.82 |
| C339A | 0.70 | 0.17 | 0.83 | 0.79 | 0.07 | 0.07 | 0.88 |
| G340T | 0.44 | 0.64 | 1.00 | 0.93 | 0.76 | 0.80 | 0.94 |
| G340C | 0.60 | 0.62 | 0.64 | 0.76 | 0.73 | 0.68 | 0.82 |
| G340A | 0.55 | 0.58 | 0.81 | 0.90 | 0.72 | 0.76 | 0.88 |
| T341G | 0.74 | 0.12 | 0.87 | 0.83 | 0.05 | 0.05 | 0.88 |
| T341C | 0.63 | 0.89 | 0.74 | 0.86 | 1.00 | 1.00 | 0.88 |
| T341A | 0.65 | 0.77 | 0.75 | 0.83 | 0.98 | 0.98 | 0.85 |
| G342T | 0.63 | 0.71 | 0.72 | 0.79 | 0.98 | 0.98 | 0.81 |
| G342A | 0.49 | 0.48 | 0.95 | 0.78 | 0.33 | 0.33 | 0.85 |
| A343T | 0.57 | 0.60 | 0.81 | 0.92 | 0.76 | 0.78 | 0.92 |

| | | | | | | | |
|---|---|---|---|---|---|---|---|
| A343G | 0.56 | 0.71 | 0.85 | 0.91 | 0.86 | 0.90 | 0.89 |
| A343C | 0.48 | 0.60 | 0.98 | 0.96 | 0.72 | 0.77 | 0.95 |
| AC344A | 0.47 | 0.91 | 0.88 | 0.84 | 0.99 | 1.00 | 0.88 |
| C344T | 0.60 | 0.74 | 0.70 | 0.79 | 0.98 | 0.99 | 0.80 |
| C344A | 0.64 | 0.18 | 0.90 | 0.91 | 0.06 | 0.09 | 0.76 |
| C345T | 0.57 | 0.53 | 0.74 | 0.74 | 0.55 | 0.46 | 0.90 |
| C345G | 0.69 | 0.53 | 0.83 | 0.99 | 0.69 | 0.70 | 0.99 |
| C345A | 0.58 | 0.48 | 0.83 | 0.85 | 0.27 | 0.32 | 0.76 |
| G346T | 0.60 | 0.83 | 0.80 | 0.84 | 1.00 | 0.98 | 0.87 |
| G346C | 0.63 | 0.73 | 0.80 | 0.88 | 0.91 | 0.88 | 0.92 |
| G346A | 0.65 | 0.88 | 0.69 | 0.86 | 1.00 | 1.00 | 0.86 |
| T347G | 0.55 | 0.48 | 0.91 | 0.87 | 0.33 | 0.38 | 0.81 |
| T347C | 0.65 | 0.91 | 0.75 | 0.87 | 1.00 | 0.99 | 0.88 |
| T347A | 0.69 | 0.90 | 0.68 | 0.87 | 1.00 | 1.00 | 0.87 |
| TG348T | 0.66 | 0.92 | 0.53 | 0.84 | 1.00 | 1.00 | 0.82 |
| G348T | 0.58 | 0.91 | 0.62 | 0.82 | 1.00 | 0.99 | 0.84 |
| G348C | 0.55 | 0.99 | 0.99 | 0.89 | 1.00 | 0.99 | 0.95 |
| G349T | 0.58 | 0.86 | 0.63 | 0.80 | 1.00 | 1.00 | 0.81 |
| G349C | 0.51 | 0.53 | 0.76 | 0.75 | 0.31 | 0.33 | 0.76 |
| G349A | 0.72 | 0.98 | 0.74 | 0.91 | 0.99 | 1.00 | 0.91 |
| G350C | 0.49 | 0.67 | 0.54 | 0.55 | 0.51 | 0.39 | 0.75 |
| G350A | 0.78 | 0.99 | 0.80 | 0.93 | 1.00 | 0.96 | 0.96 |
| G35G1 | 0.99 | 0.91 | 0.11 | 0.68 | 0.96 | 0.95 | 0.50 |
| GT351G | 0.53 | 0.93 | 0.68 | 0.82 | 1.00 | 1.00 | 0.85 |
| T351C | 0.59 | 0.89 | 0.67 | 0.83 | 1.00 | 1.00 | 0.85 |
| T351A | 0.72 | 0.96 | 0.74 | 0.90 | 1.00 | 0.98 | 0.92 |
| TG352T | 0.59 | 0.82 | 0.62 | 0.79 | 0.99 | 1.00 | 0.79 |
| G352T | 0.62 | 0.89 | 0.75 | 0.86 | 1.00 | 0.99 | 0.88 |
| G352C | 0.65 | 0.95 | 0.83 | 0.89 | 1.00 | 0.99 | 0.91 |
| G352A | 0.58 | 0.77 | 0.70 | 0.79 | 0.98 | 0.99 | 0.81 |
| G353T | 0.86 | 0.95 | 0.68 | 0.93 | 1.00 | 1.00 | 0.91 |
| G353C | 0.59 | 0.94 | 0.68 | 0.85 | 1.00 | 0.97 | 0.89 |
| G353A | 0.62 | 0.97 | 0.83 | 0.89 | 1.00 | 1.00 | 0.91 |
| GT354G | 0.95 | 0.09 | 0.91 | 0.90 | 0.05 | 0.04 | 0.95 |
| T354G | 0.66 | 0.76 | 0.69 | 0.79 | 1.00 | 0.94 | 0.85 |
| T354C | 0.56 | 0.71 | 0.67 | 0.75 | 0.96 | 0.97 | 0.76 |
| T354A | 0.66 | 0.92 | 0.75 | 0.88 | 1.00 | 1.00 | 0.89 |
| G355T | 0.74 | 0.26 | 0.64 | 0.78 | 0.10 | 0.09 | 0.77 |
| G355C | 0.33 | 0.62 | 0.99 | 0.90 | 0.61 | 0.78 | 0.80 |
| G355A | 0.62 | 0.53 | 0.70 | 0.85 | 0.38 | 0.43 | 0.75 |
| GA356G | 0.53 | 0.69 | 0.67 | 0.70 | 0.99 | 0.90 | 0.79 |
| A356T | 0.56 | 0.69 | 0.80 | 0.87 | 0.86 | 0.92 | 0.84 |
| A356G | 0.79 | 0.06 | 0.97 | 0.87 | 0.02 | 0.03 | 0.93 |
| G357T | 0.58 | 0.44 | 0.91 | 0.72 | 0.27 | 0.23 | 0.94 |
| G357A | 0.57 | 0.74 | 0.70 | 0.77 | 0.98 | 1.00 | 0.77 |
| A358T | 0.78 | 0.45 | 0.58 | 0.74 | 0.26 | 0.22 | 0.82 |
| A358G | 0.89 | 0.04 | 0.88 | 0.91 | 0.01 | 0.02 | 0.90 |
| A358C | 0.54 | 0.70 | 0.77 | 0.79 | 0.94 | 0.93 | 0.82 |
| G359T | 0.65 | 0.83 | 0.72 | 0.84 | 1.00 | 0.96 | 0.88 |
| G359C | 0.64 | 0.85 | 0.56 | 0.81 | 1.00 | 1.00 | 0.80 |
| G359A | 0.53 | 0.75 | 0.74 | 0.79 | 0.96 | 0.99 | 0.79 |
| C360T | 0.53 | 0.52 | 0.87 | 0.74 | 0.55 | 0.42 | 1.00 |
| C360A | 0.60 | 0.74 | 0.79 | 0.85 | 0.94 | 0.96 | 0.85 |

| | | | | | | | |
|---|---|---|---|---|---|---|---|
| T361G | 0.62 | 0.87 | 0.60 | 0.84 | 0.99 | 1.00 | 0.84 |
| T361C | 0.56 | 0.73 | 0.77 | 0.79 | 0.98 | 0.93 | 0.84 |
| T361A | 0.60 | 0.64 | 0.79 | 0.90 | 0.82 | 0.81 | 0.93 |
| G362C | 0.63 | 0.28 | 0.98 | 0.80 | 0.14 | 0.15 | 0.91 |
| G362A | 0.60 | 0.69 | 0.77 | 0.87 | 0.88 | 0.90 | 0.87 |
| GC363G | 0.51 | 0.65 | 0.83 | 0.84 | 0.85 | 0.86 | 0.86 |
| C363T | 0.61 | 0.59 | 0.78 | 0.90 | 0.78 | 0.75 | 0.95 |
| C363G | 0.56 | 0.57 | 0.63 | 0.75 | 0.38 | 0.41 | 0.71 |
| C363A | 0.53 | 0.74 | 0.80 | 0.81 | 0.95 | 0.99 | 0.81 |
| C364T | 0.66 | 0.46 | 0.77 | 0.77 | 0.28 | 0.26 | 0.85 |
| C364G | 0.33 | 0.64 | 1.00 | 0.82 | 0.70 | 0.72 | 0.91 |
| C364A | 0.65 | 0.83 | 0.62 | 0.82 | 0.99 | 1.00 | 0.82 |
| A365T | 0.56 | 0.59 | 0.80 | 0.91 | 0.75 | 0.79 | 0.90 |
| A365G | 0.60 | 0.83 | 0.74 | 0.83 | 1.00 | 1.00 | 0.85 |
| A365C | 0.51 | 0.53 | 0.63 | 0.76 | 0.23 | 0.30 | 0.63 |
| G366T | 0.60 | 0.73 | 0.75 | 0.81 | 0.97 | 0.98 | 0.82 |
| G366C | 0.47 | 0.48 | 0.85 | 0.69 | 0.29 | 0.27 | 0.85 |
| G366A | 0.63 | 0.66 | 0.76 | 0.91 | 0.83 | 0.83 | 0.92 |
| C367T | 0.63 | 0.64 | 0.82 | 0.96 | 0.79 | 0.79 | 0.97 |
| C367G | 0.64 | 0.71 | 0.79 | 0.92 | 0.86 | 0.89 | 0.91 |
| C367A | 0.62 | 0.87 | 0.79 | 0.86 | 0.99 | 1.00 | 0.87 |
| C367CA | 0.69 | 0.90 | 0.65 | 0.86 | 1.00 | 0.99 | 0.87 |
| A368T | 0.58 | 0.60 | 0.76 | 0.87 | 0.78 | 0.77 | 0.89 |
| A368G | 0.60 | 0.83 | 0.72 | 0.84 | 0.97 | 1.00 | 0.83 |
| A368C | 0.55 | 0.59 | 0.86 | 0.92 | 0.76 | 0.75 | 0.96 |
| C369T | 0.62 | 0.59 | 0.71 | 0.89 | 0.71 | 0.75 | 0.86 |
| C369G | 0.56 | 0.79 | 0.77 | 0.80 | 1.00 | 0.99 | 0.83 |
| C369A | 0.59 | 0.89 | 0.71 | 0.83 | 1.00 | 1.00 | 0.85 |
| G370T | 0.64 | 0.31 | 0.85 | 0.83 | 0.13 | 0.15 | 0.80 |
| G370A | 0.60 | 0.71 | 0.73 | 0.80 | 0.96 | 0.95 | 0.83 |
| A371T | 0.60 | 0.69 | 0.70 | 0.81 | 0.92 | 0.95 | 0.79 |
| A371G | 0.49 | 0.24 | 0.93 | 0.94 | 0.08 | 0.14 | 0.65 |
| G372A | 0.73 | 0.68 | 0.65 | 0.84 | 0.93 | 0.78 | 0.99 |
| GT373G | 0.32 | 0.64 | 0.96 | 0.73 | 0.80 | 0.83 | 0.76 |
| T373C | 0.52 | 0.72 | 0.78 | 0.78 | 0.96 | 0.97 | 0.80 |
| T373A | 0.66 | 0.88 | 0.71 | 0.86 | 1.00 | 1.00 | 0.87 |
| T374G | 0.50 | 0.78 | 0.62 | 0.71 | 1.00 | 0.99 | 0.73 |
| T374C | 0.54 | 0.53 | 0.81 | 0.83 | 0.52 | 0.54 | 0.84 |
| T374A | 0.61 | 0.78 | 0.75 | 0.81 | 1.00 | 0.99 | 0.83 |
| C375T | 0.60 | 0.63 | 0.70 | 0.82 | 0.84 | 0.81 | 0.87 |
| C375G | 0.68 | 0.66 | 0.67 | 0.86 | 0.87 | 0.85 | 0.88 |
| C376T | 0.55 | 0.59 | 0.81 | 0.88 | 0.77 | 0.76 | 0.92 |
| C376G | 0.71 | 0.46 | 0.87 | 0.89 | 0.31 | 0.33 | 0.86 |
| C376A | 0.63 | 0.69 | 0.68 | 0.80 | 0.94 | 0.94 | 0.80 |
| C376CG | 0.60 | 0.51 | 0.80 | 0.95 | 0.37 | 0.49 | 0.74 |
| CG377C | 0.52 | 0.88 | 0.75 | 0.81 | 1.00 | 0.99 | 0.85 |
| G377T | 0.61 | 0.89 | 0.79 | 0.86 | 1.00 | 1.00 | 0.88 |
| G377C | 0.44 | 0.82 | 0.86 | 0.78 | 1.00 | 0.99 | 0.84 |
| G377A | 0.62 | 0.89 | 0.63 | 0.83 | 1.00 | 1.00 | 0.84 |
| G378A | 0.63 | 0.93 | 0.71 | 0.87 | 1.00 | 1.00 | 0.88 |
| G379T | 0.62 | 0.94 | 0.72 | 0.87 | 1.00 | 1.00 | 0.88 |
| G379A | 0.65 | 0.97 | 0.83 | 0.90 | 1.00 | 1.00 | 0.92 |
| G380T | 0.51 | 0.92 | 0.70 | 0.81 | 1.00 | 1.00 | 0.84 |

| | | | | | | | |
|---|---|---|---|---|---|---|---|
| G380C | 0.87 | 0.92 | 0.75 | 0.93 | 1.00 | 0.99 | 0.92 |
| G380A | 0.71 | 0.95 | 0.74 | 0.90 | 1.00 | 1.00 | 0.90 |
| G38G4 | 0.63 | 0.83 | 0.71 | 0.83 | 1.00 | 1.00 | 0.84 |
| G381C | 0.56 | 0.94 | 0.70 | 0.84 | 1.00 | 1.00 | 0.87 |
| G381A | 0.63 | 0.95 | 0.68 | 0.88 | 0.99 | 1.00 | 0.88 |
| G382A | 0.61 | 0.84 | 0.62 | 0.80 | 1.00 | 1.00 | 0.81 |
| A383G | 0.59 | 0.79 | 0.72 | 0.80 | 1.00 | 0.95 | 0.86 |
| A383C | 0.57 | 0.82 | 0.62 | 0.78 | 0.99 | 1.00 | 0.79 |
| C384T | 0.56 | 0.41 | 0.93 | 0.76 | 0.23 | 0.21 | 0.88 |
| C384G | 1.00 | 0.45 | 0.51 | 0.80 | 0.33 | 0.28 | 0.87 |
| C384A | 0.59 | 0.45 | 0.89 | 0.85 | 0.26 | 0.29 | 0.81 |
| A385T | 0.60 | 0.73 | 0.65 | 0.76 | 0.99 | 0.98 | 0.77 |
| A385G | 0.59 | 0.78 | 0.78 | 0.82 | 0.98 | 0.98 | 0.85 |
| A385C | 0.46 | 0.76 | 0.82 | 0.77 | 0.97 | 0.99 | 0.80 |
| T386G | 0.37 | 0.83 | 0.95 | 0.76 | 0.99 | 1.00 | 0.82 |
| T386C | 0.64 | 0.85 | 0.70 | 0.84 | 1.00 | 1.00 | 0.85 |
| T386A | 0.65 | 0.80 | 0.73 | 0.83 | 0.99 | 0.99 | 0.85 |
| TG387T | 0.61 | 0.81 | 0.72 | 0.82 | 1.00 | 0.99 | 0.84 |
| G387T | 0.54 | 0.71 | 0.78 | 0.78 | 0.96 | 0.93 | 0.83 |
| G387A | 0.58 | 0.51 | 0.81 | 0.77 | 0.35 | 0.34 | 0.84 |
| G388T | 0.51 | 0.78 | 0.75 | 0.77 | 0.99 | 1.00 | 0.80 |
| G388C | 0.59 | 0.76 | 0.74 | 0.79 | 0.99 | 0.99 | 0.81 |
| GT389G | 0.61 | 0.80 | 0.68 | 0.80 | 1.00 | 1.00 | 0.81 |
| T389G | 0.56 | 0.69 | 0.85 | 0.92 | 0.83 | 0.89 | 0.89 |
| T389C | 0.56 | 0.92 | 0.73 | 0.84 | 1.00 | 1.00 | 0.87 |
| T389A | 0.67 | 0.96 | 0.76 | 0.89 | 1.00 | 1.00 | 0.91 |
| T390G | 0.59 | 0.77 | 0.73 | 0.80 | 0.99 | 1.00 | 0.81 |
| T390C | 0.60 | 0.94 | 0.76 | 0.86 | 1.00 | 1.00 | 0.89 |
| T390A | 0.60 | 0.49 | 0.81 | 0.77 | 0.33 | 0.31 | 0.86 |
| G391T | 0.61 | 0.52 | 0.85 | 0.96 | 0.66 | 0.68 | 0.95 |
| G391C | 0.55 | 0.63 | 0.79 | 0.91 | 0.77 | 0.85 | 0.84 |
| G391A | 0.60 | 0.63 | 0.82 | 0.92 | 0.80 | 0.79 | 0.96 |
| T392C | 0.63 | 0.78 | 0.75 | 0.84 | 0.97 | 0.98 | 0.84 |
| C393T | 0.60 | 0.49 | 0.84 | 0.90 | 0.31 | 0.38 | 0.77 |
| C393G | 0.50 | 0.71 | 0.79 | 0.83 | 0.88 | 0.99 | 0.77 |
| C393A | 0.70 | 0.65 | 0.80 | 0.99 | 0.79 | 0.80 | 0.98 |
| T394G | 0.66 | 0.82 | 0.73 | 0.84 | 0.99 | 0.99 | 0.86 |
| T394C | 0.64 | 0.93 | 0.68 | 0.86 | 1.00 | 0.97 | 0.90 |
| T394A | 0.57 | 0.72 | 0.80 | 0.82 | 0.96 | 0.92 | 0.87 |
| G395T | 0.54 | 0.81 | 0.67 | 0.78 | 0.99 | 0.99 | 0.79 |
| G395C | 0.59 | 0.79 | 0.77 | 0.81 | 1.00 | 0.99 | 0.83 |
| G395A | 0.62 | 0.79 | 0.70 | 0.81 | 0.99 | 0.99 | 0.83 |
| G395GC | 0.61 | 0.84 | 0.72 | 0.83 | 1.00 | 1.00 | 0.85 |
| GC396G | 0.57 | 0.86 | 0.70 | 0.82 | 1.00 | 1.00 | 0.84 |
| C396T | 0.58 | 0.85 | 0.69 | 0.82 | 1.00 | 1.00 | 0.83 |
| C396G | 0.56 | 0.73 | 0.86 | 0.85 | 0.93 | 0.92 | 0.89 |
| C397T | 0.55 | 0.87 | 0.71 | 0.82 | 0.99 | 1.00 | 0.84 |
| C397G | 0.60 | 0.93 | 0.83 | 0.87 | 1.00 | 1.00 | 0.90 |
| C397A | 0.65 | 0.93 | 0.77 | 0.88 | 1.00 | 0.99 | 0.91 |
| C398T | 0.64 | 0.92 | 0.69 | 0.87 | 1.00 | 1.00 | 0.88 |
| C398A | 0.60 | 0.90 | 0.70 | 0.84 | 1.00 | 1.00 | 0.86 |
| C399T | 0.66 | 0.78 | 0.75 | 0.83 | 0.99 | 0.99 | 0.85 |
| C399A | 0.60 | 0.87 | 0.72 | 0.83 | 1.00 | 1.00 | 0.85 |

| | | | | | | | |
|---|---|---|---|---|---|---|---|
| C400T | 0.60 | 0.80 | 0.71 | 0.81 | 1.00 | 0.99 | 0.82 |
| C400G | 0.62 | 0.84 | 0.68 | 0.82 | 1.00 | 0.99 | 0.84 |
| C400A | 0.60 | 0.82 | 0.71 | 0.82 | 1.00 | 0.99 | 0.84 |
| T401G | 0.75 | 0.84 | 0.56 | 0.85 | 0.99 | 1.00 | 0.83 |
| T401C | 0.58 | 0.88 | 0.71 | 0.86 | 0.97 | 1.00 | 0.85 |
| T401A | 0.58 | 0.85 | 0.70 | 0.82 | 1.00 | 1.00 | 0.84 |
| G402T | 0.49 | 0.70 | 0.77 | 0.76 | 0.94 | 0.99 | 0.75 |
| G402C | 0.57 | 0.80 | 0.74 | 0.80 | 1.00 | 0.97 | 0.83 |
| G402A | 0.54 | 0.78 | 0.73 | 0.78 | 1.00 | 0.99 | 0.80 |
| GC403G | 0.59 | 0.79 | 0.67 | 0.79 | 0.99 | 1.00 | 0.80 |
| C403T | 0.58 | 0.85 | 0.64 | 0.80 | 1.00 | 1.00 | 0.82 |
| C403G | 0.47 | 0.76 | 0.93 | 0.80 | 0.97 | 0.98 | 0.83 |
| C403A | 0.54 | 0.83 | 0.81 | 0.81 | 1.00 | 0.99 | 0.85 |
| C404T | 0.63 | 0.77 | 0.72 | 0.81 | 0.99 | 0.99 | 0.82 |
| C404A | 0.62 | 0.91 | 0.70 | 0.86 | 0.99 | 1.00 | 0.87 |
| C405T | 0.61 | 0.87 | 0.68 | 0.84 | 0.99 | 1.00 | 0.85 |
| C405A | 0.64 | 0.89 | 0.68 | 0.88 | 0.97 | 1.00 | 0.86 |
| C406T | 0.57 | 0.86 | 0.72 | 0.82 | 1.00 | 0.95 | 0.88 |
| C406G | 0.35 | 0.97 | 1.00 | 0.82 | 0.97 | 1.00 | 0.91 |
| C406A | 0.54 | 0.88 | 0.73 | 0.82 | 1.00 | 0.99 | 0.86 |
| C407T | 0.67 | 0.64 | 0.71 | 0.87 | 0.86 | 0.79 | 0.95 |
| C407G | 0.65 | 0.93 | 0.71 | 0.87 | 1.00 | 1.00 | 0.89 |
| C407A | 0.65 | 0.65 | 0.80 | 0.97 | 0.78 | 0.81 | 0.95 |
| A408T | 0.57 | 0.71 | 0.72 | 0.77 | 0.97 | 0.94 | 0.81 |
| A408G | 0.61 | 0.87 | 0.69 | 0.84 | 0.99 | 1.00 | 0.85 |
| A408C | 0.62 | 0.74 | 0.71 | 0.80 | 0.98 | 0.96 | 0.82 |
| T409G | 0.65 | 0.82 | 0.76 | 0.84 | 1.00 | 0.99 | 0.87 |
| T409C | 0.62 | 0.84 | 0.71 | 0.83 | 1.00 | 1.00 | 0.85 |
| T409A | 0.63 | 0.85 | 0.75 | 0.84 | 1.00 | 0.95 | 0.90 |
| TC410T | 0.66 | 0.71 | 0.71 | 0.90 | 0.88 | 0.96 | 0.83 |
| C410T | 0.50 | 0.79 | 0.78 | 0.78 | 0.99 | 1.00 | 0.81 |
| C410A | 0.56 | 0.79 | 0.79 | 0.81 | 0.99 | 0.99 | 0.84 |
| C411T | 0.61 | 0.92 | 0.77 | 0.87 | 1.00 | 1.00 | 0.89 |
| C411A | 0.55 | 0.94 | 0.71 | 0.84 | 1.00 | 1.00 | 0.87 |
| C412T | 0.48 | 0.60 | 0.88 | 0.88 | 0.76 | 0.78 | 0.89 |
| C412A | 0.83 | 0.75 | 0.69 | 0.95 | 0.89 | 0.89 | 0.95 |
| CT413C | 0.58 | 0.84 | 0.76 | 0.83 | 0.99 | 1.00 | 0.85 |
| T413G | 0.64 | 0.43 | 1.00 | 0.91 | 0.26 | 0.30 | 0.84 |
| T413C | 0.65 | 0.73 | 0.74 | 0.84 | 0.96 | 0.94 | 0.86 |
| T413A | 0.65 | 0.73 | 0.71 | 0.80 | 0.99 | 0.98 | 0.81 |
| G414T | 0.55 | 0.70 | 0.84 | 0.86 | 0.88 | 0.89 | 0.88 |
| G414C | 0.61 | 0.81 | 0.69 | 0.80 | 1.00 | 1.00 | 0.81 |
| G414A | 0.47 | 0.62 | 0.88 | 0.85 | 0.80 | 0.80 | 0.88 |
| C415T | 0.68 | 0.26 | 0.91 | 0.84 | 0.11 | 0.12 | 0.85 |
| C415G | 0.67 | 0.54 | 0.91 | 1.00 | 0.69 | 0.70 | 1.00 |
| C415A | 0.64 | 0.87 | 0.76 | 0.86 | 1.00 | 1.00 | 0.88 |
| A416T | 0.61 | 0.82 | 0.68 | 0.81 | 1.00 | 0.99 | 0.83 |
| A416G | 0.63 | 0.70 | 0.73 | 0.86 | 0.90 | 0.92 | 0.85 |
| A416C | 0.74 | 0.37 | 0.78 | 0.77 | 0.19 | 0.17 | 0.89 |
| G417T | 0.60 | 0.82 | 0.78 | 0.83 | 0.99 | 0.99 | 0.86 |
| G417C | 0.54 | 0.83 | 0.72 | 0.79 | 1.00 | 1.00 | 0.82 |
| G417A | 0.56 | 0.80 | 0.75 | 0.80 | 0.99 | 0.99 | 0.83 |
| C418T | 0.62 | 0.56 | 0.80 | 0.93 | 0.73 | 0.72 | 0.95 |

| | | | | | | | |
|---|---|---|---|---|---|---|---|
| C418A | 0.44 | 0.59 | 0.85 | 0.93 | 0.61 | 0.80 | 0.75 |
| CT419C | 0.42 | 0.81 | 0.66 | 0.69 | 0.99 | 0.98 | 0.74 |
| T419G | 0.59 | 0.77 | 0.71 | 0.80 | 0.98 | 1.00 | 0.80 |
| T419C | 0.60 | 0.87 | 0.76 | 0.84 | 1.00 | 1.00 | 0.87 |
| T419A | 0.62 | 0.85 | 0.72 | 0.84 | 1.00 | 1.00 | 0.85 |
| T420G | 0.73 | 0.65 | 0.77 | 0.99 | 0.79 | 0.80 | 0.99 |
| T420C | 0.63 | 0.87 | 0.69 | 0.84 | 1.00 | 1.00 | 0.85 |
| T420A | 0.78 | 0.28 | 0.91 | 0.85 | 0.14 | 0.13 | 0.90 |
| G421T | 0.61 | 0.68 | 0.68 | 0.78 | 0.96 | 0.95 | 0.79 |
| G422T | 0.58 | 0.81 | 0.82 | 0.82 | 1.00 | 0.97 | 0.88 |
| G422C | 0.49 | 0.77 | 0.78 | 0.76 | 0.99 | 0.99 | 0.80 |
| GC423G | 0.69 | 0.56 | 0.71 | 0.91 | 0.73 | 0.71 | 0.94 |
| C423T | 0.65 | 0.61 | 0.71 | 0.97 | 0.74 | 0.85 | 0.85 |
| C423G | 0.62 | 0.40 | 0.80 | 0.67 | 0.22 | 0.17 | 0.95 |
| C424T | 0.65 | 0.42 | 0.86 | 0.76 | 0.25 | 0.22 | 0.91 |
| A425G | 0.63 | 0.43 | 0.78 | 0.75 | 0.23 | 0.22 | 0.84 |
| A425C | 0.61 | 0.82 | 0.73 | 0.82 | 1.00 | 0.97 | 0.86 |
| G427T | 0.67 | 0.37 | 0.88 | 0.80 | 0.19 | 0.18 | 0.87 |
| G427C | 0.49 | 0.60 | 0.88 | 0.86 | 0.79 | 0.77 | 0.92 |
| G427A | 0.75 | 0.50 | 0.62 | 0.79 | 0.33 | 0.32 | 0.81 |
| GA428G | 0.68 | 0.53 | 0.80 | 0.98 | 0.69 | 0.69 | 0.98 |
| A428G | 0.73 | 0.25 | 0.87 | 0.88 | 0.10 | 0.12 | 0.82 |
| A428C | 0.72 | 0.36 | 0.84 | 0.77 | 0.21 | 0.17 | 0.94 |
| T429C | 0.64 | 0.85 | 0.71 | 0.84 | 1.00 | 1.00 | 0.85 |
| T429A | 0.64 | 0.72 | 0.71 | 0.82 | 0.96 | 0.96 | 0.82 |
| G430A | 0.64 | 0.46 | 0.73 | 0.87 | 0.23 | 0.29 | 0.72 |
| G431T | 0.64 | 0.80 | 0.70 | 0.83 | 0.98 | 0.99 | 0.84 |
| G431C | 0.66 | 0.81 | 0.65 | 0.82 | 0.99 | 1.00 | 0.82 |
| G431A | 0.64 | 0.91 | 0.76 | 0.87 | 1.00 | 1.00 | 0.89 |
| T432G | 0.83 | 0.61 | 0.72 | 1.00 | 0.77 | 0.77 | 1.00 |
| T432C | 0.62 | 0.86 | 0.73 | 0.84 | 1.00 | 0.97 | 0.88 |
| T432A | 0.62 | 0.93 | 0.73 | 0.86 | 1.00 | 1.00 | 0.88 |
| G433C | 0.62 | 0.78 | 0.66 | 0.81 | 0.98 | 0.99 | 0.80 |
| G433A | 0.62 | 0.75 | 0.67 | 0.79 | 0.99 | 0.99 | 0.79 |
| GC434G | 0.62 | 0.75 | 0.72 | 0.82 | 0.97 | 0.98 | 0.82 |
| C434G | 0.77 | 0.48 | 0.55 | 0.83 | 0.25 | 0.28 | 0.71 |
| C434A | 0.67 | 0.89 | 0.76 | 0.87 | 1.00 | 0.99 | 0.90 |
| C435A | 0.53 | 0.90 | 0.82 | 0.86 | 0.98 | 1.00 | 0.87 |
| C436A | 0.61 | 0.87 | 0.70 | 0.84 | 1.00 | 1.00 | 0.85 |
| C437T | 0.61 | 0.74 | 0.75 | 0.79 | 1.00 | 0.92 | 0.88 |
| C437G | 0.58 | 0.87 | 0.71 | 0.83 | 1.00 | 1.00 | 0.84 |
| C437A | 0.62 | 0.79 | 0.62 | 0.79 | 1.00 | 1.00 | 0.79 |
| CA438C | 0.77 | 0.31 | 0.52 | 0.79 | 0.13 | 0.13 | 0.70 |
| A438T | 0.68 | 0.71 | 0.71 | 0.94 | 0.85 | 0.97 | 0.82 |
| A438G | 0.61 | 0.73 | 0.73 | 0.81 | 0.96 | 0.96 | 0.83 |
| A438C | 0.52 | 0.63 | 0.88 | 0.92 | 0.77 | 0.81 | 0.91 |
| A438AT | 0.65 | 0.67 | 0.61 | 0.77 | 0.94 | 0.95 | 0.77 |
| AT439A | 0.64 | 0.70 | 0.67 | 0.79 | 0.97 | 0.96 | 0.80 |
| T439C | 0.64 | 0.69 | 0.76 | 0.89 | 0.88 | 0.85 | 0.93 |
| T439A | 0.67 | 0.88 | 0.76 | 0.87 | 1.00 | 1.00 | 0.88 |
| T440G | 0.70 | 0.77 | 0.76 | 0.84 | 1.00 | 0.97 | 0.88 |
| T440C | 0.71 | 0.84 | 0.72 | 0.92 | 0.93 | 1.00 | 0.86 |
| T440A | 0.64 | 0.73 | 0.64 | 0.78 | 0.99 | 0.97 | 0.79 |

| | | | | | | | |
|---|---|---|---|---|---|---|---|
| G441T | 0.63 | 0.80 | 0.63 | 0.79 | 1.00 | 1.00 | 0.80 |
| G441C | 0.66 | 0.75 | 0.64 | 0.79 | 0.99 | 0.99 | 0.80 |
| G441A | 0.63 | 0.83 | 0.66 | 0.82 | 1.00 | 0.94 | 0.88 |
| C442T | 0.68 | 0.18 | 0.95 | 0.93 | 0.07 | 0.09 | 0.79 |
| C442G | 0.82 | 0.26 | 0.88 | 0.87 | 0.11 | 0.11 | 0.88 |
| C442A | 0.67 | 0.87 | 0.76 | 0.87 | 1.00 | 1.00 | 0.88 |
| A443T | 0.91 | 0.05 | 0.97 | 0.93 | 0.02 | 0.02 | 0.93 |
| A443G | 0.96 | 0.03 | 0.96 | 0.95 | 0.01 | 0.01 | 0.93 |
| A443C | 0.99 | 0.04 | 0.99 | 0.98 | 0.02 | 0.01 | 0.92 |
| AG444A | 0.99 | 0.01 | 0.98 | 0.92 | 0.00 | 0.00 | 0.99 |
| G444T | 0.91 | 0.01 | 0.97 | 0.96 | 0.01 | 0.01 | 0.91 |
| G444C | 1.00 | 0.00 | 1.00 | 1.00 | 0.00 | 0.00 | 0.96 |
| G444A | 0.95 | 0.02 | 0.99 | 0.94 | 0.01 | 0.01 | 0.95 |
| G445C | 0.94 | 0.02 | 0.86 | 0.97 | 0.01 | 0.01 | 0.86 |
| G445A | 0.96 | 0.01 | 0.95 | 0.95 | 0.00 | 0.00 | 0.94 |
| T446C | 0.95 | 0.01 | 0.94 | 0.88 | 0.01 | 0.00 | 1.00 |
| T446A | 0.96 | 0.02 | 0.79 | 0.90 | 0.01 | 0.01 | 0.90 |
| A447T | 0.96 | 0.02 | 0.97 | 0.96 | 0.01 | 0.01 | 0.93 |
| A447G | 0.97 | 0.03 | 0.92 | 0.90 | 0.01 | 0.01 | 0.96 |
| G448A | 0.68 | 0.97 | 0.78 | 0.90 | 1.00 | 1.00 | 0.92 |
| G449T | 0.76 | 0.10 | 0.98 | 0.87 | 0.04 | 0.04 | 0.91 |
| G449A | 0.90 | 0.08 | 0.84 | 0.85 | 0.04 | 0.03 | 0.94 |
| C450T | 0.62 | 0.89 | 0.81 | 0.86 | 1.00 | 0.96 | 0.92 |
| C450A | 0.64 | 0.85 | 0.72 | 0.84 | 1.00 | 1.00 | 0.86 |
| A451T | 0.60 | 0.85 | 0.70 | 0.83 | 1.00 | 1.00 | 0.84 |
| A451G | 0.60 | 0.67 | 0.81 | 0.90 | 0.84 | 0.83 | 0.93 |
| A451C | 0.68 | 0.84 | 0.76 | 0.85 | 1.00 | 0.98 | 0.87 |
| G452T | 0.58 | 0.82 | 0.70 | 0.80 | 1.00 | 1.00 | 0.81 |
| G452A | 0.65 | 0.85 | 0.70 | 0.85 | 1.00 | 1.00 | 0.86 |
| GC453G | 0.67 | 0.60 | 0.53 | 0.73 | 0.80 | 0.71 | 0.81 |
| C453T | 0.65 | 0.74 | 0.72 | 0.83 | 0.96 | 0.97 | 0.83 |
| C453G | 0.59 | 0.72 | 0.69 | 0.79 | 0.96 | 0.97 | 0.79 |
| C453A | 0.69 | 0.72 | 0.64 | 0.82 | 0.95 | 0.98 | 0.80 |
| C454T | 0.53 | 0.66 | 0.83 | 0.87 | 0.84 | 0.86 | 0.88 |
| C454G | 0.65 | 0.61 | 0.80 | 0.97 | 0.76 | 0.78 | 0.96 |
| C454A | 0.63 | 0.75 | 0.70 | 0.80 | 0.99 | 0.98 | 0.82 |
| C455T | 0.62 | 0.68 | 0.72 | 0.82 | 0.92 | 0.90 | 0.85 |
| C455CA | 0.53 | 0.84 | 0.76 | 0.80 | 1.00 | 0.98 | 0.85 |
| CA456C | 0.51 | 0.77 | 0.42 | 0.60 | 0.49 | 0.55 | 0.52 |
| A456T | 0.64 | 0.80 | 0.76 | 0.84 | 0.99 | 0.98 | 0.86 |
| A456G | 0.62 | 0.81 | 0.67 | 0.81 | 1.00 | 1.00 | 0.82 |
| A456C | 0.61 | 0.88 | 0.72 | 0.84 | 1.00 | 1.00 | 0.85 |
| G457T | 0.61 | 0.84 | 0.74 | 0.83 | 1.00 | 1.00 | 0.85 |
| G457C | 0.58 | 0.63 | 0.84 | 0.92 | 0.80 | 0.78 | 0.96 |
| G457A | 0.64 | 0.84 | 0.71 | 0.84 | 1.00 | 0.99 | 0.85 |
| C458T | 0.63 | 0.84 | 0.75 | 0.84 | 1.00 | 0.97 | 0.88 |
| C458G | 0.62 | 0.79 | 0.71 | 0.81 | 0.98 | 1.00 | 0.81 |
| C458A | 0.61 | 0.85 | 0.67 | 0.82 | 1.00 | 1.00 | 0.83 |
| T459G | 0.53 | 0.87 | 0.82 | 0.83 | 1.00 | 0.99 | 0.87 |
| T459C | 0.58 | 0.78 | 0.72 | 0.80 | 0.99 | 0.99 | 0.82 |
| T459A | 0.67 | 0.64 | 0.80 | 0.98 | 0.78 | 0.79 | 0.98 |
| TG460T | 0.77 | 0.30 | 0.87 | 0.89 | 0.13 | 0.14 | 0.83 |
| G460T | 0.72 | 0.55 | 0.70 | 0.94 | 0.72 | 0.71 | 0.96 |

| | | | | | | | |
|---|---|---|---|---|---|---|---|
| G460C | 0.76 | 0.14 | 0.60 | 0.90 | 0.06 | 0.07 | 0.67 |
| G460A | 0.69 | 0.20 | 0.93 | 0.89 | 0.08 | 0.09 | 0.82 |
| G461T | 0.62 | 0.54 | 0.85 | 0.96 | 0.70 | 0.69 | 0.99 |
| G461C | 0.85 | 0.08 | 0.98 | 0.93 | 0.03 | 0.08 | 0.93 |
| G461A | 0.60 | 0.50 | 0.90 | 0.93 | 0.50 | 0.54 | 0.89 |
| A462T | 0.60 | 0.75 | 0.84 | 0.90 | 0.90 | 0.97 | 0.86 |
| A462G | 0.59 | 0.68 | 0.75 | 0.83 | 0.90 | 0.89 | 0.86 |
| A462C | 0.88 | 0.51 | 0.95 | 1.00 | 0.72 | 0.73 | 1.00 |
| AC463A | 0.72 | 0.63 | 0.76 | 0.98 | 0.78 | 0.77 | 1.00 |
| C463T | 0.56 | 0.83 | 0.73 | 0.81 | 0.99 | 1.00 | 0.83 |
| C464T | 0.57 | 0.83 | 0.75 | 0.81 | 1.00 | 0.99 | 0.84 |
| C464A | 0.62 | 0.82 | 0.72 | 0.82 | 1.00 | 0.98 | 0.85 |
| CT465C | 0.69 | 0.60 | 0.69 | 0.90 | 0.79 | 0.75 | 0.96 |
| T465G | 0.66 | 0.82 | 0.68 | 0.83 | 1.00 | 0.99 | 0.84 |
| T465C | 0.64 | 0.69 | 0.76 | 0.91 | 0.85 | 0.89 | 0.88 |
| T465A | 0.57 | 0.73 | 0.71 | 0.78 | 0.97 | 0.98 | 0.78 |
| TC466T | 0.63 | 0.63 | 0.65 | 0.80 | 0.89 | 0.87 | 0.82 |
| C466T | 0.57 | 0.54 | 0.90 | 0.97 | 0.69 | 0.70 | 0.97 |
| C466G | 0.92 | 0.15 | 0.76 | 0.85 | 0.06 | 0.06 | 0.90 |
| C466A | 0.71 | 0.25 | 0.79 | 0.77 | 0.11 | 0.10 | 0.87 |
| C467T | 0.80 | 0.32 | 0.79 | 0.83 | 0.15 | 0.15 | 0.86 |
| C467G | 0.99 | 0.17 | 0.51 | 0.82 | 0.09 | 0.08 | 0.79 |
| C467A | 0.66 | 0.27 | 0.74 | 0.78 | 0.11 | 0.11 | 0.79 |
| C468T | 0.63 | 0.75 | 0.76 | 0.82 | 0.98 | 0.99 | 0.83 |
| CT469C | 0.58 | 0.82 | 0.61 | 0.78 | 1.00 | 1.00 | 0.78 |
| T469G | 0.62 | 0.82 | 0.62 | 0.80 | 0.99 | 1.00 | 0.81 |
| T469C | 0.55 | 0.75 | 0.77 | 0.79 | 0.98 | 0.97 | 0.82 |
| T469A | 0.66 | 0.70 | 0.69 | 0.85 | 0.91 | 0.93 | 0.84 |
| T469TG | 0.68 | 0.70 | 0.84 | 0.98 | 0.82 | 0.83 | 0.99 |
| G470A | 0.61 | 0.60 | 0.56 | 0.67 | 0.50 | 0.42 | 0.80 |
| G471T | 0.64 | 0.52 | 0.66 | 0.69 | 0.36 | 0.30 | 0.87 |
| G471C | 0.59 | 0.53 | 0.69 | 0.73 | 0.34 | 0.32 | 0.80 |
| G471A | 0.61 | 0.59 | 0.63 | 0.83 | 0.69 | 0.74 | 0.78 |
| G472A | 0.59 | 0.66 | 0.55 | 0.69 | 0.89 | 0.87 | 0.71 |
| GA473G | 0.45 | 0.71 | 0.86 | 0.75 | 0.97 | 0.91 | 0.84 |
| A473T | 0.58 | 0.79 | 0.64 | 0.78 | 0.99 | 0.99 | 0.80 |
| A473G | 0.61 | 0.74 | 0.78 | 0.83 | 0.96 | 0.98 | 0.84 |
| A473C | 0.65 | 0.51 | 0.69 | 0.74 | 0.35 | 0.31 | 0.85 |
| A474T | 0.61 | 0.78 | 0.76 | 0.82 | 0.98 | 1.00 | 0.83 |
| A474G | 0.56 | 0.78 | 0.73 | 0.79 | 0.99 | 1.00 | 0.81 |
| A474C | 0.50 | 0.70 | 0.88 | 0.83 | 0.91 | 0.89 | 0.88 |
| A475T | 0.56 | 0.77 | 0.67 | 0.77 | 0.99 | 1.00 | 0.77 |
| A475G | 0.59 | 0.80 | 0.70 | 0.80 | 1.00 | 0.99 | 0.82 |
| C476T | 0.61 | 0.84 | 0.70 | 0.82 | 1.00 | 1.00 | 0.84 |
| C476G | 0.58 | 0.82 | 0.79 | 0.82 | 1.00 | 0.99 | 0.85 |
| C476A | 0.67 | 0.78 | 0.85 | 0.98 | 0.85 | 0.88 | 0.96 |
| C476CA | 0.72 | 0.76 | 0.77 | 0.92 | 0.91 | 0.91 | 0.93 |
| A477T | 0.59 | 0.79 | 0.69 | 0.80 | 1.00 | 1.00 | 0.81 |
| A477G | 0.57 | 0.81 | 0.78 | 0.81 | 1.00 | 1.00 | 0.83 |
| AC478A | 0.71 | 0.75 | 0.75 | 0.95 | 0.87 | 0.94 | 0.88 |
| C478T | 0.60 | 0.80 | 0.71 | 0.81 | 0.99 | 1.00 | 0.83 |
| C478G | 0.65 | 0.70 | 0.92 | 1.00 | 0.82 | 0.84 | 0.99 |
| C478A | 0.64 | 0.82 | 0.79 | 0.88 | 0.95 | 1.00 | 0.86 |

| | | | | | | | |
|---|---|---|---|---|---|---|---|
| G479A | 0.61 | 0.61 | 0.63 | 0.82 | 0.73 | 0.76 | 0.79 |
| G481C | 0.57 | 0.83 | 0.43 | 0.67 | 0.94 | 0.92 | 0.66 |
| C482T | 0.68 | 0.82 | 0.73 | 0.85 | 1.00 | 1.00 | 0.86 |
| C482G | 0.48 | 0.74 | 0.63 | 0.66 | 0.99 | 1.00 | 0.67 |
| C482A | 0.74 | 0.84 | 0.87 | 0.92 | 0.96 | 0.91 | 0.98 |
| A483T | 0.59 | 0.88 | 0.69 | 0.83 | 1.00 | 1.00 | 0.85 |
| A483G | 0.57 | 0.86 | 0.85 | 0.86 | 0.97 | 1.00 | 0.87 |
| A483C | 0.88 | 0.98 | 0.44 | 0.89 | 1.00 | 1.00 | 0.85 |
| G484T | 0.62 | 0.82 | 0.63 | 0.81 | 0.99 | 1.00 | 0.81 |
| G484C | 0.67 | 0.87 | 0.44 | 0.85 | 0.93 | 1.00 | 0.75 |
| G484A | 0.57 | 0.81 | 0.75 | 0.80 | 1.00 | 0.99 | 0.83 |
| A485T | 0.63 | 0.77 | 0.78 | 0.83 | 0.99 | 0.98 | 0.85 |
| A485G | 0.60 | 0.78 | 0.90 | 0.92 | 0.91 | 0.90 | 0.95 |
| A485C | 0.57 | 0.82 | 0.63 | 0.78 | 1.00 | 1.00 | 0.79 |
| G486T | 0.61 | 0.73 | 0.87 | 0.95 | 0.85 | 0.87 | 0.96 |
| G486A | 0.55 | 0.80 | 0.63 | 0.76 | 1.00 | 1.00 | 0.78 |
| G487T | 0.54 | 0.85 | 0.63 | 0.78 | 1.00 | 1.00 | 0.80 |
| G487A | 0.51 | 0.81 | 0.52 | 0.71 | 0.96 | 1.00 | 0.69 |
| G488T | 0.49 | 0.82 | 0.79 | 0.78 | 1.00 | 1.00 | 0.81 |
| G488C | 0.54 | 0.79 | 0.58 | 0.73 | 0.99 | 0.98 | 0.74 |
| C489T | 0.64 | 0.76 | 0.84 | 0.92 | 0.90 | 0.92 | 0.92 |
| C490T | 0.64 | 0.75 | 0.87 | 0.98 | 0.85 | 0.90 | 0.94 |
| C490A | 0.60 | 0.80 | 0.79 | 0.83 | 0.99 | 0.98 | 0.85 |
| T491C | 0.63 | 0.77 | 0.66 | 0.80 | 0.99 | 0.99 | 0.80 |
| T491A | 0.67 | 0.79 | 0.68 | 0.82 | 1.00 | 1.00 | 0.83 |
| A492T | 0.62 | 0.76 | 0.76 | 0.81 | 0.99 | 0.97 | 0.84 |
| A492G | 0.60 | 0.81 | 0.68 | 0.80 | 0.99 | 0.99 | 0.82 |
| A492C | 0.66 | 0.82 | 0.77 | 0.85 | 0.99 | 1.00 | 0.86 |
| C493T | 0.62 | 0.79 | 0.71 | 0.81 | 1.00 | 0.95 | 0.86 |
| C493G | 0.66 | 0.80 | 0.73 | 0.84 | 0.99 | 0.98 | 0.85 |
| C493A | 0.65 | 0.78 | 0.79 | 0.87 | 0.96 | 0.97 | 0.87 |
| A494T | 0.64 | 0.81 | 0.64 | 0.81 | 1.00 | 0.99 | 0.82 |
| A494G | 0.58 | 0.81 | 0.79 | 0.82 | 1.00 | 0.99 | 0.85 |
| A494C | 0.63 | 0.75 | 0.76 | 0.81 | 1.00 | 0.95 | 0.86 |
| G495T | 0.59 | 0.77 | 0.69 | 0.79 | 0.99 | 0.99 | 0.80 |
| G495A | 0.70 | 0.80 | 0.74 | 0.85 | 1.00 | 0.98 | 0.87 |
| G496T | 0.60 | 0.77 | 0.62 | 0.77 | 1.00 | 0.99 | 0.78 |
| G496A | 0.56 | 0.78 | 0.82 | 0.81 | 0.99 | 0.98 | 0.85 |
| C497T | 0.70 | 0.84 | 0.44 | 0.77 | 1.00 | 0.99 | 0.75 |
| C497G | 0.78 | 0.81 | 0.36 | 0.75 | 0.98 | 1.00 | 0.70 |
| C497A | 0.61 | 0.84 | 0.71 | 0.83 | 0.99 | 1.00 | 0.83 |
| T498G | 0.59 | 0.85 | 0.64 | 0.80 | 1.00 | 0.99 | 0.82 |
| T498C | 0.57 | 0.75 | 0.55 | 0.73 | 0.97 | 0.97 | 0.72 |
| T498A | 0.62 | 0.82 | 0.73 | 0.83 | 0.99 | 1.00 | 0.84 |
| G499T | 0.54 | 0.80 | 0.64 | 0.79 | 0.96 | 1.00 | 0.77 |
| G499A | 0.58 | 0.71 | 0.74 | 0.80 | 0.95 | 0.96 | 0.81 |
| G500T | 0.63 | 0.75 | 0.83 | 0.88 | 0.93 | 0.93 | 0.90 |
| G500C | 0.60 | 0.84 | 0.68 | 0.82 | 1.00 | 0.99 | 0.83 |
| G500A | 0.60 | 0.63 | 0.92 | 0.99 | 0.77 | 0.78 | 0.99 |
| G501T | 0.56 | 0.76 | 0.94 | 0.93 | 0.87 | 0.89 | 0.95 |
| G501A | 0.62 | 0.75 | 0.83 | 0.89 | 0.92 | 0.91 | 0.92 |
| C502T | 0.60 | 0.79 | 0.97 | 0.97 | 0.86 | 0.89 | 0.98 |
| C502A | 0.60 | 0.80 | 0.99 | 0.96 | 0.88 | 0.89 | 0.99 |

| | | | | | | | |
|---|---|---|---|---|---|---|---|
| G505T | 0.59 | 0.84 | 0.56 | 0.78 | 0.99 | 1.00 | 0.78 |
| G505A | 0.54 | 0.78 | 0.85 | 0.80 | 1.00 | 0.99 | 0.84 |
| G507T | 0.65 | 0.75 | 0.98 | 1.00 | 0.84 | 0.87 | 1.00 |
| G507C | 0.70 | 0.68 | 0.98 | 1.00 | 0.82 | 0.84 | 1.00 |
| G507A | 0.57 | 0.69 | 0.94 | 0.98 | 0.80 | 0.83 | 0.98 |
| T508C | 0.65 | 0.81 | 0.62 | 0.81 | 0.99 | 0.99 | 0.81 |
| T508A | 0.64 | 0.77 | 0.61 | 0.78 | 1.00 | 0.99 | 0.79 |
| T509C | 0.60 | 0.75 | 0.67 | 0.80 | 0.97 | 1.00 | 0.78 |
| T509A | 0.63 | 0.80 | 0.60 | 0.79 | 1.00 | 0.99 | 0.80 |
| G510T | 0.64 | 0.83 | 1.00 | 1.00 | 0.86 | 0.91 | 1.00 |
| G510C | 0.64 | 0.82 | 0.56 | 0.78 | 1.00 | 0.99 | 0.78 |
| G510A | 0.64 | 0.79 | 0.76 | 0.85 | 0.97 | 1.00 | 0.84 |
| GC511G | 0.61 | 0.87 | 0.61 | 0.82 | 1.00 | 1.00 | 0.83 |
| C511T | 0.61 | 0.79 | 0.95 | 0.97 | 0.87 | 0.91 | 0.96 |
| C511G | 0.65 | 0.79 | 0.55 | 0.77 | 0.99 | 0.99 | 0.77 |
| C511A | 0.63 | 0.80 | 0.81 | 0.84 | 0.99 | 0.98 | 0.87 |
| C512T | 0.67 | 0.81 | 0.79 | 0.86 | 0.98 | 0.99 | 0.87 |
| C512A | 0.59 | 0.79 | 0.78 | 0.82 | 1.00 | 0.99 | 0.84 |
| A513T | 0.60 | 0.75 | 0.88 | 0.91 | 0.89 | 0.90 | 0.93 |
| A513G | 0.64 | 0.80 | 0.64 | 0.80 | 1.00 | 1.00 | 0.80 |
| C514T | 0.63 | 0.76 | 0.94 | 0.99 | 0.84 | 0.87 | 0.99 |
| C514A | 0.58 | 0.70 | 0.63 | 0.78 | 0.92 | 0.91 | 0.80 |
| C515T | 0.66 | 0.70 | 0.98 | 1.00 | 0.82 | 0.84 | 1.00 |
| C515G | 0.92 | 0.12 | 0.54 | 0.92 | 0.04 | 0.05 | 0.69 |
| C515A | 0.59 | 0.74 | 0.72 | 0.79 | 0.98 | 0.97 | 0.81 |
| T516C | 0.60 | 0.80 | 0.67 | 0.79 | 0.99 | 1.00 | 0.80 |
| G517T | 0.63 | 0.67 | 1.00 | 1.00 | 0.77 | 0.83 | 1.00 |
| G517C | 0.61 | 0.69 | 0.97 | 1.00 | 0.80 | 0.83 | 1.00 |
| G517A | 0.60 | 0.77 | 0.82 | 0.84 | 0.97 | 0.99 | 0.84 |
| G517GC | 0.62 | 0.74 | 0.92 | 0.97 | 0.84 | 0.86 | 0.99 |
| C518T | 0.57 | 0.75 | 0.90 | 0.93 | 0.87 | 0.93 | 0.90 |
| C519T | 0.51 | 0.82 | 0.60 | 0.73 | 1.00 | 0.99 | 0.75 |
| C520T | 0.56 | 0.78 | 0.78 | 0.80 | 0.99 | 0.98 | 0.83 |
| C520A | 0.57 | 0.79 | 0.53 | 0.72 | 0.99 | 1.00 | 0.72 |
| C521T | 0.61 | 0.82 | 0.71 | 0.83 | 0.98 | 1.00 | 0.83 |
| C521G | 0.57 | 0.77 | 0.87 | 0.90 | 0.90 | 0.96 | 0.86 |
| C521A | 0.60 | 0.73 | 0.99 | 1.00 | 0.81 | 0.85 | 1.00 |
| C522T | 0.58 | 0.78 | 0.88 | 0.85 | 0.96 | 0.97 | 0.87 |
| C527T | 0.56 | 0.82 | 0.56 | 0.75 | 0.99 | 0.99 | 0.75 |
| C527G | 0.66 | 0.83 | 0.28 | 0.46 | 0.98 | 0.75 | 0.54 |
| C527A | 0.63 | 0.87 | 0.50 | 0.79 | 1.00 | 1.00 | 0.77 |
| T528G | 0.60 | 0.71 | 0.64 | 0.78 | 0.95 | 0.99 | 0.75 |
| T528C | 0.60 | 0.79 | 0.70 | 0.80 | 1.00 | 0.99 | 0.82 |
| T528A | 0.51 | 0.80 | 0.48 | 0.63 | 0.92 | 0.93 | 0.63 |
| G529T | 0.63 | 0.79 | 0.73 | 0.82 | 1.00 | 0.94 | 0.89 |
| G529C | 0.74 | 0.84 | 0.86 | 0.93 | 0.96 | 0.98 | 0.92 |
| G529A | 0.57 | 0.74 | 0.68 | 0.76 | 0.99 | 0.99 | 0.78 |
| C530T | 0.59 | 0.83 | 0.54 | 0.76 | 0.99 | 1.00 | 0.76 |
| G531T | 0.68 | 0.71 | 0.39 | 0.79 | 0.64 | 0.89 | 0.53 |
| G531C | 0.40 | 0.83 | 1.00 | 0.97 | 0.77 | 0.89 | 0.98 |
| G531A | 0.68 | 0.77 | 0.68 | 0.82 | 1.00 | 0.99 | 0.83 |
| T532C | 0.62 | 0.70 | 0.93 | 0.99 | 0.81 | 0.83 | 1.00 |
| T532A | 0.51 | 0.70 | 0.92 | 0.91 | 0.84 | 0.88 | 0.91 |

| | | | | | | | |
|---|---|---|---|---|---|---|---|
| A533T | 0.63 | 0.72 | 0.85 | 0.93 | 0.87 | 0.83 | 0.99 |
| A533G | 0.55 | 0.70 | 0.91 | 0.94 | 0.83 | 0.86 | 0.94 |
| G534T | 0.68 | 0.82 | 0.61 | 0.83 | 1.00 | 1.00 | 0.83 |
| G534C | 0.62 | 0.76 | 0.68 | 0.80 | 0.98 | 0.98 | 0.80 |
| G534A | 0.66 | 0.83 | 0.63 | 0.82 | 1.00 | 1.00 | 0.82 |
| A535T | 0.57 | 0.80 | 0.51 | 0.74 | 0.95 | 0.97 | 0.72 |
| A535G | 0.60 | 0.83 | 0.72 | 0.82 | 1.00 | 0.99 | 0.85 |
| T536C | 0.66 | 0.77 | 0.72 | 0.83 | 0.98 | 0.98 | 0.84 |
| T536A | 0.65 | 0.85 | 0.59 | 0.82 | 1.00 | 1.00 | 0.82 |
| TG537T | 0.77 | 0.80 | 0.77 | 0.93 | 0.93 | 0.93 | 0.93 |
| G537T | 0.59 | 0.82 | 0.49 | 0.75 | 0.98 | 0.99 | 0.74 |
| G537C | 0.55 | 0.78 | 0.83 | 0.81 | 0.99 | 0.98 | 0.85 |
| G537A | 0.49 | 0.90 | 0.35 | 0.39 | 1.00 | 0.79 | 0.47 |
| G538T | 0.64 | 0.84 | 0.60 | 0.81 | 1.00 | 1.00 | 0.81 |
| G538C | 0.60 | 0.76 | 0.76 | 0.82 | 0.97 | 0.99 | 0.83 |
| G538A | 0.59 | 0.80 | 0.63 | 0.80 | 0.97 | 1.00 | 0.79 |
| T539G | 0.79 | 0.89 | 0.40 | 0.82 | 1.00 | 1.00 | 0.78 |
| T539C | 0.57 | 0.77 | 0.82 | 0.81 | 0.99 | 1.00 | 0.83 |
| T539A | 0.68 | 0.77 | 0.65 | 0.86 | 0.94 | 1.00 | 0.81 |
| G540T | 0.60 | 0.89 | 0.40 | 0.72 | 0.99 | 0.99 | 0.69 |
| G540A | 0.56 | 0.87 | 0.35 | 0.56 | 0.94 | 0.97 | 0.52 |
| A541T | 0.59 | 0.82 | 0.60 | 0.78 | 1.00 | 0.99 | 0.80 |
| A541G | 0.59 | 0.77 | 0.77 | 0.82 | 0.98 | 0.97 | 0.84 |
| A541C | 0.64 | 0.83 | 0.68 | 0.83 | 1.00 | 1.00 | 0.84 |
| A542G | 0.61 | 0.81 | 0.55 | 0.77 | 0.99 | 0.99 | 0.77 |
| T543G | 0.49 | 0.54 | 1.00 | 0.96 | 0.62 | 0.67 | 0.96 |
| T543C | 0.54 | 0.75 | 0.96 | 0.91 | 0.89 | 0.88 | 0.96 |
| T543A | 0.61 | 0.79 | 0.66 | 0.80 | 0.99 | 0.99 | 0.82 |
| G544T | 0.60 | 0.88 | 0.57 | 0.81 | 1.00 | 1.00 | 0.81 |
| G544C | 0.58 | 0.78 | 0.70 | 0.80 | 0.99 | 1.00 | 0.81 |
| G544A | 0.55 | 0.82 | 0.69 | 0.78 | 1.00 | 1.00 | 0.81 |
| T545G | 0.56 | 0.83 | 0.83 | 0.84 | 0.99 | 1.00 | 0.86 |
| T545C | 0.63 | 0.81 | 0.70 | 0.82 | 0.99 | 1.00 | 0.83 |
| T545A | 0.54 | 0.76 | 0.82 | 0.80 | 0.99 | 0.97 | 0.83 |
| C546T | 0.60 | 0.86 | 0.55 | 0.78 | 1.00 | 1.00 | 0.78 |
| C546A | 0.55 | 0.84 | 0.50 | 0.72 | 1.00 | 0.97 | 0.74 |
| A547T | 0.62 | 0.83 | 0.65 | 0.81 | 1.00 | 1.00 | 0.82 |
| A547G | 0.54 | 0.76 | 0.92 | 0.92 | 0.87 | 0.94 | 0.90 |
| A547C | 0.69 | 0.73 | 0.86 | 0.99 | 0.84 | 0.85 | 0.99 |
| T548C | 0.58 | 0.72 | 0.87 | 0.93 | 0.85 | 0.89 | 0.91 |
| T548A | 0.60 | 0.77 | 0.75 | 0.81 | 0.99 | 0.98 | 0.84 |
| A549G | 0.63 | 0.77 | 0.74 | 0.82 | 0.99 | 0.99 | 0.83 |
| A549C | 0.59 | 0.86 | 0.64 | 0.80 | 1.00 | 1.00 | 0.81 |
| AT550A | 0.73 | 0.75 | 0.63 | 0.82 | 0.99 | 0.98 | 0.82 |
| T550C | 0.63 | 0.80 | 0.69 | 0.82 | 0.99 | 0.99 | 0.83 |
| T550A | 0.58 | 0.84 | 0.59 | 0.79 | 1.00 | 1.00 | 0.79 |
| TC551T | 0.68 | 0.84 | 0.57 | 0.82 | 0.99 | 1.00 | 0.80 |
| C551T | 0.64 | 0.85 | 0.56 | 0.81 | 1.00 | 1.00 | 0.81 |
| C551G | 0.40 | 0.73 | 0.64 | 0.61 | 0.79 | 0.84 | 0.60 |
| C551A | 0.75 | 0.90 | 0.55 | 0.87 | 1.00 | 0.98 | 0.86 |
| C552G | 0.65 | 0.82 | 0.76 | 0.84 | 1.00 | 0.99 | 0.86 |
| C552A | 0.54 | 0.83 | 0.59 | 0.77 | 0.99 | 1.00 | 0.77 |
| T553G | 0.60 | 0.83 | 0.71 | 0.82 | 1.00 | 0.99 | 0.83 |

| | | | | | | | |
|---|---|---|---|---|---|---|---|
| T553C | 0.60 | 0.82 | 0.78 | 0.85 | 0.97 | 1.00 | 0.85 |
| T553A | 0.59 | 0.82 | 0.64 | 0.79 | 1.00 | 0.99 | 0.81 |
| TG554T | 0.70 | 0.73 | 0.92 | 1.00 | 0.84 | 0.85 | 1.00 |
| G555C | 0.56 | 0.76 | 0.95 | 0.96 | 0.84 | 0.88 | 0.96 |
| G555A | 0.57 | 0.76 | 0.90 | 0.91 | 0.89 | 0.93 | 0.91 |
| G556C | 0.62 | 0.79 | 0.89 | 0.89 | 0.94 | 0.95 | 0.91 |
| G556A | 0.58 | 0.69 | 0.97 | 0.99 | 0.80 | 0.83 | 0.99 |
| T557C | 0.60 | 0.75 | 0.87 | 0.89 | 0.91 | 0.89 | 0.94 |
| T557A | 0.61 | 0.69 | 0.96 | 1.00 | 0.80 | 0.83 | 1.00 |
| A558T | 0.67 | 0.76 | 0.73 | 0.83 | 0.98 | 0.97 | 0.85 |
| A558G | 0.62 | 0.75 | 0.72 | 0.80 | 0.99 | 0.99 | 0.80 |
| G559T | 0.67 | 0.84 | 0.61 | 0.83 | 1.00 | 0.99 | 0.83 |
| G559A | 0.61 | 0.79 | 0.74 | 0.81 | 1.00 | 0.99 | 0.84 |
| A560T | 0.60 | 0.83 | 0.68 | 0.81 | 1.00 | 1.00 | 0.82 |
| A560C | 0.52 | 0.89 | 0.73 | 0.81 | 1.00 | 1.00 | 0.84 |
| G561C | 0.63 | 0.77 | 0.70 | 0.80 | 1.00 | 0.99 | 0.82 |
| G561A | 0.59 | 0.85 | 0.67 | 0.81 | 1.00 | 0.99 | 0.83 |
| T562C | 0.61 | 0.75 | 0.83 | 0.86 | 0.94 | 0.93 | 0.89 |
| T562A | 0.63 | 0.82 | 0.75 | 0.83 | 1.00 | 0.99 | 0.86 |
| TG563T | 0.81 | 0.85 | 0.75 | 0.90 | 0.99 | 0.99 | 0.89 |
| G563T | 0.52 | 0.80 | 0.74 | 0.78 | 0.99 | 0.99 | 0.81 |
| G563C | 0.78 | 0.73 | 0.85 | 1.00 | 0.85 | 0.86 | 1.00 |
| G563A | 0.56 | 0.88 | 0.54 | 0.78 | 1.00 | 1.00 | 0.78 |
| G564C | 0.69 | 0.71 | 0.81 | 0.99 | 0.83 | 0.85 | 0.97 |
| G564A | 0.56 | 0.68 | 0.92 | 0.97 | 0.80 | 0.83 | 0.96 |
| T565G | 0.73 | 0.94 | 0.23 | 0.65 | 1.00 | 0.98 | 0.58 |
| T565C | 0.59 | 0.75 | 0.82 | 0.84 | 0.96 | 0.92 | 0.89 |
| T565A | 0.57 | 0.85 | 0.74 | 0.82 | 1.00 | 1.00 | 0.84 |
| G566T | 0.59 | 0.43 | 0.88 | 0.88 | 0.21 | 0.27 | 0.75 |
| G566A | 0.63 | 0.77 | 0.66 | 0.81 | 0.97 | 1.00 | 0.80 |
| GC567G | 0.75 | 0.89 | 0.48 | 0.85 | 1.00 | 1.00 | 0.81 |
| C567T | 0.61 | 0.81 | 0.63 | 0.79 | 1.00 | 1.00 | 0.80 |
| C567G | 0.63 | 0.92 | 0.25 | 0.51 | 0.81 | 0.92 | 0.43 |
| C567A | 0.61 | 0.54 | 0.70 | 0.79 | 0.64 | 0.57 | 0.90 |
| G568T | 0.61 | 0.87 | 0.75 | 0.84 | 1.00 | 1.00 | 0.87 |
| G568A | 0.63 | 0.80 | 0.70 | 0.82 | 0.99 | 0.99 | 0.83 |
| G569T | 0.60 | 0.78 | 0.83 | 0.84 | 0.98 | 0.97 | 0.87 |
| G569C | 0.55 | 0.68 | 0.81 | 0.84 | 0.89 | 0.88 | 0.87 |
| G569A | 0.55 | 0.72 | 0.85 | 0.86 | 0.90 | 0.91 | 0.88 |
| GC570G | 0.74 | 0.78 | 0.82 | 0.97 | 0.88 | 0.90 | 0.96 |
| C570T | 0.54 | 0.83 | 0.48 | 0.71 | 0.96 | 0.97 | 0.70 |
| C570G | 0.64 | 0.91 | 0.27 | 0.52 | 0.97 | 0.88 | 0.51 |
| C570A | 0.51 | 0.80 | 0.75 | 0.78 | 1.00 | 0.99 | 0.81 |
| C571T | 0.58 | 0.88 | 0.60 | 0.80 | 1.00 | 1.00 | 0.81 |
| C571G | 0.69 | 0.73 | 0.88 | 0.99 | 0.83 | 0.84 | 1.00 |
| C571A | 0.63 | 0.85 | 0.87 | 0.88 | 0.98 | 0.99 | 0.89 |
| CA572C | 0.68 | 0.66 | 0.97 | 1.00 | 0.80 | 0.81 | 1.00 |
| A572T | 0.57 | 0.81 | 0.81 | 0.81 | 1.00 | 0.99 | 0.84 |
| A572G | 0.63 | 0.82 | 0.61 | 0.80 | 1.00 | 1.00 | 0.80 |
| A572C | 0.56 | 0.69 | 0.90 | 0.95 | 0.81 | 0.85 | 0.94 |
| A572AG | 0.83 | 0.79 | 0.90 | 1.00 | 0.88 | 0.88 | 1.00 |
| G573T | 0.63 | 0.80 | 0.88 | 0.89 | 0.95 | 0.95 | 0.91 |
| G573A | 0.53 | 0.77 | 0.87 | 0.80 | 1.00 | 0.98 | 0.84 |

| | | | | | | | |
|---|---|---|---|---|---|---|---|
| G574T | 0.70 | 0.83 | 0.77 | 0.87 | 0.99 | 0.99 | 0.88 |
| G574C | 0.61 | 0.80 | 0.82 | 0.84 | 0.99 | 0.99 | 0.86 |
| G574A | 0.56 | 0.70 | 0.96 | 0.99 | 0.79 | 0.83 | 0.98 |
| G575T | 0.51 | 0.69 | 0.98 | 0.97 | 0.79 | 0.82 | 0.98 |
| G575C | 0.62 | 0.74 | 0.84 | 0.92 | 0.88 | 0.89 | 0.93 |
| G575A | 0.62 | 0.75 | 0.85 | 0.88 | 0.93 | 0.86 | 0.98 |
| GC576G | 0.86 | 0.90 | 0.49 | 0.87 | 1.00 | 1.00 | 0.84 |
| C576T | 0.61 | 0.82 | 0.62 | 0.80 | 1.00 | 0.99 | 0.81 |
| C576G | 0.66 | 0.86 | 0.62 | 0.83 | 1.00 | 0.99 | 0.84 |
| C576A | 0.60 | 0.86 | 0.63 | 0.82 | 1.00 | 1.00 | 0.82 |
| C577T | 0.56 | 0.87 | 0.47 | 0.73 | 1.00 | 0.99 | 0.72 |
| C577G | 0.45 | 0.51 | 1.00 | 0.90 | 0.50 | 0.56 | 0.89 |
| C577A | 0.54 | 0.85 | 0.68 | 0.82 | 0.97 | 1.00 | 0.81 |
| A578T | 0.64 | 0.78 | 0.70 | 0.81 | 1.00 | 0.99 | 0.82 |
| A578G | 0.62 | 0.75 | 0.86 | 0.92 | 0.89 | 0.89 | 0.95 |
| A578C | 0.67 | 0.61 | 0.80 | 0.98 | 0.76 | 0.77 | 0.98 |
| G579T | 0.67 | 0.83 | 0.66 | 0.84 | 1.00 | 1.00 | 0.84 |
| G579C | 0.64 | 0.79 | 0.78 | 0.84 | 0.99 | 0.98 | 0.86 |
| G579A | 0.62 | 0.71 | 0.61 | 0.76 | 0.97 | 0.97 | 0.76 |
| A580T | 0.61 | 0.81 | 0.65 | 0.80 | 0.99 | 0.99 | 0.82 |
| A580G | 0.62 | 0.85 | 0.49 | 0.77 | 0.99 | 0.99 | 0.76 |
| T581G | 0.67 | 0.95 | 0.68 | 0.88 | 1.00 | 1.00 | 0.88 |
| T581C | 0.55 | 0.76 | 0.79 | 0.81 | 0.97 | 0.99 | 0.82 |
| T581A | 0.60 | 0.77 | 0.80 | 0.83 | 0.97 | 0.97 | 0.85 |
| T581TC | 0.47 | 0.79 | 0.69 | 0.73 | 1.00 | 1.00 | 0.75 |
| T581TA | 0.59 | 0.90 | 0.58 | 0.82 | 1.00 | 1.00 | 0.82 |
| TG582T | 0.74 | 0.83 | 0.88 | 0.97 | 0.91 | 0.94 | 0.95 |
| G582T | 0.65 | 0.79 | 0.65 | 0.81 | 0.99 | 0.97 | 0.84 |
| G582A | 0.52 | 0.83 | 0.44 | 0.59 | 0.91 | 0.91 | 0.58 |
| G583T | 0.43 | 0.78 | 0.76 | 0.72 | 0.99 | 0.98 | 0.77 |
| G583A | 0.55 | 0.72 | 0.84 | 0.85 | 0.92 | 0.93 | 0.87 |
| G584T | 0.72 | 0.73 | 0.73 | 0.87 | 0.93 | 0.83 | 0.99 |
| G584C | 0.43 | 0.70 | 1.00 | 0.92 | 0.79 | 0.81 | 0.95 |
| G584A | 0.58 | 0.71 | 0.99 | 1.00 | 0.80 | 0.84 | 1.00 |
| G585T | 0.47 | 0.80 | 0.77 | 0.77 | 0.99 | 0.99 | 0.81 |
| G585C | 0.54 | 0.78 | 0.64 | 0.75 | 1.00 | 0.99 | 0.78 |
| G585A | 0.62 | 0.74 | 0.79 | 0.82 | 0.97 | 0.97 | 0.84 |
| T586G | 0.58 | 0.75 | 0.76 | 0.79 | 0.98 | 0.94 | 0.85 |
| T586C | 0.60 | 0.77 | 0.77 | 0.84 | 0.96 | 0.98 | 0.83 |
| T586A | 0.57 | 0.74 | 0.92 | 0.93 | 0.87 | 0.88 | 0.95 |
| TC587T | 0.71 | 0.77 | 0.80 | 0.95 | 0.89 | 0.91 | 0.94 |
| C587T | 0.61 | 0.78 | 0.68 | 0.80 | 0.99 | 1.00 | 0.81 |
| C587G | 0.43 | 1.00 | 1.00 | 0.86 | 0.97 | 0.99 | 0.97 |
| C587A | 0.62 | 0.86 | 0.65 | 0.82 | 1.00 | 1.00 | 0.83 |
| C588T | 0.62 | 0.85 | 0.70 | 0.83 | 1.00 | 1.00 | 0.85 |
| C588A | 0.50 | 0.77 | 0.79 | 0.77 | 0.99 | 1.00 | 0.80 |
| C589T | 0.57 | 0.82 | 0.73 | 0.82 | 0.98 | 1.00 | 0.83 |
| C589G | 0.58 | 0.69 | 0.75 | 0.84 | 0.89 | 0.97 | 0.78 |
| C589A | 0.57 | 0.78 | 0.74 | 0.80 | 0.99 | 0.99 | 0.82 |
| A590T | 0.60 | 0.77 | 0.66 | 0.78 | 0.99 | 0.99 | 0.79 |
| A590G | 0.59 | 0.79 | 0.74 | 0.80 | 1.00 | 0.98 | 0.84 |
| A590C | 0.48 | 0.60 | 0.74 | 0.86 | 0.67 | 0.81 | 0.72 |
| C591T | 0.59 | 0.85 | 0.62 | 0.80 | 1.00 | 1.00 | 0.81 |

| | | | | | | | |
|---|---|---|---|---|---|---|---|
| C591G | 0.58 | 0.78 | 0.72 | 0.81 | 0.98 | 1.00 | 0.81 |
| C591A | 0.59 | 0.85 | 0.63 | 0.81 | 1.00 | 1.00 | 0.81 |
| A592T | 0.53 | 0.82 | 0.78 | 0.80 | 1.00 | 1.00 | 0.83 |
| A592G | 0.59 | 0.84 | 0.56 | 0.77 | 1.00 | 1.00 | 0.77 |
| A592C | 0.49 | 0.74 | 0.84 | 0.78 | 0.98 | 1.00 | 0.80 |
| G593T | 0.58 | 0.77 | 0.64 | 0.76 | 1.00 | 0.99 | 0.78 |
| G593A | 0.63 | 0.79 | 0.75 | 0.83 | 0.99 | 0.99 | 0.84 |
| A594T | 0.56 | 0.78 | 0.65 | 0.77 | 0.99 | 0.99 | 0.78 |
| A594G | 0.51 | 0.85 | 0.42 | 0.56 | 0.94 | 0.93 | 0.56 |
| A594C | 0.46 | 0.75 | 0.81 | 0.75 | 0.98 | 0.99 | 0.78 |
| G595T | 0.59 | 0.78 | 0.77 | 0.82 | 0.98 | 0.99 | 0.83 |
| G595C | 0.55 | 0.67 | 0.77 | 0.85 | 0.85 | 0.84 | 0.86 |
| G595A | 0.53 | 0.73 | 0.93 | 0.97 | 0.81 | 0.91 | 0.90 |
| C596T | 0.62 | 0.79 | 0.68 | 0.80 | 1.00 | 0.99 | 0.82 |
| C596A | 0.64 | 0.76 | 0.79 | 0.86 | 0.95 | 0.91 | 0.91 |
| A597T | 0.63 | 0.81 | 0.78 | 0.84 | 0.99 | 0.98 | 0.87 |
| A597G | 0.58 | 0.76 | 0.76 | 0.81 | 0.97 | 0.98 | 0.82 |
| A597C | 0.65 | 0.70 | 0.64 | 0.77 | 0.99 | 0.96 | 0.80 |
| C598T | 0.65 | 0.83 | 0.68 | 0.83 | 1.00 | 1.00 | 0.84 |
| C598G | 0.59 | 0.86 | 0.51 | 0.77 | 0.99 | 1.00 | 0.76 |
| C598A | 0.62 | 0.80 | 0.55 | 0.77 | 1.00 | 0.99 | 0.78 |
| G599A | 0.61 | 0.80 | 0.72 | 0.81 | 1.00 | 0.98 | 0.84 |
| C600T | 0.41 | 0.77 | 0.95 | 0.77 | 0.98 | 0.97 | 0.83 |
| C600A | 0.61 | 0.79 | 0.70 | 0.81 | 0.99 | 1.00 | 0.82 |
| T601C | 0.55 | 0.81 | 0.64 | 0.77 | 0.99 | 1.00 | 0.78 |
| T601A | 0.62 | 0.85 | 0.59 | 0.81 | 1.00 | 1.00 | 0.81 |
| TC602T | 0.77 | 0.79 | 0.79 | 0.96 | 0.89 | 0.90 | 0.96 |
| C602T | 0.63 | 0.82 | 0.82 | 0.84 | 1.00 | 0.96 | 0.90 |
| C602G | 0.69 | 0.78 | 0.68 | 0.83 | 0.98 | 1.00 | 0.82 |
| C603T | 0.52 | 0.81 | 0.58 | 0.73 | 0.98 | 0.99 | 0.73 |
| C603G | 0.61 | 0.79 | 0.77 | 0.82 | 0.99 | 0.99 | 0.84 |
| C603A | 0.57 | 0.79 | 0.66 | 0.78 | 0.99 | 1.00 | 0.79 |
| CT604C | 0.72 | 0.68 | 0.90 | 1.00 | 0.82 | 0.83 | 1.00 |
| T604G | 0.63 | 0.87 | 0.59 | 0.82 | 1.00 | 1.00 | 0.82 |
| T604C | 0.55 | 0.78 | 0.75 | 0.79 | 0.99 | 0.99 | 0.80 |
| T604A | 0.58 | 0.79 | 0.74 | 0.81 | 0.99 | 1.00 | 0.83 |
| T605G | 0.61 | 0.84 | 0.63 | 0.81 | 1.00 | 1.00 | 0.81 |
| T605C | 0.59 | 0.80 | 0.81 | 0.82 | 1.00 | 1.00 | 0.85 |
| T605A | 0.65 | 0.84 | 0.56 | 0.81 | 1.00 | 1.00 | 0.80 |
| TG606T | 0.73 | 0.71 | 0.80 | 0.99 | 0.83 | 0.83 | 0.99 |
| G606T | 0.63 | 0.76 | 0.68 | 0.81 | 0.98 | 0.99 | 0.82 |
| G606A | 0.56 | 0.83 | 0.56 | 0.75 | 1.00 | 1.00 | 0.76 |
| G607T | 0.68 | 0.65 | 0.56 | 0.75 | 0.92 | 0.86 | 0.80 |
| G607C | 0.60 | 0.82 | 0.60 | 0.78 | 0.99 | 0.99 | 0.78 |
| G607A | 0.58 | 0.79 | 0.71 | 0.79 | 1.00 | 0.98 | 0.82 |
| T608C | 0.58 | 0.74 | 0.70 | 0.77 | 0.99 | 0.99 | 0.79 |
| T608A | 0.63 | 0.71 | 0.87 | 0.98 | 0.82 | 0.84 | 0.98 |
| A609T | 0.70 | 0.85 | 0.75 | 0.87 | 0.99 | 1.00 | 0.87 |
| A609G | 0.64 | 0.79 | 0.75 | 0.83 | 0.99 | 0.98 | 0.85 |
| T610G | 0.71 | 0.80 | 0.63 | 0.83 | 0.99 | 0.99 | 0.82 |
| T610C | 0.59 | 0.83 | 0.55 | 0.77 | 1.00 | 1.00 | 0.77 |
| T610A | 0.58 | 0.82 | 0.69 | 0.81 | 0.99 | 1.00 | 0.82 |
| C611T | 0.61 | 0.77 | 0.72 | 0.80 | 1.00 | 0.99 | 0.82 |

| | | | | | | | |
|---|---|---|---|---|---|---|---|
| C611G | 0.65 | 0.87 | 0.63 | 0.83 | 1.00 | 1.00 | 0.83 |
| C611A | 0.58 | 0.81 | 0.64 | 0.79 | 1.00 | 1.00 | 0.80 |
| C612T | 0.59 | 0.85 | 0.60 | 0.79 | 1.00 | 1.00 | 0.80 |
| C612G | 0.57 | 0.83 | 0.54 | 0.76 | 0.99 | 0.99 | 0.76 |
| C612A | 0.62 | 0.84 | 0.61 | 0.80 | 1.00 | 1.00 | 0.81 |
| CT613C | 0.87 | 0.79 | 0.91 | 1.00 | 0.89 | 0.90 | 1.00 |
| T613C | 0.65 | 0.82 | 0.70 | 0.83 | 1.00 | 0.99 | 0.84 |
| T613A | 0.63 | 0.78 | 0.67 | 0.80 | 0.99 | 0.97 | 0.83 |
| G614T | 0.61 | 0.84 | 0.59 | 0.80 | 1.00 | 1.00 | 0.81 |
| G614C | 0.72 | 0.80 | 0.66 | 0.84 | 0.99 | 1.00 | 0.83 |
| G614A | 0.58 | 0.83 | 0.59 | 0.79 | 0.99 | 1.00 | 0.79 |
| C615T | 0.58 | 0.83 | 0.60 | 0.78 | 0.99 | 0.99 | 0.79 |
| C615G | 0.64 | 0.77 | 0.63 | 0.79 | 0.99 | 0.99 | 0.80 |
| C615A | 0.62 | 0.78 | 0.68 | 0.80 | 0.99 | 0.99 | 0.81 |
| T616C | 0.64 | 0.82 | 0.69 | 0.83 | 1.00 | 0.99 | 0.84 |
| G617T | 0.76 | 0.75 | 0.61 | 0.83 | 0.97 | 0.96 | 0.84 |
| G617C | 0.58 | 0.81 | 0.71 | 0.80 | 1.00 | 0.99 | 0.83 |
| G617A | 0.60 | 0.80 | 0.71 | 0.80 | 1.00 | 0.97 | 0.84 |
| GC618G | 0.71 | 0.73 | 0.97 | 1.00 | 0.84 | 0.85 | 1.00 |
| C618T | 0.59 | 0.75 | 0.86 | 0.89 | 0.91 | 0.93 | 0.89 |
| C618G | 0.43 | 0.77 | 1.00 | 0.88 | 0.86 | 0.88 | 0.94 |
| C618A | 0.63 | 0.76 | 1.00 | 1.00 | 0.84 | 0.86 | 1.00 |
| C619T | 0.66 | 0.83 | 0.79 | 0.85 | 0.99 | 0.98 | 0.88 |
| C619G | 0.60 | 0.77 | 0.88 | 0.88 | 0.94 | 0.93 | 0.91 |
| C619A | 0.59 | 0.80 | 0.86 | 0.90 | 0.92 | 1.00 | 0.86 |
| C619CT | 0.74 | 0.78 | 0.96 | 1.00 | 0.87 | 0.89 | 1.00 |
| CT620C | 0.77 | 0.65 | 0.86 | 1.00 | 0.80 | 0.81 | 1.00 |
| T620G | 0.65 | 0.77 | 0.81 | 0.87 | 0.95 | 0.98 | 0.87 |
| T620C | 0.52 | 0.74 | 0.82 | 0.78 | 0.99 | 0.94 | 0.84 |
| T620A | 0.57 | 0.78 | 0.73 | 0.79 | 0.99 | 0.98 | 0.82 |
| T621G | 0.61 | 0.69 | 0.70 | 0.80 | 0.93 | 0.95 | 0.80 |
| T621C | 0.69 | 0.81 | 0.72 | 0.86 | 0.98 | 0.99 | 0.86 |
| T621A | 0.55 | 0.78 | 0.83 | 0.82 | 0.98 | 0.98 | 0.85 |
| T622C | 0.70 | 0.82 | 0.77 | 0.86 | 1.00 | 0.99 | 0.88 |
| T622A | 0.60 | 0.81 | 0.67 | 0.80 | 1.00 | 1.00 | 0.81 |
| G623T | 0.49 | 0.89 | 0.31 | 0.36 | 1.00 | 0.84 | 0.41 |
| G623A | 0.60 | 0.78 | 0.85 | 0.83 | 0.99 | 0.95 | 0.89 |
| C624T | 0.61 | 0.79 | 0.80 | 0.83 | 0.98 | 0.98 | 0.85 |
| C624G | 0.54 | 0.81 | 0.65 | 0.77 | 0.99 | 1.00 | 0.78 |
| C624A | 0.67 | 0.70 | 0.83 | 0.98 | 0.82 | 0.83 | 0.98 |
| T625G | 0.55 | 0.78 | 0.72 | 0.80 | 0.98 | 0.99 | 0.80 |
| T625C | 0.62 | 0.83 | 0.68 | 0.82 | 1.00 | 1.00 | 0.83 |
| T625A | 0.53 | 0.76 | 0.56 | 0.71 | 0.95 | 0.96 | 0.71 |
| G626T | 0.65 | 0.66 | 0.82 | 0.97 | 0.80 | 0.81 | 0.97 |
| G626A | 0.62 | 0.79 | 0.69 | 0.80 | 1.00 | 0.99 | 0.82 |
| C627T | 0.59 | 0.78 | 0.72 | 0.80 | 0.99 | 0.99 | 0.82 |
| C627G | 0.60 | 0.82 | 0.66 | 0.80 | 0.99 | 0.99 | 0.81 |
| C627A | 0.50 | 0.73 | 0.92 | 0.83 | 0.93 | 0.89 | 0.91 |
| T628G | 0.51 | 0.76 | 0.75 | 0.75 | 1.00 | 0.98 | 0.78 |
| T628C | 0.62 | 0.84 | 0.71 | 0.83 | 1.00 | 1.00 | 0.85 |
| T628A | 0.57 | 0.83 | 0.63 | 0.78 | 1.00 | 1.00 | 0.80 |
| TG629T | 0.68 | 0.84 | 0.74 | 0.86 | 0.99 | 1.00 | 0.86 |
| G629T | 0.60 | 0.77 | 0.70 | 0.79 | 1.00 | 0.99 | 0.81 |

| | | | | | | | |
|---|---|---|---|---|---|---|---|
| G629C | 0.60 | 0.78 | 0.67 | 0.80 | 0.99 | 0.99 | 0.81 |
| G629A | 0.57 | 0.78 | 0.65 | 0.77 | 0.99 | 0.99 | 0.79 |
| C630T | 0.58 | 0.77 | 0.71 | 0.82 | 0.95 | 1.00 | 0.80 |
| C630G | 0.58 | 0.83 | 0.77 | 0.82 | 0.99 | 1.00 | 0.84 |
| C630A | 0.55 | 0.79 | 0.75 | 0.79 | 0.99 | 0.99 | 0.82 |
| T631G | 0.49 | 0.74 | 0.95 | 0.86 | 0.91 | 0.89 | 0.92 |
| T631C | 0.68 | 0.81 | 0.72 | 0.84 | 1.00 | 0.99 | 0.85 |
| T631A | 0.54 | 0.79 | 0.78 | 0.79 | 0.99 | 0.99 | 0.82 |
| T632C | 0.62 | 0.80 | 0.75 | 0.82 | 1.00 | 0.99 | 0.84 |
| T632A | 0.65 | 0.83 | 0.63 | 0.82 | 1.00 | 1.00 | 0.82 |
| G633T | 0.56 | 0.85 | 0.61 | 0.79 | 1.00 | 1.00 | 0.80 |
| G633C | 0.62 | 0.75 | 0.92 | 0.98 | 0.84 | 0.87 | 0.97 |
| G633A | 0.62 | 0.79 | 0.65 | 0.79 | 1.00 | 0.99 | 0.80 |
| T634C | 0.54 | 0.74 | 0.75 | 0.78 | 0.98 | 0.98 | 0.79 |
| T634A | 0.53 | 0.76 | 0.79 | 0.78 | 0.99 | 0.98 | 0.82 |
| TG635T | 0.74 | 0.76 | 0.91 | 1.00 | 0.86 | 0.87 | 1.00 |
| G635T | 0.59 | 0.72 | 0.80 | 0.84 | 0.94 | 0.93 | 0.86 |
| G635C | 0.57 | 0.86 | 0.52 | 0.77 | 0.99 | 1.00 | 0.76 |
| G635A | 0.62 | 0.85 | 0.68 | 0.83 | 1.00 | 1.00 | 0.84 |
| G636C | 0.66 | 0.80 | 0.75 | 0.90 | 0.92 | 1.00 | 0.84 |
| G636A | 0.50 | 0.75 | 0.93 | 0.86 | 0.92 | 0.92 | 0.90 |
| C637T | 0.59 | 0.80 | 0.67 | 0.79 | 1.00 | 1.00 | 0.80 |
| C637G | 0.59 | 0.74 | 1.00 | 1.00 | 0.81 | 0.86 | 1.00 |
| C637A | 0.65 | 0.71 | 0.91 | 0.99 | 0.82 | 0.84 | 0.99 |
| T638C | 0.62 | 0.83 | 0.67 | 0.82 | 1.00 | 1.00 | 0.83 |
| T638A | 0.55 | 0.83 | 0.68 | 0.79 | 0.99 | 0.99 | 0.81 |
| G639T | 0.59 | 0.78 | 0.83 | 0.83 | 0.98 | 0.98 | 0.85 |
| G639C | 0.57 | 0.79 | 0.71 | 0.79 | 0.99 | 0.98 | 0.82 |
| G639A | 0.61 | 0.76 | 0.87 | 0.91 | 0.90 | 0.90 | 0.94 |
| C640T | 0.74 | 0.84 | 0.73 | 0.88 | 0.99 | 0.99 | 0.88 |
| C640G | 0.88 | 0.72 | 0.40 | 0.78 | 0.98 | 0.95 | 0.75 |
| C640A | 0.67 | 0.81 | 0.57 | 0.80 | 1.00 | 1.00 | 0.79 |
| A641T | 0.54 | 0.82 | 0.85 | 0.83 | 0.99 | 1.00 | 0.85 |
| A641G | 0.65 | 0.77 | 0.75 | 0.87 | 0.94 | 0.99 | 0.83 |
| A641C | 0.72 | 0.87 | 0.72 | 0.87 | 1.00 | 1.00 | 0.88 |
| C642T | 0.61 | 0.79 | 0.70 | 0.81 | 0.99 | 0.99 | 0.82 |
| C642G | 0.68 | 0.83 | 0.70 | 0.85 | 1.00 | 0.99 | 0.86 |
| C642A | 0.64 | 0.85 | 0.69 | 0.83 | 1.00 | 0.99 | 0.85 |
| T643G | 0.62 | 0.77 | 0.65 | 0.79 | 0.99 | 0.99 | 0.80 |
| T643C | 0.63 | 0.78 | 0.83 | 0.85 | 0.98 | 0.97 | 0.87 |
| T643A | 0.64 | 0.79 | 0.69 | 0.83 | 0.99 | 0.99 | 0.83 |
| G644T | 0.56 | 0.79 | 0.81 | 0.81 | 0.99 | 0.99 | 0.84 |
| G644C | 0.58 | 0.75 | 0.85 | 0.86 | 0.94 | 0.95 | 0.87 |
| G644A | 0.57 | 0.78 | 0.76 | 0.80 | 0.99 | 0.99 | 0.82 |
| G645T | 0.51 | 0.75 | 0.84 | 0.79 | 0.99 | 0.99 | 0.81 |
| G645A | 0.60 | 0.75 | 0.91 | 0.97 | 0.85 | 0.89 | 0.95 |
| C646T | 0.66 | 0.89 | 0.56 | 0.83 | 1.00 | 0.99 | 0.83 |
| C646G | 0.62 | 0.88 | 0.69 | 0.84 | 1.00 | 1.00 | 0.85 |
| C646A | 0.55 | 0.86 | 0.73 | 0.82 | 1.00 | 1.00 | 0.84 |
| G647T | 0.34 | 0.83 | 0.72 | 0.65 | 0.99 | 0.98 | 0.70 |
| G647C | 0.62 | 0.85 | 0.51 | 0.78 | 1.00 | 0.99 | 0.78 |
| G647A | 0.62 | 0.83 | 0.66 | 0.82 | 1.00 | 1.00 | 0.82 |
| A648T | 0.58 | 0.76 | 0.71 | 0.80 | 0.98 | 0.99 | 0.81 |

| | | | | | | | |
|---|---|---|---|---|---|---|---|
| A648G | 0.59 | 0.87 | 0.54 | 0.78 | 1.00 | 1.00 | 0.79 |
| A648C | 0.57 | 0.80 | 0.68 | 0.79 | 1.00 | 1.00 | 0.82 |
| C649T | 0.61 | 0.81 | 0.67 | 0.81 | 1.00 | 0.99 | 0.83 |
| C649G | 0.72 | 0.80 | 0.76 | 0.88 | 0.97 | 0.99 | 0.87 |
| C649A | 0.67 | 0.78 | 0.79 | 0.87 | 0.96 | 0.93 | 0.91 |
| T650G | 0.54 | 0.71 | 0.80 | 0.79 | 0.96 | 0.93 | 0.84 |
| T650C | 0.61 | 0.84 | 0.55 | 0.79 | 1.00 | 1.00 | 0.78 |
| T650A | 0.56 | 0.80 | 0.66 | 0.78 | 1.00 | 0.99 | 0.80 |
| G651T | 0.66 | 0.80 | 0.76 | 0.85 | 0.98 | 1.00 | 0.85 |
| G651C | 0.59 | 0.68 | 0.63 | 0.76 | 0.92 | 0.95 | 0.74 |
| G651A | 0.57 | 0.79 | 0.72 | 0.79 | 0.99 | 0.98 | 0.82 |
| C652T | 0.58 | 0.81 | 0.65 | 0.79 | 1.00 | 1.00 | 0.80 |
| C652G | 0.62 | 0.81 | 0.93 | 0.93 | 0.92 | 0.93 | 0.95 |
| C652A | 0.50 | 0.81 | 0.93 | 0.82 | 0.99 | 0.98 | 0.86 |
| A653T | 0.65 | 0.75 | 0.66 | 0.82 | 0.96 | 1.00 | 0.80 |
| A653G | 0.66 | 0.89 | 0.44 | 0.79 | 1.00 | 1.00 | 0.76 |
| A653C | 0.61 | 0.80 | 0.56 | 0.77 | 0.99 | 0.99 | 0.77 |
| C654T | 0.49 | 0.75 | 0.90 | 0.85 | 0.91 | 0.96 | 0.84 |
| C654G | 0.61 | 0.87 | 0.59 | 0.82 | 1.00 | 1.00 | 0.82 |
| C654A | 0.57 | 0.81 | 0.75 | 0.82 | 0.98 | 1.00 | 0.83 |
| T655G | 0.49 | 0.72 | 0.92 | 0.87 | 0.88 | 0.90 | 0.89 |
| T655C | 0.60 | 0.81 | 0.69 | 0.81 | 0.99 | 1.00 | 0.82 |
| T655A | 0.61 | 0.80 | 0.70 | 0.81 | 1.00 | 0.99 | 0.83 |
| TG656T | 0.57 | 0.64 | 0.99 | 1.00 | 0.74 | 0.78 | 1.00 |
| G656T | 0.60 | 0.79 | 0.72 | 0.81 | 0.99 | 0.98 | 0.83 |
| G656C | 0.63 | 0.84 | 0.76 | 0.84 | 1.00 | 0.99 | 0.87 |
| G656A | 0.61 | 0.80 | 0.64 | 0.80 | 0.99 | 1.00 | 0.80 |
| G657C | 0.60 | 0.79 | 0.82 | 0.83 | 0.98 | 0.96 | 0.87 |
| G657A | 0.59 | 0.77 | 0.79 | 0.82 | 0.98 | 0.98 | 0.84 |
| T658G | 0.56 | 0.91 | 0.57 | 0.79 | 1.00 | 1.00 | 0.80 |
| T658C | 0.61 | 0.80 | 0.70 | 0.81 | 1.00 | 0.99 | 0.82 |
| T658A | 0.49 | 0.75 | 0.81 | 0.77 | 0.99 | 0.99 | 0.79 |
| C659T | 0.65 | 0.83 | 0.61 | 0.82 | 1.00 | 1.00 | 0.82 |
| C659A | 0.70 | 0.78 | 0.57 | 0.80 | 1.00 | 0.99 | 0.80 |
| T660C | 0.62 | 0.81 | 0.61 | 0.79 | 1.00 | 0.99 | 0.80 |
| T660A | 0.57 | 0.78 | 0.67 | 0.78 | 0.99 | 0.99 | 0.79 |
| T661G | 0.81 | 0.80 | 0.60 | 0.86 | 0.98 | 0.99 | 0.83 |
| T661C | 0.65 | 0.81 | 0.65 | 0.82 | 1.00 | 1.00 | 0.82 |
| T661A | 0.62 | 0.79 | 0.74 | 0.82 | 0.99 | 1.00 | 0.83 |
| C662T | 0.63 | 0.87 | 0.45 | 0.76 | 0.99 | 0.99 | 0.74 |
| C662A | 0.63 | 0.90 | 0.51 | 0.82 | 1.00 | 1.00 | 0.80 |
| A663T | 0.60 | 0.83 | 0.64 | 0.80 | 1.00 | 1.00 | 0.81 |
| A663G | 0.61 | 0.80 | 0.76 | 0.82 | 0.99 | 0.99 | 0.84 |
| A663C | 0.59 | 0.78 | 0.54 | 0.73 | 0.99 | 1.00 | 0.73 |
| G664T | 0.64 | 0.82 | 0.70 | 0.83 | 1.00 | 1.00 | 0.84 |
| G664C | 0.62 | 0.81 | 0.58 | 0.79 | 0.99 | 1.00 | 0.79 |
| G664A | 0.54 | 0.70 | 0.81 | 0.82 | 0.91 | 0.91 | 0.85 |
| C665T | 0.61 | 0.82 | 0.63 | 0.80 | 1.00 | 0.99 | 0.81 |
| C665G | 0.64 | 0.87 | 0.69 | 0.84 | 1.00 | 0.99 | 0.86 |
| C665A | 0.61 | 0.78 | 0.66 | 0.80 | 0.98 | 0.99 | 0.80 |
| T666G | 0.71 | 0.83 | 0.41 | 0.78 | 0.98 | 1.00 | 0.73 |
| T666C | 0.68 | 0.81 | 0.57 | 0.80 | 1.00 | 1.00 | 0.80 |
| T666A | 0.57 | 0.77 | 0.77 | 0.80 | 0.99 | 0.98 | 0.83 |

| | | | | | | | |
|---|---|---|---|---|---|---|---|
| A667T | 0.59 | 0.74 | 0.75 | 0.81 | 0.97 | 0.97 | 0.83 |
| A667G | 0.57 | 0.81 | 0.68 | 0.79 | 1.00 | 1.00 | 0.81 |
| C668T | 0.62 | 0.82 | 0.68 | 0.82 | 1.00 | 1.00 | 0.83 |
| C668A | 0.59 | 0.86 | 0.67 | 0.82 | 1.00 | 1.00 | 0.83 |
| T669G | 0.65 | 0.63 | 0.66 | 0.83 | 0.86 | 0.85 | 0.85 |
| T669C | 0.63 | 0.81 | 0.63 | 0.81 | 0.99 | 1.00 | 0.81 |
| T669A | 0.60 | 0.82 | 0.76 | 0.83 | 1.00 | 0.99 | 0.86 |
| TG670T | 0.74 | 0.66 | 0.96 | 1.00 | 0.81 | 0.83 | 1.00 |
| G670T | 0.61 | 0.83 | 0.68 | 0.81 | 1.00 | 0.99 | 0.83 |
| G670C | 0.54 | 0.80 | 0.79 | 0.80 | 0.99 | 1.00 | 0.83 |
| G670A | 0.67 | 0.78 | 0.66 | 0.82 | 1.00 | 1.00 | 0.82 |
| G671T | 0.73 | 0.74 | 0.96 | 1.00 | 0.84 | 0.86 | 1.00 |
| G671A | 0.54 | 0.76 | 0.76 | 0.79 | 0.98 | 0.99 | 0.81 |
| GT672G | 0.81 | 0.94 | 0.98 | 0.94 | 0.99 | 0.98 | 0.98 |
| T672G | 0.71 | 0.86 | 0.62 | 0.87 | 0.99 | 1.00 | 0.85 |
| T672C | 0.64 | 0.75 | 0.75 | 0.83 | 0.97 | 0.98 | 0.83 |
| T672A | 0.57 | 0.79 | 0.73 | 0.80 | 0.99 | 0.98 | 0.83 |
| TG673T | 0.76 | 0.83 | 0.73 | 0.87 | 1.00 | 0.99 | 0.88 |
| G673T | 0.63 | 0.78 | 0.75 | 0.82 | 0.99 | 0.99 | 0.84 |
| G673C | 0.42 | 0.77 | 0.79 | 0.72 | 0.98 | 0.99 | 0.75 |
| G673A | 0.62 | 0.78 | 0.63 | 0.79 | 0.99 | 0.98 | 0.80 |
| G674T | 0.59 | 0.77 | 0.73 | 0.79 | 1.00 | 0.93 | 0.87 |
| G674C | 0.64 | 0.84 | 0.59 | 0.82 | 0.99 | 1.00 | 0.81 |
| G674A | 0.51 | 0.77 | 0.86 | 0.80 | 0.99 | 0.98 | 0.84 |
| C675T | 0.62 | 0.83 | 0.72 | 0.83 | 1.00 | 0.99 | 0.85 |
| C675G | 0.72 | 0.84 | 0.59 | 0.84 | 1.00 | 1.00 | 0.83 |
| C675A | 0.61 | 0.75 | 0.70 | 0.80 | 0.98 | 0.98 | 0.81 |
| G676T | 0.63 | 0.85 | 0.59 | 0.82 | 1.00 | 1.00 | 0.81 |
| G676C | 0.48 | 0.76 | 0.82 | 0.77 | 0.99 | 0.99 | 0.80 |
| G676A | 0.66 | 0.83 | 0.62 | 0.82 | 1.00 | 1.00 | 0.82 |
| G677T | 0.74 | 0.85 | 0.61 | 0.86 | 1.00 | 1.00 | 0.85 |
| G677A | 0.58 | 0.62 | 0.65 | 0.73 | 0.86 | 0.74 | 0.86 |
| A678T | 0.65 | 0.80 | 0.66 | 0.81 | 0.99 | 0.99 | 0.81 |
| A678G | 0.62 | 0.83 | 0.59 | 0.80 | 0.99 | 1.00 | 0.79 |
| AG679A | 0.79 | 0.82 | 0.93 | 1.00 | 0.89 | 0.90 | 1.00 |
| G679T | 0.59 | 0.76 | 0.75 | 0.80 | 0.99 | 0.99 | 0.82 |
| G679A | 0.58 | 0.80 | 0.81 | 0.82 | 0.99 | 1.00 | 0.85 |
| G680T | 0.55 | 0.73 | 0.71 | 0.78 | 0.96 | 1.00 | 0.76 |
| G680A | 0.66 | 0.80 | 0.72 | 0.84 | 0.99 | 1.00 | 0.84 |
| GA681G | 0.80 | 0.87 | 0.83 | 0.93 | 0.97 | 1.00 | 0.91 |
| A681T | 0.66 | 0.80 | 0.57 | 0.79 | 1.00 | 1.00 | 0.78 |
| A681G | 0.64 | 0.87 | 0.72 | 0.85 | 1.00 | 1.00 | 0.86 |
| A682T | 0.60 | 0.81 | 0.71 | 0.81 | 0.99 | 0.99 | 0.83 |
| A682G | 0.67 | 0.84 | 0.63 | 0.83 | 1.00 | 0.99 | 0.84 |
| G683T | 0.64 | 0.76 | 0.82 | 0.87 | 0.94 | 0.90 | 0.94 |
| G683C | 0.64 | 0.69 | 0.77 | 0.91 | 0.86 | 0.86 | 0.92 |
| G683A | 0.49 | 0.75 | 0.80 | 0.76 | 0.98 | 0.99 | 0.79 |
| C684T | 0.65 | 0.80 | 0.70 | 0.82 | 0.99 | 0.99 | 0.83 |
| C684G | 0.66 | 0.91 | 0.53 | 0.85 | 1.00 | 1.00 | 0.83 |
| C684A | 0.56 | 0.79 | 0.75 | 0.81 | 0.98 | 1.00 | 0.82 |
| A685T | 0.61 | 0.79 | 0.71 | 0.81 | 1.00 | 0.99 | 0.82 |
| A685G | 0.61 | 0.79 | 0.71 | 0.81 | 0.99 | 0.98 | 0.83 |
| A685C | 0.49 | 0.78 | 0.60 | 0.71 | 0.98 | 0.99 | 0.72 |

| | | | | | | | |
|---|---|---|---|---|---|---|---|
| G686T | 0.55 | 0.79 | 0.76 | 0.81 | 0.98 | 1.00 | 0.82 |
| G686C | 0.72 | 0.78 | 0.70 | 0.85 | 0.98 | 0.98 | 0.85 |
| G686A | 0.56 | 0.75 | 0.75 | 0.78 | 1.00 | 0.99 | 0.80 |
| C687T | 0.65 | 0.77 | 0.62 | 0.82 | 0.97 | 0.99 | 0.80 |
| C687G | 0.61 | 0.78 | 0.57 | 0.76 | 0.99 | 0.98 | 0.77 |
| C687A | 0.62 | 0.85 | 0.68 | 0.83 | 1.00 | 1.00 | 0.85 |
| T688G | 0.61 | 0.78 | 0.69 | 0.80 | 1.00 | 0.99 | 0.81 |
| T688C | 0.55 | 0.76 | 0.68 | 0.78 | 0.97 | 1.00 | 0.77 |
| T688A | 0.60 | 0.79 | 0.63 | 0.80 | 0.97 | 1.00 | 0.78 |
| A689T | 0.66 | 0.78 | 0.70 | 0.82 | 1.00 | 0.99 | 0.83 |
| A689G | 0.63 | 0.78 | 0.74 | 0.83 | 0.99 | 0.99 | 0.84 |
| A689C | 0.66 | 0.80 | 0.76 | 0.83 | 1.00 | 1.00 | 0.85 |
| G690T | 0.79 | 0.89 | 0.57 | 0.90 | 0.97 | 1.00 | 0.85 |
| G690C | 0.78 | 0.91 | 0.69 | 0.90 | 1.00 | 1.00 | 0.90 |
| G690A | 0.62 | 0.91 | 0.33 | 0.64 | 0.96 | 0.93 | 0.61 |
| G691T | 0.93 | 0.79 | 0.07 | 0.64 | 0.51 | 0.50 | 0.28 |
| G691A | 0.60 | 0.89 | 0.07 | 0.37 | 0.58 | 0.29 | 0.27 |
| T692G | 0.79 | 0.86 | 0.04 | 0.53 | 0.55 | 0.48 | 0.16 |
| T692C | 0.66 | 0.81 | 0.11 | 0.36 | 0.61 | 0.22 | 0.49 |
| T692A | 0.69 | 0.84 | 0.05 | 0.27 | 0.83 | 0.10 | 0.67 |
| G693T | 0.43 | 0.84 | 0.16 | 0.28 | 0.53 | 0.17 | 0.47 |
| G693C | 0.86 | 0.90 | 0.25 | 0.76 | 1.00 | 0.99 | 0.68 |
| G693A | 0.50 | 0.75 | 0.79 | 0.78 | 0.98 | 0.99 | 0.80 |
| A694T | 0.60 | 0.90 | 0.14 | 0.52 | 0.46 | 0.52 | 0.27 |
| A694G | 0.57 | 0.90 | 0.28 | 0.43 | 0.92 | 0.94 | 0.37 |
| A694C | 0.59 | 0.96 | 0.32 | 0.69 | 0.99 | 0.98 | 0.65 |
| G695T | 0.54 | 0.90 | 0.29 | 0.61 | 0.54 | 0.84 | 0.34 |
| G695A | 0.66 | 0.88 | 0.12 | 0.46 | 0.58 | 0.44 | 0.34 |
| T696C | 0.72 | 0.88 | 0.44 | 0.81 | 1.00 | 1.00 | 0.78 |
| T696A | 0.74 | 0.87 | 0.64 | 0.87 | 1.00 | 0.97 | 0.89 |
| T697G | 0.91 | 0.79 | 0.78 | 1.00 | 0.89 | 0.87 | 1.00 |
| T697C | 0.66 | 0.83 | 0.63 | 0.83 | 1.00 | 0.99 | 0.83 |
| T697A | 0.49 | 0.72 | 0.78 | 0.75 | 0.97 | 0.96 | 0.79 |
| C698T | 0.64 | 0.80 | 0.71 | 0.83 | 0.98 | 0.99 | 0.84 |
| C698A | 0.53 | 0.74 | 0.71 | 0.75 | 0.98 | 0.99 | 0.76 |
| T699C | 0.55 | 0.77 | 0.77 | 0.78 | 1.00 | 0.98 | 0.82 |
| T699A | 0.57 | 0.81 | 0.69 | 0.79 | 1.00 | 0.99 | 0.81 |
| C700T | 0.53 | 0.79 | 0.74 | 0.78 | 1.00 | 1.00 | 0.80 |
| C700G | 0.66 | 0.79 | 0.68 | 0.87 | 0.94 | 1.00 | 0.83 |
| C700A | 0.60 | 0.78 | 0.59 | 0.77 | 0.99 | 1.00 | 0.77 |
| T701G | 0.37 | 0.91 | 0.96 | 0.80 | 0.98 | 1.00 | 0.87 |
| T701C | 0.61 | 0.78 | 0.66 | 0.79 | 0.99 | 0.99 | 0.80 |
| T701A | 0.58 | 0.81 | 0.73 | 0.81 | 1.00 | 1.00 | 0.83 |
| TG702T | 0.80 | 0.72 | 0.82 | 1.00 | 0.84 | 0.84 | 1.00 |
| G702T | 0.64 | 0.79 | 0.87 | 0.90 | 0.93 | 0.92 | 0.94 |
| G702C | 0.55 | 0.83 | 0.68 | 0.79 | 0.99 | 1.00 | 0.81 |
| G702A | 0.60 | 0.81 | 0.73 | 0.82 | 1.00 | 1.00 | 0.83 |
| C703T | 0.53 | 0.78 | 0.80 | 0.80 | 0.99 | 0.99 | 0.83 |
| C703G | 0.39 | 0.65 | 0.96 | 0.80 | 0.83 | 0.81 | 0.89 |
| C703A | 0.61 | 0.78 | 0.61 | 0.78 | 0.99 | 1.00 | 0.78 |
| CT704C | 0.69 | 0.77 | 0.77 | 0.87 | 0.96 | 0.93 | 0.91 |
| T704G | 0.72 | 0.80 | 0.84 | 0.96 | 0.90 | 0.92 | 0.96 |
| T704C | 0.54 | 0.77 | 0.76 | 0.78 | 1.00 | 0.96 | 0.84 |

| | | | | | | | |
|---|---|---|---|---|---|---|---|
| T704A | 0.64 | 0.77 | 0.70 | 0.81 | 1.00 | 1.00 | 0.82 |
| T704TC | 0.35 | 0.78 | 1.00 | 0.79 | 0.90 | 0.89 | 0.91 |
