## Supplementary material for "Exon definition facilitates reliable control of alternative splicing in the *RON* proto-oncogene": Table S2

The following tables were obtained with the Human Splicing Finder, Version 3.1

http://www.umd.be/HSF/ and FO Desmet, Hamroun D, Lalande M, Collod-Beroud G, Claustres M, Beroud C. Human Splicing Finder: an online bioinformatics tool to predict splicing signals. Nucleic Acid Research, 2009

### ESE Finder matrices for SRp40, SC35, SF2/ASF and SRp55 proteins

Cartegni L., Wang J., Zhu Z., Zhang M. Q., Krainer A. R.; 2003. ESEfinder: a web resource to identify exonic splicing enhancers, Nucleic Acid Research, 2003, Vol. 31, No.13 3568-3571

| Sequence Position | cDNA Position | Linked SR protein | Enhancer motif |
|---|---|---|---|
| +3 | SRp40 | TGAGAGG | 84.79 |
| +11 | SC35 | AGCTTCCA | 76.46 |
| +12 | SC35 | GCTTCCAG | 76.34 |
| +16 | SRp40 | CCAGAGC | 82.46 |
| +17 | SF2/ASF (IgM-BRCA1) | CAGAGCA | 80.54 |
| +17 | SF2/ASF | CAGAGCA | 80.02 |
| +20 | SRp55 | AGCAGC | 74.95 |
| +23 | SRp55 | AGCAGC | 74.95 |
| +25 | SF2/ASF (IgM-BRCA1) | CAGCTGT | 76.00 |
| +25 | SF2/ASF | CAGCTGT | 76.06 |
| +31 | SRp55 | TGCCGC | 76.16 |
| +33 | SF2/ASF (IgM-BRCA1) | CCGCCTT | 72.77 |
| +42 | SRp55 | TGAATA | 76.10 |
| +51 | SC35 | GGTCCGAG | 84.63 |
| +58 | SC35 | GACCCCCA | 91.09 |
| +60 | SF2/ASF (IgM-BRCA1) | CCCCCAG | 72.38 |
| +61 | SRp40 | CCCCAGG | 83.47 |
| +61 | SF2/ASF (IgM-BRCA1) | CCCCAGG | 71.38 |
| +62 | SF2/ASF (IgM-BRCA1) | CCCAGGG | 87.62 |
| +62 | SF2/ASF | CCCAGGG | 82.18 |
| +74 | SRp40 | TGGCAGG | 79.46 |
| +75 | SF2/ASF (IgM-BRCA1) | GGCAGGG | 72.77 |
| +75 | SF2/ASF | GGCAGGG | 80.37 |
| +85 | SF2/ASF (IgM-BRCA1) | CTGAGTG | 74.69 |
| +86 | SRp40 | TGAGTGC | 78.20 |
| +90 | SC35 | TGCCCGAG | 75.54 |
| +90 | SF2/ASF (IgM-BRCA1) | TGCCCGA | 73.00 |
| +90 | SF2/ASF | TGCCCGA | 74.43 |
| +93 | SF2/ASF (IgM-BRCA1) | CCGAGGG | 84.69 |
| +93 | SF2/ASF | CCGAGGG | 80.72 |
| +105 | SC35 | AGCTGCTG | 82.36 |
| +107 | SRp40 | CTGCTGG | 78.86 |
| +107 | SF2/ASF (IgM-BRCA1) | CTGCTGG | 71.15 |

| | | | |
|---|---|---|---|
| +116 | SRp40 | TTACACT | 82.04 |
| +118 | SRp40 | ACACTGC | 87.13 |
| +121 | SF2/ASF (IgM-BRCA1) | CTGCCTG | 73.15 |
| +122 | SRp40 | TGCCTGG | 79.22 |
| +127 | SC35 | GGCTTTCG | 79.41 |
| +133 | SRp55 | CGCTTC | 76.04 |
| +134 | SC35 | GCTTCCTA | 79.84 |
| +139 | SRp40 | CTACCCC | 78.62 |
| +140 | SC35 | TACCCCCA | 78.49 |
| +145 | SRp40 | CCACCCC | 80.48 |
| +146 | SF2/ASF (IgM-BRCA1) | CACCCCA | 81.92 |
| +146 | SF2/ASF | CACCCCA | 79.27 |
| +155 | SF2/ASF (IgM-BRCA1) | CACCCAG | 77.62 |
| +155 | SF2/ASF | CACCCAG | 75.01 |
| +157 | SF2/ASF (IgM-BRCA1) | CCCAGTG | 75.69 |
| +158 | SRp40 | CCAGTGC | 80.96 |
| +173 | SC35 | TTCCACTG | 80.52 |
| +175 | SRp40 | CCACTGA | 82.10 |
| +184 | SRp40 | CCTGAGG | 81.26 |
| +185 | SF2/ASF (IgM-BRCA1) | CTGAGGA | 89.92 |
| +185 | SF2/ASF | CTGAGGA | 88.47 |
| +186 | SRp55 | TGAGGA | 74.82 |
| +205 | SRp40 | TTTGAGG | 81.56 |
| +216 | SRp40 | TGTAAGG | 79.40 |
| +229 | SF2/ASF (IgM-BRCA1) | GGCAGGG | 72.77 |
| +229 | SF2/ASF | GGCAGGG | 80.37 |
| +262 | SF2/ASF (IgM-BRCA1) | CAGCCTA | 79.77 |
| +262 | SF2/ASF | CAGCCTA | 80.43 |
| +264 | SC35 | GCCTACTG | 83.22 |
| +266 | SRp40 | CTACTGG | 94.49 |
| +275 | SC35 | GGTCCTCA | 79.23 |
| +278 | SRp40 | CCTCATG | 79.46 |
| +279 | SF2/ASF (IgM-BRCA1) | CTCATGA | 80.54 |
| +279 | SF2/ASF | CTCATGA | 78.80 |
| +289 | SRp40 | TCTCTGC | 87.13 |
| +292 | SRp40 | CTGCAGG | 80.36 |
| +293 | SRp55 | TGCAGG | 75.65 |
| +293 | SF2/ASF (IgM-BRCA1) | TGCAGGA | 74.54 |
| +293 | SF2/ASF | TGCAGGA | 76.65 |
| +297 | SC35 | GGATATTG | 78.92 |
| +309 | SC35 | GGGCGCTG | 80.33 |
| +315 | SRp55 | TGTGGC | 80.97 |
| +320 | SF2/ASF (IgM-BRCA1) | CTGACTG | 76.31 |

| | | | |
|---|---|---|---|
| +320 | SF2/ASF | CTGACTG | 75.42 |
| +321 | SRp40 | TGACTGT | 79.34 |
| +325 | SRp55 | TGTGTG | 74.82 |
| +329 | SRp55 | TGGGTA | 74.12 |
| +341 | SC35 | TGACCGTG | 75.60 |
| +344 | SF2/ASF (IgM-BRCA1) | CCGTGGG | 71.92 |
| +354 | SRp40 | TGAGAGC | 79.70 |
| +358 | SC35 | AGCTGCCA | 75.66 |
| +360 | SF2/ASF (IgM-BRCA1) | CTGCCAG | 71.38 |
| +365 | SF2/ASF (IgM-BRCA1) | AGCACGA | 75.92 |
| +365 | SF2/ASF | AGCACGA | 82.76 |
| +375 | SF2/ASF (IgM-BRCA1) | CCGGGGG | 71.46 |
| +376 | SF2/ASF (IgM-BRCA1) | CGGGGGG | 78.92 |
| +390 | SRp40 | TGTCTGC | 82.22 |
| +395 | SC35 | GCCCCCTG | 87.52 |
| +396 | SF2/ASF (IgM-BRCA1) | CCCCCTG | 74.15 |
| +401 | SC35 | TGCCCCCA | 80.70 |
| +403 | SF2/ASF (IgM-BRCA1) | CCCCCAT | 73.92 |
| +407 | SC35 | CATCCCTG | 81.81 |
| +409 | SRp40 | TCCCTGC | 79.04 |
| +413 | SRp55 | TGCAGC | 86.98 |
| +422 | SF2/ASF | GCCAGGA | 77.23 |
| +426 | SC35 | GGATGGTG | 81.32 |
| +431 | SC35 | GTGCCCCA | 75.11 |
| +439 | SRp40 | TTGCAGG | 82.51 |
| +440 | SRp55 | TGCAGG | 75.65 |
| +440 | SF2/ASF (IgM-BRCA1) | TGCAGGT | 72.77 |
| +440 | SF2/ASF | TGCAGGT | 74.61 |
| +450 | SF2/ASF (IgM-BRCA1) | CAGCCCA | 79.00 |
| +450 | SF2/ASF | CAGCCCA | 77.81 |
| +453 | SF2/ASF (IgM-BRCA1) | CCCAGCT | 76.46 |
| +455 | SF2/ASF (IgM-BRCA1) | CAGCTGG | 74.46 |
| +455 | SF2/ASF | CAGCTGG | 73.85 |
| +461 | SC35 | GACCTCCC | 75.29 |
| +464 | SF2/ASF (IgM-BRCA1) | CTCCCTG | 76.08 |
| +465 | SRp40 | TCCCTGG | 84.13 |
| +466 | SF2/ASF (IgM-BRCA1) | CCCTGGG | 74.85 |
| +473 | SRp40 | AAACACG | 78.32 |
| +474 | SF2/ASF | AACACGG | 81.13 |
| +475 | SRp40 | ACACGGG | 78.44 |
| +476 | SF2/ASF (IgM-BRCA1) | CACGGGC | 72.00 |
| +482 | SF2/ASF (IgM-BRCA1) | CAGAGGG | 89.92 |
| +482 | SF2/ASF | CAGAGGG | 90.74 |

| | | | |
|---|---|---|---|
| +487 | SC35 | GGCCTACA | 80.82 |
| +488 | SC35 | GCCTACAG | 76.95 |
| +490 | SRp40 | CTACAGG | 95.99 |
| +500 | SC35 | GGCCTGAG | 83.47 |
| +503 | SF2/ASF (IgM-BRCA1) | CTGAGTT | 76.23 |
| +503 | SF2/ASF | CTGAGTT | 74.08 |
| +516 | SC35 | TGCCCCCA | 80.70 |
| +517 | SC35 | GCCCCCAG | 81.25 |
| +518 | SF2/ASF (IgM-BRCA1) | CCCCCAG | 72.38 |
| +519 | SRp40 | CCCCAGG | 83.47 |
| +519 | SF2/ASF (IgM-BRCA1) | CCCCAGG | 71.38 |
| +520 | SF2/ASF (IgM-BRCA1) | CCCAGGT | 89.15 |
| +520 | SF2/ASF | CCCAGGT | 84.39 |
| +524 | SC35 | GGTCTGCG | 82.30 |
| +527 | SF2/ASF (IgM-BRCA1) | CTGCGTA | 74.85 |
| +528 | SRp55 | TGCGTA | 93.71 |
| +547 | SC35 | ATATCCTG | 77.63 |
| +553 | SRp55 | TGGGTA | 74.12 |
| +565 | SRp55 | TGCGGC | 90.89 |
| +567 | SF2/ASF (IgM-BRCA1) | CGGCCAG | 76.92 |
| +568 | SC35 | GGCCAGGG | 77.26 |
| +577 | SF2/ASF (IgM-BRCA1) | CAGATGG | 77.62 |
| +577 | SF2/ASF | CAGATGG | 79.62 |
| +583 | SC35 | GGGTCCCA | 77.69 |
| +584 | SC35 | GGTCCCAC | 76.34 |
| +585 | SC35 | GTCCCACA | 81.50 |
| +587 | SF2/ASF (IgM-BRCA1) | CCCACAG | 75.54 |
| +588 | SRp40 | CCACAGA | 83.59 |
| +589 | SRp55 | CACAGA | 75.40 |
| +589 | SF2/ASF (IgM-BRCA1) | CACAGAG | 79.15 |
| +589 | SF2/ASF | CACAGAG | 77.23 |
| +590 | SRp40 | ACAGAGC | 78.32 |
| +591 | SF2/ASF (IgM-BRCA1) | CAGAGCA | 80.54 |
| +591 | SF2/ASF | CAGAGCA | 80.02 |
| +598 | SF2/ASF (IgM-BRCA1) | CGCTCCT | 72.77 |
| +607 | SC35 | GTATCCTG | 88.38 |
| +613 | SRp55 | TGCTGC | 76.16 |
| +615 | SF2/ASF (IgM-BRCA1) | CTGCCTT | 74.69 |
| +622 | SRp55 | TGCTGC | 76.16 |
| +622 | SC35 | TGCTGCTG | 80.52 |
| +625 | SRp55 | TGCTGC | 76.16 |
| +628 | SC35 | TGCTTGTG | 75.72 |
| +632 | SRp55 | TGTGGC | 80.97 |

| | | | |
|---|---|---|---|
| +639 | SRp40 | GCACTGG | 83.53 |
| +640 | SF2/ASF (IgM-BRCA1) | CACTGGC | 72.46 |
| +642 | SF2/ASF (IgM-BRCA1) | CTGGCGA | 78.31 |
| +642 | SF2/ASF | CTGGCGA | 75.13 |
| +644 | SC35 | GGCGACTG | 80.58 |
| +646 | SRp40 | CGACTGC | 86.35 |
| +651 | SRp40 | GCACTGG | 83.53 |
| +652 | SF2/ASF (IgM-BRCA1) | CACTGGT | 81.62 |
| +652 | SF2/ASF | CACTGGT | 80.14 |
| +657 | SC35 | GTCTTCAG | 83.65 |
| +659 | SRp40 | CTTCAGC | 84.61 |
| +663 | SC35 | AGCTACTG | 84.94 |
| +665 | SRp40 | CTACTGG | 94.49 |
| +675 | SF2/ASF (IgM-BRCA1) | CGGAGGA | 95.46 |
| +675 | SF2/ASF | CGGAGGA | 92.37 |
| +682 | SRp55 | AGCAGC | 74.95 |
| +695 | SC35 | GTTCTCTG | 87.83 |
| +697 | SRp40 | TCTCTGC | 87.13 |

### RESCUE ESE hexamers

Fairbrother WG, Yeh RF, Sharp PA, Burge CB. Predictive identification of exonic splicing enhancers in human genes. Science. 2002 Aug 9;297(5583):1007-13

| Sequence Position | cDNA Position | Enhancer motif |
|---|---|---|
| 40 | +40 | CCTGAA |
| 100 | +100 | GATGGA |
| 177 | +177 | ACTGAA |
| 178 | +178 | CTGAAG |
| 179 | +179 | TGAAGC |
| 187 | +187 | GAGGAG |
| 248 | +248 | CTGAAA |
| 249 | +249 | TGAAAG |
| 321 | +321 | TGACTG |
| 335 | +335 | TCAACG |
| 471 | +471 | GGAAAC |
| 472 | +472 | GAAACA |
| 482 | +482 | CAGAGG |
| 658 | +658 | TCTTCA |
| 659 | +659 | CTTCAG |
| 676 | +676 | GGAGGA |
| 677 | +677 | GAGGAA |
| 678 | +678 | AGGAAG |

| | | |
|---|---|---|
| 680 | +680 | GAAGCA |
| 681 | +681 | AAGCAG |

### Predicted PESE Octamers from Zhang & Chasin.

Zhang XH, Chasin LA. Computational definition of sequence motifs governing constitutive exon splicing. Genes Dev. 2004 Jun 1;18(11):1241-50

| Sequence Position | cDNA Position | Enhancer motif | Motif value (0-100) |
|---|---|---|---|
| 9 | +9 | GCAGCTTC | 30.44 |
| 21 | +21 | GCAGCAGC | 10.87 |
| 24 | +24 | GCAGCTGT | 39.34 |
| 25 | +25 | CAGCTGTG | 38.37 |
| 57 | +57 | AGACCCCC | 38.13 |
| 81 | +81 | GAATCTGA | 30.83 |
| 101 | +101 | ATGGAGCT | 30.38 |
| 156 | +156 | ACCCAGTG | 26.53 |
| 176 | +176 | CACTGAAG | 38.12 |
| 177 | +177 | ACTGAAGC | 37.74 |
| 184 | +184 | CCTGAGGA | 44.48 |
| 185 | +185 | CTGAGGAG | 48.52 |
| 186 | +186 | TGAGGAGC | 40.81 |
| 187 | +187 | GAGGAGCA | 57.89 |
| 188 | +188 | AGGAGCAT | 33.2 |
| 245 | +245 | GATCTGAA | 36.71 |
| 268 | +268 | ACTGGCTG | 31.31 |
| 338 | +338 | ACGTGACC | 31.54 |
| 360 | +360 | CTGCCAGC | 47.59 |
| 454 | +454 | CCAGCTGG | 42.69 |
| 455 | +455 | CAGCTGGA | 49.24 |
| 457 | +457 | GCTGGACC | 40.48 |
| 505 | +505 | GAGTTGCC | 30.64 |
| 508 | +508 | TTGCCACC | 24.19 |
| 511 | +511 | CCACCTGC | 26.78 |
| 513 | +513 | ACCTGCCC | 55 |
| 609 | +609 | ATCCTGCT | 23.21 |
| 610 | +610 | TCCTGCTG | 44.28 |
| 618 | +618 | CCTTTGCT | 32.21 |
| 622 | +622 | TGCTGCTG | 34.72 |
| 657 | +657 | GTCTTCAG | 34.58 |
| 673 | +673 | GGCGGAGG | 1.21 |
| 676 | +676 | GGAGGAAG | 47.57 |
| 677 | +677 | GAGGAAGC | 42.39 |

Ratio PESE/PESS: 6.8

### Silencer motifs from Sironi et al.

Sironi M, Menozzi G, Riva L, Cagliani R, Comi GP, Bresolin N, Giorda R, Pozzoli U. Silencer elements as possible inhibitors of pseudoexon splicing. Nucleic Acids Res. 2004 Mar 19;32(5):1783-91

| Sequence Position | cDNA Position | Sironi's motif | Silencer motif |
|---|---|---|---|
| +1 | Motif 2 - [T/G]G[T/A]GGGG | TGTGAGAG | 72.42 |
| +3 | Motif 1 - CTAGAGGT | TGAGAGGC | 80.93 |
| +3 | Motif 2 - [T/G]G[T/A]GGGG | TGAGAGGC | 80.20 |
| +5 | Motif 2 - [T/G]G[T/A]GGGG | AGAGGCAG | 62.54 |
| +12 | Motif 3 - TCTCCCAA | GCTTCCAG | 76.97 |
| +16 | Motif 1 - CTAGAGGT | CCAGAGCA | 79.23 |
| +29 | Motif 2 - [T/G]G[T/A]GGGG | TGTGCCGC | 66.35 |
| +36 | Motif 3 - TCTCCCAA | CCTTCCTG | 69.14 |
| +46 | Motif 2 - [T/G]G[T/A]GGGG | TATGTGGT | 66.35 |
| +48 | Motif 2 - [T/G]G[T/A]GGGG | TGTGGTCC | 73.23 |
| +51 | Motif 3 - TCTCCCAA | GGTCCGAG | 62.64 |
| +55 | Motif 1 - CTAGAGGT | CGAGACCC | 64.72 |
| +59 | Motif 3 - TCTCCCAA | ACCCCCAG | 77.56 |
| +63 | Motif 1 - CTAGAGGT | CCAGGGAT | 68.31 |
| +67 | Motif 2 - [T/G]G[T/A]GGGG | GGATGGGT | 86.07 |
| +68 | Motif 2 - [T/G]G[T/A]GGGG | GATGGGTG | 68.12 |
| +70 | Motif 2 - [T/G]G[T/A]GGGG | TGGGTGGC | 70.71 |
| +71 | Motif 2 - [T/G]G[T/A]GGGG | GGGTGGCA | 66.56 |
| +72 | Motif 2 - [T/G]G[T/A]GGGG | GGTGGCAG | 69.35 |
| +75 | Motif 2 - [T/G]G[T/A]GGGG | GGCAGGGA | 69.39 |
| +76 | Motif 2 - [T/G]G[T/A]GGGG | GCAGGGAA | 66.30 |
| +86 | Motif 2 - | TGAGTGCC | 70.18 |

| | | | |
|---|---|---|---|
| | [T/G]G[T/A]GGGG | | |
| +92 | Motif 1 - CTAGAGGT | CCCGAGGG | 72.64 |
| +94 | Motif 1 - CTAGAGGT | CGAGGGGA | 73.01 |
| +94 | Motif 2 - [T/G]G[T/A]GGGG | CGAGGGGA | 87.24 |
| +99 | Motif 2 - [T/G]G[T/A]GGGG | GGATGGAG | 75.24 |
| +100 | Motif 2 - [T/G]G[T/A]GGGG | GATGGAGC | 63.28 |
| +101 | Motif 1 - CTAGAGGT | ATGGAGCT | 68.62 |
| +103 | Motif 2 - [T/G]G[T/A]GGGG | GGAGCTGC | 60.23 |
| +108 | Motif 2 - [T/G]G[T/A]GGGG | TGCTGGCT | 69.63 |
| +120 | Motif 3 - TCTCCCAA | ACTGCCTG | 66.29 |
| +134 | Motif 3 - TCTCCCAA | GCTTCCTA | 71.27 |
| +135 | Motif 3 - TCTCCCAA | CTTCCTAC | 66.41 |
| +138 | Motif 3 - TCTCCCAA | CCTACCCC | 67.86 |
| +140 | Motif 3 - TCTCCCAA | TACCCCCA | 65.38 |
| +141 | Motif 3 - TCTCCCAA | ACCCCCAC | 77.98 |
| +145 | Motif 3 - TCTCCCAA | CCACCCCA | 73.01 |
| +146 | Motif 3 - TCTCCCAA | CACCCCAT | 63.78 |
| +150 | Motif 3 - TCTCCCAA | CCATCCAC | 63.64 |
| +153 | Motif 3 - TCTCCCAA | TCCACCCA | 61.63 |
| +154 | Motif 3 - TCTCCCAA | CCACCCAG | 80.41 |
| +158 | Motif 1 - CTAGAGGT | CCAGTGCC | 65.60 |
| +160 | Motif 3 - TCTCCCAA | AGTGCCAA | 65.08 |
| +171 | Motif 3 - TCTCCCAA | AGTTCCAC | 61.79 |
| +184 | Motif 1 - CTAGAGGT | CCTGAGGA | 72.58 |
| +186 | Motif 2 - [T/G]G[T/A]GGGG | TGAGGAGC | 80.20 |
| +193 | Motif 3 - TCTCCCAA | CATGCCAT | 61.93 |
| +205 | Motif 1 - CTAGAGGT | TTTGAGGT | 74.76 |
| +205 | Motif 2 - [T/G]G[T/A]GGGG | TTTGAGGT | 66.35 |
| +207 | Motif 2 - [T/G]G[T/A]GGGG | TGAGGTAA | 69.37 |
| +216 | Motif 2 - [T/G]G[T/A]GGGG | TGTAAGGG | 68.11 |
| +217 | Motif 2 - [T/G]G[T/A]GGGG | GTAAGGGA | 61.98 |
| +218 | Motif 1 - CTAGAGGT | TAAGGGAT | 60.88 |
| +218 | Motif 2 - [T/G]G[T/A]GGGG | TAAGGGAT | 69.37 |
| +222 | Motif 1 - CTAGAGGT | GGATAGGG | 62.30 |

| | | | |
|---|---|---|---|
| +222 | Motif 2 - [T/G]G[T/A]GGGG | GGATAGGG | 69.17 |
| +223 | Motif 2 - [T/G]G[T/A]GGGG | GATAGGGG | 65.03 |
| +224 | Motif 1 - CTAGAGGT | ATAGGGGC | 71.07 |
| +224 | Motif 2 - [T/G]G[T/A]GGGG | ATAGGGGC | 73.38 |
| +225 | Motif 2 - [T/G]G[T/A]GGGG | TAGGGGCA | 60.69 |
| +229 | Motif 2 - [T/G]G[T/A]GGGG | GGCAGGGA | 69.39 |
| +230 | Motif 2 - [T/G]G[T/A]GGGG | GCAGGGAC | 66.30 |
| +238 | Motif 2 - [T/G]G[T/A]GGGG | AGTTGGGG | 85.37 |
| +239 | Motif 2 - [T/G]G[T/A]GGGG | GTTGGGGA | 80.18 |
| +253 | Motif 2 - [T/G]G[T/A]GGGG | AGTAGGGG | 78.18 |
| +254 | Motif 1 - CTAGAGGT | GTAGGGGC | 68.44 |
| +254 | Motif 2 - [T/G]G[T/A]GGGG | GTAGGGGC | 77.13 |
| +255 | Motif 2 - [T/G]G[T/A]GGGG | TAGGGGCC | 60.69 |
| +266 | Motif 1 - CTAGAGGT | CTACTGGC | 60.99 |
| +271 | Motif 2 - [T/G]G[T/A]GGGG | GGCTGGTC | 64.52 |
| +283 | Motif 3 - TCTCCCAA | TGACCCTC | 62.90 |
| +287 | Motif 3 - TCTCCCAA | CCTCTCTG | 69.14 |
| +293 | Motif 2 - [T/G]G[T/A]GGGG | TGCAGGAT | 61.64 |
| +300 | Motif 2 - [T/G]G[T/A]GGGG | TATTGGGC | 75.30 |
| +301 | Motif 2 - [T/G]G[T/A]GGGG | ATTGGGCT | 66.40 |
| +305 | Motif 2 - [T/G]G[T/A]GGGG | GGCTGGGC | 76.58 |
| +306 | Motif 2 - [T/G]G[T/A]GGGG | GCTGGGCG | 70.15 |
| +308 | Motif 2 - [T/G]G[T/A]GGGG | TGGGCGCT | 60.69 |
| +313 | Motif 2 - [T/G]G[T/A]GGGG | GCTGTGGC | 63.28 |
| +315 | Motif 2 - [T/G]G[T/A]GGGG | TGTGGCTG | 71.20 |
| +325 | Motif 2 - [T/G]G[T/A]GGGG | TGTGTGGG | 83.25 |

| | | | |
|---|---|---|---|
| +327 | Motif 2 - [T/G]G[T/A]GGGG | TGTGGGTA | 88.10 |
| +345 | Motif 2 - [T/G]G[T/A]GGGG | CGTGGGTG | 78.23 |
| +347 | Motif 2 - [T/G]G[T/A]GGGG | TGGGTGGT | 70.71 |
| +348 | Motif 2 - [T/G]G[T/A]GGGG | GGGTGGTG | 64.52 |
| +349 | Motif 2 - [T/G]G[T/A]GGGG | GGTGGTGA | 80.18 |
| +351 | Motif 2 - [T/G]G[T/A]GGGG | TGGTGAGA | 62.75 |
| +352 | Motif 2 - [T/G]G[T/A]GGGG | GGTGAGAG | 69.35 |
| +354 | Motif 1 - CTAGAGGT | TGAGAGCT | 78.48 |
| +354 | Motif 2 - [T/G]G[T/A]GGGG | TGAGAGCT | 70.18 |
| +359 | Motif 3 - TCTCCCAA | GCTGCCAG | 76.97 |
| +367 | Motif 1 - CTAGAGGT | CACGAGTT | 69.74 |
| +374 | Motif 2 - [T/G]G[T/A]GGGG | TCCGGGGG | 70.71 |
| +375 | Motif 2 - [T/G]G[T/A]GGGG | CCGGGGGG | 60.84 |
| +376 | Motif 2 - [T/G]G[T/A]GGGG | CGGGGGGA | 77.74 |
| +377 | Motif 2 - [T/G]G[T/A]GGGG | GGGGGGAC | 73.71 |
| +387 | Motif 2 - [T/G]G[T/A]GGGG | GGTTGTCT | 62.20 |
| +392 | Motif 3 - TCTCCCAA | TCTGCCCC | 73.67 |
| +395 | Motif 3 - TCTCCCAA | GCCCCCTG | 69.41 |
| +399 | Motif 3 - TCTCCCAA | CCTGCCCC | 67.86 |
| +401 | Motif 3 - TCTCCCAA | TGCCCCCA | 64.49 |
| +402 | Motif 3 - TCTCCCAA | GCCCCCAT | 75.65 |
| +407 | Motif 3 - TCTCCCAA | CATCCCTG | 72.88 |
| +417 | Motif 2 - [T/G]G[T/A]GGGG | GCTTGGCC | 62.20 |
| +426 | Motif 2 - [T/G]G[T/A]GGGG | GGATGGTG | 74.01 |
| +427 | Motif 2 - [T/G]G[T/A]GGGG | GATGGTGC | 63.28 |
| +432 | Motif 3 - TCTCCCAA | TGCCCCAT | 68.71 |
| +440 | Motif 2 - [T/G]G[T/A]GGGG | TGCAGGTA | 60.41 |
| +444 | Motif 2 - [T/G]G[T/A]GGGG | GGTAGGCA | 71.91 |

| | | | |
|---|---|---|---|
| +450 | Motif 3 - TCTCCCAA | CAGCCCAG | 66.96 |
| +456 | Motif 2 - [T/G]G[T/A]GGGG | AGCTGGAC | 62.00 |
| +463 | Motif 3 - TCTCCCAA | CCTCCCTG | 86.32 |
| +467 | Motif 2 - [T/G]G[T/A]GGGG | CCTGGGAA | 62.55 |
| +481 | Motif 1 - CTAGAGGT | GCAGAGGG | 78.78 |
| +481 | Motif 2 - [T/G]G[T/A]GGGG | GCAGAGGG | 60.23 |
| +483 | Motif 2 - [T/G]G[T/A]GGGG | AGAGGGCC | 80.25 |
| +490 | Motif 1 - CTAGAGGT | CTACAGGC | 76.59 |
| +495 | Motif 2 - [T/G]G[T/A]GGGG | GGCTGGGC | 76.58 |
| +496 | Motif 2 - [T/G]G[T/A]GGGG | GCTGGGCC | 70.15 |
| +502 | Motif 1 - CTAGAGGT | CCTGAGTT | 67.52 |
| +504 | Motif 2 - [T/G]G[T/A]GGGG | TGAGTTGC | 63.30 |
| +507 | Motif 3 - TCTCCCAA | GTTGCCAC | 64.83 |
| +514 | Motif 3 - TCTCCCAA | CCTGCCCC | 67.86 |
| +516 | Motif 3 - TCTCCCAA | TGCCCCCA | 64.49 |
| +517 | Motif 3 - TCTCCCAA | GCCCCCAG | 78.83 |
| +519 | Motif 1 - CTAGAGGT | CCCCAGGT | 64.15 |
| +526 | Motif 3 - TCTCCCAA | TCTGCGTA | 61.48 |
| +531 | Motif 1 - CTAGAGGT | GTAGATGG | 65.22 |
| +533 | Motif 2 - [T/G]G[T/A]GGGG | AGATGGTG | 70.26 |
| +534 | Motif 2 - [T/G]G[T/A]GGGG | GATGGTGA | 63.28 |
| +550 | Motif 2 - [T/G]G[T/A]GGGG | TCCTGGGT | 62.75 |
| +551 | Motif 2 - [T/G]G[T/A]GGGG | CCTGGGTA | 61.33 |
| +555 | Motif 2 - [T/G]G[T/A]GGGG | GGTAGAGT | 65.03 |
| +556 | Motif 1 - CTAGAGGT | GTAGAGTG | 68.59 |
| +558 | Motif 1 - CTAGAGGT | AGAGTGGT | 74.65 |
| +558 | Motif 2 - [T/G]G[T/A]GGGG | AGAGTGGT | 73.38 |
| +560 | Motif 2 - [T/G]G[T/A]GGGG | AGTGGTGC | 76.43 |
| +562 | Motif 2 - [T/G]G[T/A]GGGG | TGGTGCGG | 62.75 |
| +563 | Motif 2 - [T/G]G[T/A]GGGG | GGTGCGGC | 80.18 |

| | | | |
|---|---|---|---|
| +565 | Motif 2 - [T/G]G[T/A]GGGG | TGCGGCCA | 60.69 |
| +566 | Motif 3 - TCTCCCAA | GCGGCCAG | 61.64 |
| +570 | Motif 1 - CTAGAGGT | CCAGGGCC | 65.60 |
| +570 | Motif 2 - [T/G]G[T/A]GGGG | CCAGGGCC | 60.31 |
| +576 | Motif 1 - CTAGAGGT | CCAGATGG | 71.33 |
| +578 | Motif 2 - [T/G]G[T/A]GGGG | AGATGGGG | 82.32 |
| +579 | Motif 2 - [T/G]G[T/A]GGGG | GATGGGGT | 80.18 |
| +581 | Motif 2 - [T/G]G[T/A]GGGG | TGGGGTCC | 60.69 |
| +584 | Motif 3 - TCTCCCAA | GGTCCCAC | 80.25 |
| +586 | Motif 3 - TCTCCCAA | TCCCACAG | 69.03 |
| +590 | Motif 1 - CTAGAGGT | ACAGAGCA | 72.40 |
| +599 | Motif 3 - TCTCCCAA | GCTCCTTG | 67.56 |
| +614 | Motif 3 - TCTCCCAA | GCTGCCTT | 64.38 |
| +630 | Motif 1 - CTAGAGGT | CTTGTGGC | 62.30 |
| +632 | Motif 2 - [T/G]G[T/A]GGGG | TGTGGCTG | 71.20 |
| +665 | Motif 1 - CTAGAGGT | CTACTGGT | 67.50 |
| +668 | Motif 1 - CTAGAGGT | CTGGTGGC | 62.30 |
| +669 | Motif 2 - [T/G]G[T/A]GGGG | TGGTGGCG | 69.63 |
| +670 | Motif 2 - [T/G]G[T/A]GGGG | GGTGGCGG | 80.18 |
| +672 | Motif 2 - [T/G]G[T/A]GGGG | TGGCGGAG | 61.64 |
| +673 | Motif 2 - [T/G]G[T/A]GGGG | GGCGGAGG | 67.64 |
| +674 | Motif 1 - CTAGAGGT | GCGGAGGA | 63.13 |
| +676 | Motif 2 - [T/G]G[T/A]GGGG | GGAGGAAG | 66.30 |
| +690 | Motif 2 - [T/G]G[T/A]GGGG | GGTGAGTT | 68.12 |

### ESS decamers from Wang et al.

Wang Z, Rolish ME, Yeo G, Tung V, Mawson M, Burge CB. Systematic identification and analysis of exonic splicing silencers. Cell. 2004 Dec 17;119(6):831-45

| Sequence Position | cDNA Position | Silencer motif |
|---|---|---|
| 210 | +210 | GGTAAGTGTA |

### Predicted PESS Octamers from Zhang & Chasin.

Zhang XH, Chasin LA. Computational definition of sequence motifs governing constitutive exon splicing. Genes Dev. 2004 Jun 1;18(11):1241-50

| Sequence Position | cDNA Position | Silencer motif | Motif value (0-100) |
|---|---|---|---|
| 68 | +68 | GATGGGTG | 26.14 |
| 97 | +97 | GGGGATGG | 33.2 |
| 375 | +375 | CCGGGGGG | 2.24 |
| 493 | +493 | CAGGCTGG | 3.81 |
| 533 | +533 | AGATGGTG | 32.13 |
